# Methane emissions from upland tree stems reflect a balance of microbial production and consumption

**DOI:** 10.1101/2025.09.30.679632

**Authors:** Jonathan Gewirtzman, Wyatt Arnold, Meghan Taylor, Hannah Burrows, Carter Merenstein, David J. Woodbury, Naomi Whitlock, Kendall Kraut, Leslie Gonzalez, Craig R. Brodersen, Marlyse C. Duguid, Peter A. Raymond, Jordan Peccia, Mark A. Bradford

**Affiliations:** Yale School of the Environment, Yale University, New Haven, CT, USA; Department of Chemical and Environmental Engineering, School of Engineering and Applied Science, Yale University, New Haven, CT, USA; WCRP Climate and the Cryosphere, University of Massachusetts, Amherst, Amherst, MA, USA; Harvard College, Harvard University, Cambridge, MA, USA; Department of Astronomy and Astrophysics, University of Chicago, Chicago, IL, USA; Department of Biostatistics, Harvard T.H. Chan School of Public Health, Boston, MA, USA; Division of Infectious Disease, Boston Children’s Hospital, Boston, MA, USA; New Mexico Highlands University, Department of Forestry, Las Vegas, NM, USA; Wesleyan University, Middletown, CT, USA; Yale College, Yale University, New Haven, CT, USA; The Forest School, Yale School of the Environment, Yale University, New Haven, CT, USA; Yale Center for Natural Carbon Capture, Yale University, New Haven, CT, USA

**Keywords:** forest carbon cycle, heartwood, methane emissions, methanogens, methanotrophs, temperate forest, tree microbiome, tree species

## Abstract

**Rationale:** Upland trees emit methane (CH₄), but whether it is produced inside the wood or transported from soil remains debated. Methanogens occur in the heartwood of several species, yet their relation to co-occurring methanotrophs is poorly understood.

**Methods:** We measured 1,200 stem and 288 soil fluxes, sampled internal stem gases including δ¹³CH₄, quantified methanogens and methanotrophs via droplet digital PCR (ddPCR) in 592 samples, sequenced 16S rRNA genes, and bounded stand-scale fluxes.

**Key Results:** Methanogens were detected in 97% of heartwood samples, about tenfold above soil; methanotroph genes (*pmoA*, *mmoX*) were near-ubiquitous. Wood harbored distinct methanogen communities dominated by hydrogenotrophic Methanobacteriaceae, consistent with depleted δ¹³CH₄. Fluxes were roughly uniform with height for most species, consistent with internal production, declining steeply only in wet microsites (*Betula alleghaniensis*). Across ten tree species, the methanogen:methanotroph ratio tracked emissions (R² = 0.46). Stems below 2 m offset about 1% of soil CH₄ uptake, the whole woody surface 2.4–19%; the stand remained a net sink.

**Main Conclusion:** Internal methanogenesis contributes widely to upland tree methane emissions, and across tree species net flux tracked the methanogen:methanotroph balance. The stand-scale contribution depends most on how stem flux changes above 2 m, then on woody surface area.

## Introduction

Forests are key regulators of the global methane (CH₄) cycle, functioning as both sinks and sources of this potent greenhouse gas. Methane has a global warming potential more than 80 times that of carbon dioxide (CO₂) over 20 years (Forster *et al*., 2021; Wood *et al*., 2023). While the role of soil in forest methane cycling has been extensively studied, trees also take part in the CH₄ cycle through several concurrent pathways, including internal microbial methanogenesis, transport of soil-produced CH₄, and oxidation by methanotrophic bacteria colonizing bark and woody surfaces (Barba *et al*., 2019a; Covey & Megonigal, 2019; Gauci *et al*., 2024).

The source of tree methane differs between wetland and upland forests. In wetland forests, trees primarily act as conduits for soil-derived methane transported through vascular pathways (Pangala *et al*., 2013, 2015, 2017; Pitz *et al*., 2018; Jeffrey *et al*., 2019; Jeffrey *et al*., 2023). Upland forests, where well-drained soils are traditionally methane sinks, also have trees that emit methane (Pitz & Megonigal, 2017; Warner *et al*., 2017; Covey & Megonigal, 2019; Epron *et al*., 2022; Machacova *et al*., 2023; Bréchet *et al*., 2025; Hettwer *et al*., 2025), raising the possibility that the methane is produced within the trees. Recent detections of endophytic methanogens support this hypothesis (Yip *et al*., 2019; Feng *et al*., 2022; Harada *et al*., 2024), but factors governing production remain poorly understood.

Methanogens (archaea that produce methane by hydrogenotrophic, acetoclastic or methylotrophic pathways, generally under anoxic conditions) (Le Mer & Roger, 2001) have been detected in heartwood of multiple species (Zeikus & Ward, 1974; Zeikus & Henning, 1975; Covey *et al*., 2012; Yip *et al*., 2019; Flanagan *et al*., 2021; Feng *et al*., 2022; Harada *et al*., 2024; Mochidome *et al*., 2025; Moisan *et al*., 2025). Anaerobic conditions in heartwood (Jensen, 1967; Shigo & Hillis, 1973; Schink & Ward, 1984; Arnold *et al*., 2025), where oxygen diffusion is restricted, may create suitable methanogenic niches (Uroz *et al*., 2016; Mieszkin *et al*., 2021). Yet how widely methanogens colonize wood, which pathways they use, and how much they add to forest methane budgets remain unclear (Barba *et al*., 2019a, 2024; Putkinen *et al*., 2021).

Methanotrophs oxidize CH₄ to CO₂ generally under aerobic conditions. Methanotrophs dwelling in tree outer tissues may intercept a substantial fraction of internally produced or transported CH₄ before atmospheric release (Jeffrey *et al*., 2021a,b; Carmichael *et al*., 2024; Leung *et al*., 2026), and on some upland trees woody-surface CH₄ uptake has been proposed to exceed emission, particularly higher on the stem (Gauci *et al*., 2024). However, net uptake has been reported from few sites. Because methane producers and consumers co-occur, net fluxes at tree and ecosystem scales can obscure the underlying gross processes, and quantitative data on methanogen and methanotroph abundance, activity, and spatial distribution remain sparse (Putkinen *et al*., 2021). Whether methanotroph activity varies systematically with tree species, tissue type, or environmental conditions is largely unknown. Given that global woody surface area may approximate total land surface area (Gauci *et al*., 2024; Gewirtzman, 2026), even small per-area net fluxes, whether sources from internal methanogenesis or sinks from bark methanotrophy, could matter for the global CH₄ budget.

Here we ask whether the methane emitted by upland tree stems is produced inside the wood, and whether co-occurring methanotrophs also shape the net flux. Across a moisture gradient from upland to wetland margin in a mixed temperate forest in the northeastern USA, we (1) quantified stem methane flux, using vertical profiles to distinguish soil-derived transport from internal production; (2) characterized methanogenic and methanotrophic communities across tree species and tissues; (3) related gene abundances to flux at the individual and species levels, and to species traits such as wood and bark density, soil-moisture niche and longevity; and (4) bounded the stand-scale contribution of stems relative to soil uptake.

## Materials and Methods

### Study site and design

Fieldwork was at Yale-Myers Forest, a 3,213-ha second-growth mixed-deciduous research forest in northeastern Connecticut, USA (41°56′N, 72°07′W; 170–300 m a.s.l.; mean annual temperature 9.5°C; mean annual precipitation 123 cm), on moderately to well-drained stony loams derived from glacial till. Flux measurements were made within and immediately adjacent to a 4-ha (200 × 200 m) permanent inventory plot (Fig. S1); destructive sampling for microbiology took place immediately adjacent to the plot, because it is not permitted within it. The data come from four campaigns and the plot’s 2018–2019 stem census (Table 1); procedural detail for every step below is given in Methods S1. The monthly survey covered six plots, two in each of three moisture classes along the gradient: upland, intermediate, and wetland margin (the wettest, at the edge of a wetland). All data and code are openly archived (Data Availability).

**Table 1.**
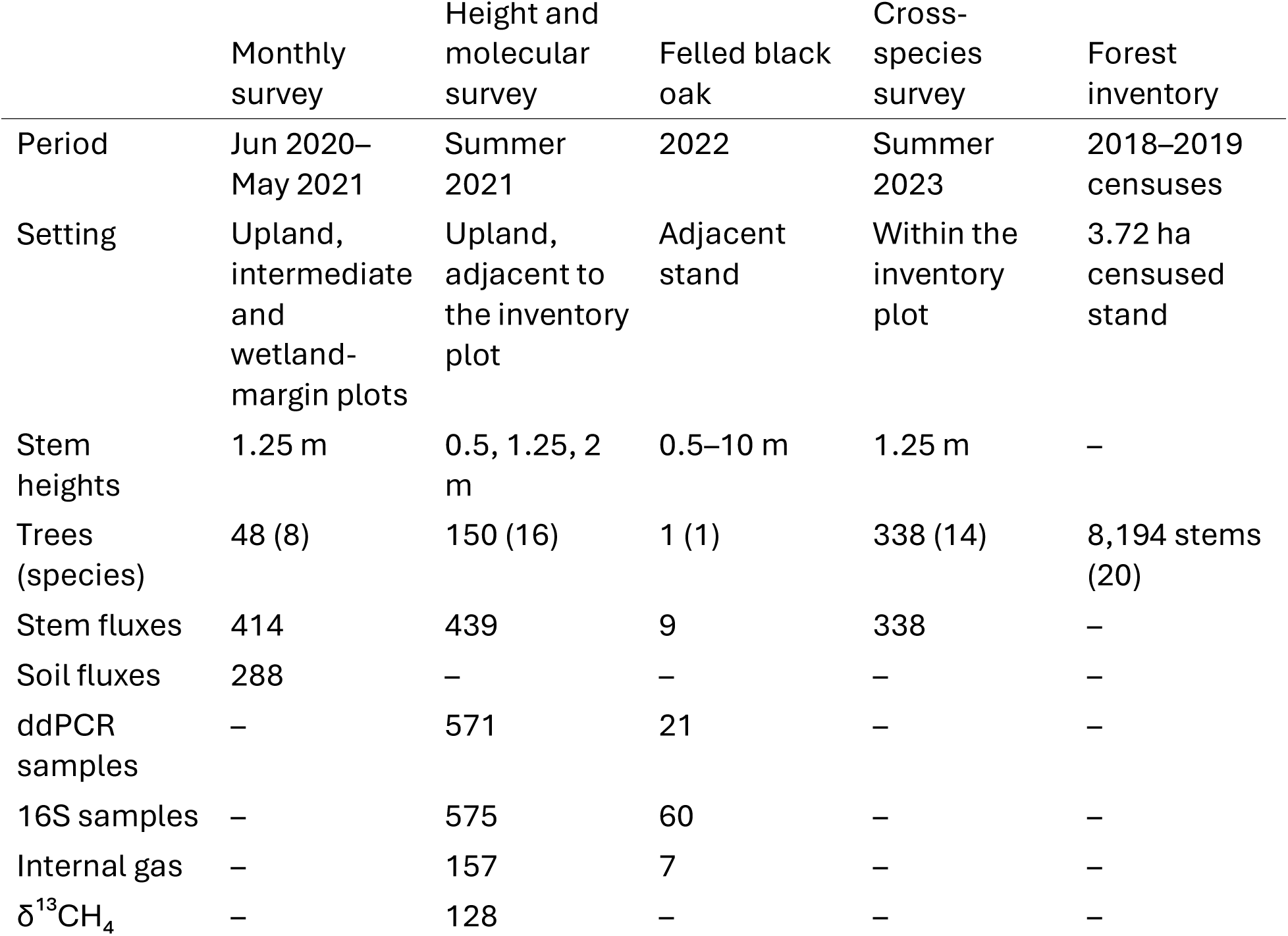
Field campaigns and sample sizes. Trees are those with flux measurements; 33 trees of the monthly survey were measured again in 2023, so the campaigns cover 503 trees. In the 2021 survey, 155 trees were cored, 12 of them without flux measurements; internal gas was sampled in 157 trees, including three hemlocks that were not cored. Stem and soil fluxes are deployments with a CH₄ value.

### Flux measurements

Stem fluxes were measured with semi-rigid chambers (Siegenthaler *et al*., 2016) in 2020–2021 and rigid chambers in 2021–2023, and soil fluxes with static chambers on PVC collars (Fig. S2; Methods S1). Chambers were connected to a portable off-axis integrated cavity output spectroscopy analyzer (GLA131-GGA, Los Gatos Research) for 3–16 min, excluding the first 30 s. Trees, species, heights and periods for each campaign are in Table 1. Soil temperature (10 cm) and volumetric water content (VWC, 12 cm) were recorded at each measurement, and VWC was mapped across the plot in December 2020.

Fluxes were calculated with goFlux v0.5.0.9003 (https://doi.org/10.5281/zenodo.23383778), a fork of goFlux (Rheault *et al*., 2024), whose best.flux() criteria chose between linear and Hutchinson–Mosier fits (Hutchinson & Mosier, 1981; Hüppi *et al*., 2018) (Methods S1, S2). Each flux was classed as detected or not detected against its own minimum detectable flux. Small, negative and undetected fluxes were all kept, because discarding or replacing values below detection biases the data (Parkin *et al*., 2012; de Klein *et al*., 2020; Maier *et al*., 2022) (Methods S2). Only two soil deployments were excluded, because the chamber was not sampling ambient air at closure (Methods S2).

### Sample collection and DNA extraction

Cores were taken at 1.25 m with flame-sterilized 5-mm increment borers and frozen on dry ice. Each core was cut into an outer and an inner 5-cm segment, operationally defined as sapwood and heartwood. Segments were lyophilized and cryo-ground (Arnold *et al*., 2024). Organic and mineral soil (to 30 cm) was composited from four points around each tree. DNA was extracted from 100 mg wood or 250 mg soil (Methods S1).

### Gene abundance

Genes for methyl-coenzyme M reductase (*mcrA*) and particulate and soluble methane monooxygenase (*pmoA*, *mmoX*) were quantified by droplet digital PCR (ddPCR; Bio-Rad QX200): *mcrA* with a custom probe-based assay (Arnold *et al*., 2024), which reduces false positives relative to the standard assay (Fig. S3), and *pmoA* and *mmoX* with EvaGreen assays (Fuse *et al*., 1998; Bourne *et al*., 2001) (Table S1; full protocol in Notes S1). Gene abundances are expressed per gram of dry wood or fresh soil (dry-soil equivalents in Methods S1). The probe assay’s limit of detection was about 360 gene copies in a 100 mg wood sample and 480 in a 250 mg soil sample (Notes S1). Copies per gram were calculated from each reaction’s concentration, the template, eluate and lysate volumes and the sample mass (Methods S1; Notes S1). Bacterial 16S rRNA and fungal internal transcribed spacer (ITS) copy numbers in each extract were quantified by quantitative PCR (qPCR) at the sequencing facility. Means of gene abundances are means of log₁₀ values.

### 16S sequencing and community analyses

The 16S rRNA V4 region was sequenced (515F/806R with peptide nucleic acid blockers; Illumina MiSeq 2 × 300 bp) and processed into amplicon sequence variants (ASVs) with DADA2 (Callahan *et al*., 2016) via Nephele (Weber *et al*., 2018) against SILVA v138.1 (Quast *et al*., 2013), removing chloroplast and mitochondrial reads and rarefying to 3,500 reads (Methods S1); these sequences were also reported by Arnold *et al*. (2025). Functional potential was inferred with PICRUSt2 (Douglas *et al*., 2020) and FAPROTAX (Louca *et al*., 2016), and community analyses used phyloseq and microeco (McMurdie & Holmes, 2013; Liu *et al*., 2021). Methanogens were ASVs in methanogenic archaeal families (Table S2).

Methanotrophs were defined by genetic capacity for methane monooxygenase, particulate (pMMO) or soluble (sMMO) (Knief, 2015), among ASVs in families that contain methanotrophs (Table S2). ASVs in an established methanotroph genus, or in a family whose characterized members are all methanotrophs (including ASVs unresolved at genus level), were counted as Known methanotrophs. ASVs in mixed families (Beijerinckiaceae, Methylocystaceae, Methylacidiphilaceae) that were unresolved at genus level, and ASVs from genera with no sequenced genome or with the enzyme in only some genomes, were counted as Putative methanotrophs, an upper bound. Genus capacity was screened in NCBI protein records against reference PmoA and MmoX proteins (Table S3). The ASV tree was used to estimate how much of the Putative tier lies within about 97% identity of a methanotroph genus, and methanotrophs were also summarized at order and family level (Methods S3; Table S4). Agreement between each 16S group and the corresponding ddPCR gene was assessed in the same samples, by detection and by correlation (Fig. S4; Table S5).

### Internal gas and stable isotopes

Internal gas was sampled from increment-borer holes drilled bark to pith, sealed and equilibrated for 5 min, and analyzed for CH₄, CO₂, N₂O and O₂ by gas chromatography. δ¹³CH₄ and δ¹³CO₂ were measured on a Picarro G2201-i analyzer. The tree-produced source was estimated as the intercept of δ¹³CH₄ against 1/[CH₄] (Keeling plot; Keeling, 1958), fitted by MM robust regression (Yohai, 1987; Venables & Ripley, 2002), and by two-endmember mixing with atmospheric CH₄ (1.9 ppm, −47‰); sensitivity checks and the apparent CO₂–CH₄ fractionation factor are in Methods S1.

### Whole-tree profiling of a felled black oak

One mature black oak (*Ǫuercus velutina*), the individual described by Arnold *et al*. (2025), was profiled while standing: stem flux and internal gas were measured at 0.5, 1.25, 2, 4, 6, 8 and 10 m, with heights above 2 m reached by an arborist climber. It was then felled and sampled destructively at each height (Methods S1), and is analyzed separately from the cross-species and upscaling analyses.

### Data analysis

Height effects were tested for each species with at least three trees by linear mixed-effects models of flux on height, with tree as a random intercept (lme4, Bates *et al*., 2015). Variance in stem flux among trees was partitioned using one value per tree, the mean of its growing-season (May–September) breast-height measurements on an inverse hyperbolic sine (arcsinh) scale. Predictors were environment (diameter, air and soil temperature, soil VWC), species and their interactions, with chamber design as a covariate. Bark-loss scores (2023; Methods S2) and ITS load were related to flux and *mcrA* by linear and quadratic mixed models, compared by AIC. Tree-level gene abundances were area-weighted, averaged over the breast-height cross-section (Methods S4), and the methanogen:methanotroph ratio (hereafter, the ratio) was *mcrA*:(*pmoA* + *mmoX*). Gene–flux relationships were tested between trees, with linear models that included species, and between species, on species medians (Methods S4). Per-species plant traits (wood and bark density, wood and bark chemistry and decay, root and whole-plant traits from the Global Wood Density Database and the TRY Plant Trait Database (Kattge *et al*., 2020; Fischer *et al*., 2026)) and each species’ realized soil-moisture niche were related to species-level gene abundances and flux by Spearman correlation across the ten species with at least five trees and across the eight broadleaf species among them, with Benjamini–Hochberg false discovery rate (FDR) correction (Benjamini & Hochberg, 1995) (Methods S5).

### Stand-scale bounds

Stem and soil fluxes were predicted across the inventory with random forests (ranger, Wright & Ziegler, 2017), trained on live stems only (1,126 measurements from 488 trees) because the inventory contains no dead stems; skill was assessed with whole trees or collars held out. Predictions used monthly soil-temperature and soil-moisture climatologies and a per-species calibration (Methods S6). The measured component of the stem budget is the lateral surface of every inventoried stem between 0 and 2 m. Above 2 m, we combined every value of four unconstrained assumptions (woody area index, bole shape, vertical placement of branch area, and six forms of flux change above 2 m) in a grid of 240 combinations, and report the measured band and the extrapolation separately (Methods S6).

## Results

### Tree and soil methane fluxes from upland to wetland-margin plots

In upland plots (Fig. 1), tree stems showed predominantly positive CH₄ fluxes (mean 0.049 nmol m⁻² s⁻¹; 70% positive; p = 0.008, one-sided one-sample t-test), whereas upland soils took up CH₄ (mean −1.57 nmol m⁻² s⁻¹). In intermediate plots, tree fluxes remained predominantly positive but lower (mean 0.014 nmol m⁻² s⁻¹), and soils continued to take up CH₄ (mean −1.34 nmol m⁻² s⁻¹). In wetland-margin plots, trees had the highest emissions (mean 0.109 nmol m⁻² s⁻¹; 78% positive), and soil fluxes were variable: 59% of measurements were negative (median −0.076 nmol m⁻² s⁻¹), but occasional large emissions raised the mean to 1.40 nmol m⁻² s⁻¹.

**Figure 1.**
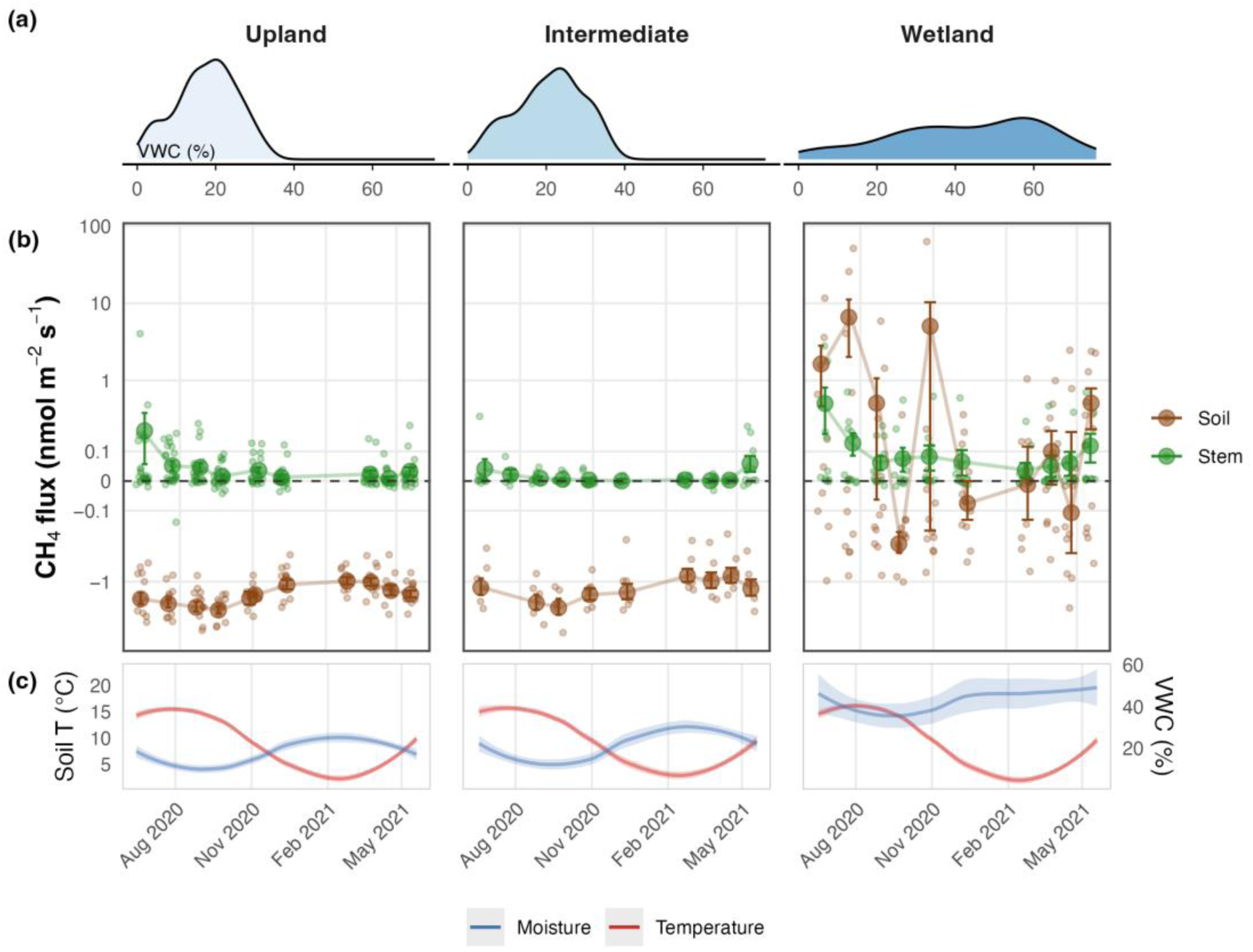
Soil moisture regimes and temporal patterns of CH₄ flux across a moisture gradient. (a) Kernel density distributions of volumetric water content (VWC, %) in the three plot classes: Upland, Intermediate, and Wetland (wetland margin). (b) Tree stem (green) and soil (brown) CH₄ flux (nmol m⁻² s⁻¹) measured across the three plot classes during 2020–2021. Small points are individual measurements; large points and error bars show mean ± SE for each sampling campaign (measurements within 7 days grouped). Arcsinh y-axis. (c) Loess-smoothed soil temperature (°C; red) and volumetric water content (blue) at each site.

### Methane flux by height and analysis of height effects

Methane flux measurements at three stem heights (0.5, 1.25, 2.0 m) showed predominantly positive fluxes for all upland species (Fig. 2a). Linear mixed-effects models revealed significant height effects in only 2 of 11 species tested (Fig. 2b). *Betula alleghaniensis* showed the strongest negative relationship (−0.797 nmol m⁻² s⁻¹ per m of stem height, p = 0.002), with fluxes decreasing from 1.47 ± 0.51 nmol m⁻² s⁻¹ at 0.5 m to 0.28 ± 0.08 nmol m⁻² s⁻¹ at 2.0 m. This effect occurred where soil moisture (35.9% VWC) and soil methanogens (10⁶·⁴ *mcrA* copies g⁻¹) were highest. *Tsuga canadensis* showed a much weaker negative effect (−0.019 nmol m⁻² s⁻¹ per m, p = 0.026). In leave-one-tree-out jackknife tests, the height effect stayed significant in every iteration for *B. alleghaniensis* and in 82% for *T. canadensis*. *Acer saccharum* showed a positive trend (+0.196, p = 0.050) driven largely by one tree (Fig. 2b).

**Figure 2.**
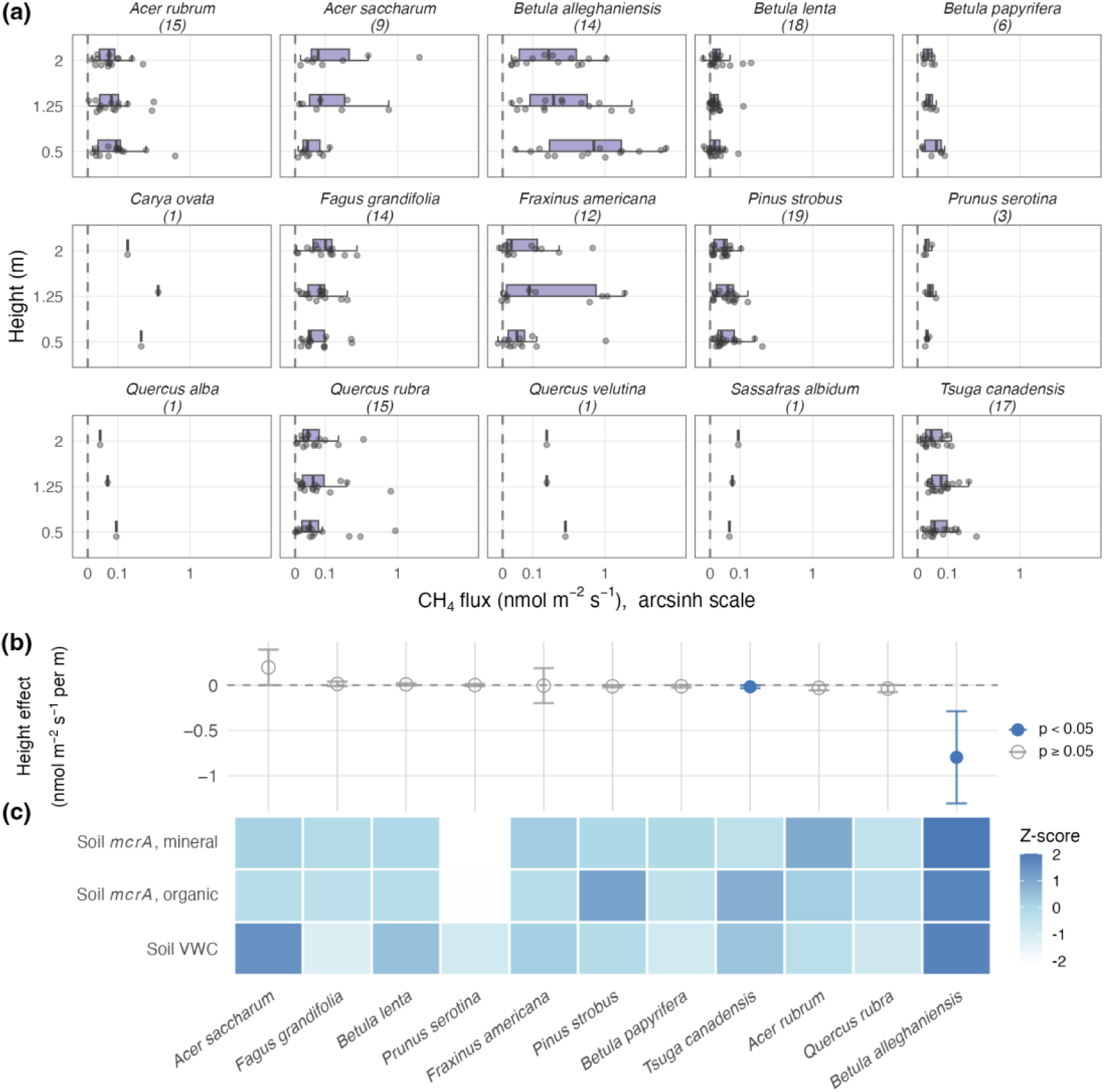
Height-dependent CH₄ flux patterns across tree species. (a) Species-level CH₄ flux (nmol m⁻² s⁻¹) measured at 0.5, 1.25 and 2 m stem height (n, trees), on a shared arcsinh axis with the same breaks in every panel. Points are individual measurements with half-boxplots; dashed line, zero flux (left, net uptake; right, net emission). (b) Linear mixed-effects model coefficients for the height effect on CH₄ flux (nmol m⁻² s⁻¹ per m), with 95% confidence intervals, for species with at least three trees measured. Filled points indicate significant effects (p < 0.05); open points are non-significant. Species ordered by coefficient; leave-one-tree-out jackknife results are given in the text (11 of 14 *B. alleghaniensis* trees and 71% of *T. canadensis* trees had negative slopes; the high-flux *A. saccharum* tree emitted 1.90 nmol m⁻² s⁻¹ at 2 m). (c) Z-score heatmap of log₁₀ soil *mcrA* abundance (mineral, organic) and soil VWC by species, ordered to match panel (b).

Across the 11 species, height coefficients were negatively correlated with soil methanogen abundance (r = −0.64, p = 0.035; Fig. 2c). The correlation with soil moisture alone was weaker (r = −0.41, p = 0.21; r = −0.61, p = 0.047 using species medians of per-tree slopes, Fig. S5), though soil moisture and methanogen abundance were positively correlated (r = 0.46, p < 0.001). Across individual trees, the height slope steepened with soil moisture (r = −0.32, p = 0.0001, 138 trees; Fig. S5a).

### Species trends in flux and individual variance partitioning

Averaging growing-season (May–September) breast-height (1.25 m) measurements to one value per tree (480 trees; 691 measurements from the 2020–2021, 2021 and 2023 surveys) showed that flux varied widely within and among species (Fig. 3a). Species occupied different soil-moisture niches within the plot (Fig. S6), and sampled diameters by species and campaign are given in Table S6. Variance partitioning attributed 14.6% of among-tree variance to species identity, 12.3% to species × environment interactions and 1.9% to environment alone; 3.2% was shared between species and environment, and 68.1% was unexplained (Fig. 3b; effect sizes in Fig. 3c).

**Figure 3.**
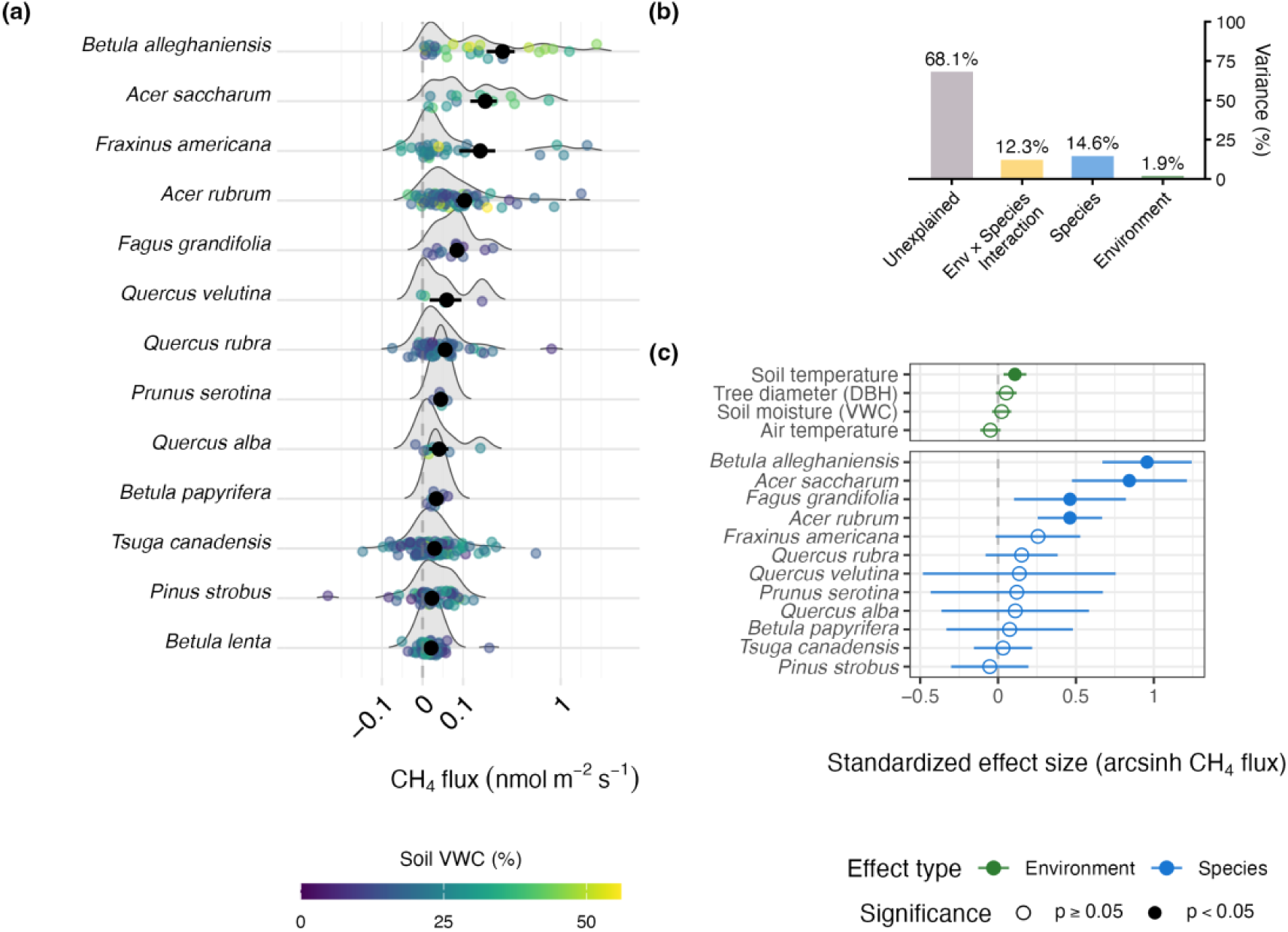
Species-level CH₄ flux distributions and variance partitioning. (a) Ridgeline density plots of per-tree mean CH₄ flux (growing-season, breast-height measurements; arcsinh axis) by species, ordered by mean flux. Points are trees, colored by soil volumetric water content (viridis scale); black points with error bars show species mean ± SE across trees. (b) Partitioning of among-tree variance in arcsinh-transformed flux into species identity, species × environment interaction, environment, and unexplained components; the remaining 3.2% is shared between species and environment and cannot be assigned to either. (c) Standardized effect sizes from an additive linear model fitted to tree means, with chamber design as a covariate, for environmental (green) and species (blue) predictors. Filled points indicate p < 0.05; open points are non-significant.

*Betula* species had both the highest (*B. alleghaniensis*, yellow birch, 0.276 ± 0.087 nmol m⁻² s⁻¹; mean ± SE across 29 trees) and lowest (*B. lenta*, black birch, 0.018 ± 0.004; 62 trees) mean emissions (Fig. 3a). Next highest were *A. saccharum* (sugar maple, 0.182 ± 0.059; 13 trees) and *Fraxinus americana* (white ash, 0.160 ± 0.072; 31 trees); the gymnosperms *T. canadensis* (eastern hemlock, 0.026 ± 0.007; 124 trees) and *Pinus strobus* (white pine, 0.020 ± 0.010; 53 trees) were also low (natural-history context in Methods S4).

### Quantifying methanogens and methanotrophs across tree and soil compartments

In the 2021 survey of 155 trees of 16 species, 97.3% of heartwood samples contained detectable *mcrA* (Fig. 4a; Fig. S7). Among positive samples, mean abundance was 10⁴·⁷ copies g⁻¹ dry wood (median 10⁴·²), with maximum values reaching 10⁸·⁴ copies g⁻¹ in *A. saccharum* (sugar maple). Ninety-two percent of heartwood samples were at or above the assay’s limit of detection (10³·⁶ copies g⁻¹ dry wood; Notes S1). Sapwood showed significantly lower methanogen abundance (Wilcoxon rank-sum test, p < 0.001), with 69% of samples detectable and 40% at or above the limit, averaging 10³·⁸ copies g⁻¹ (median 10³·⁶, max 10⁵·⁷) among positive samples.

**Figure 4.**
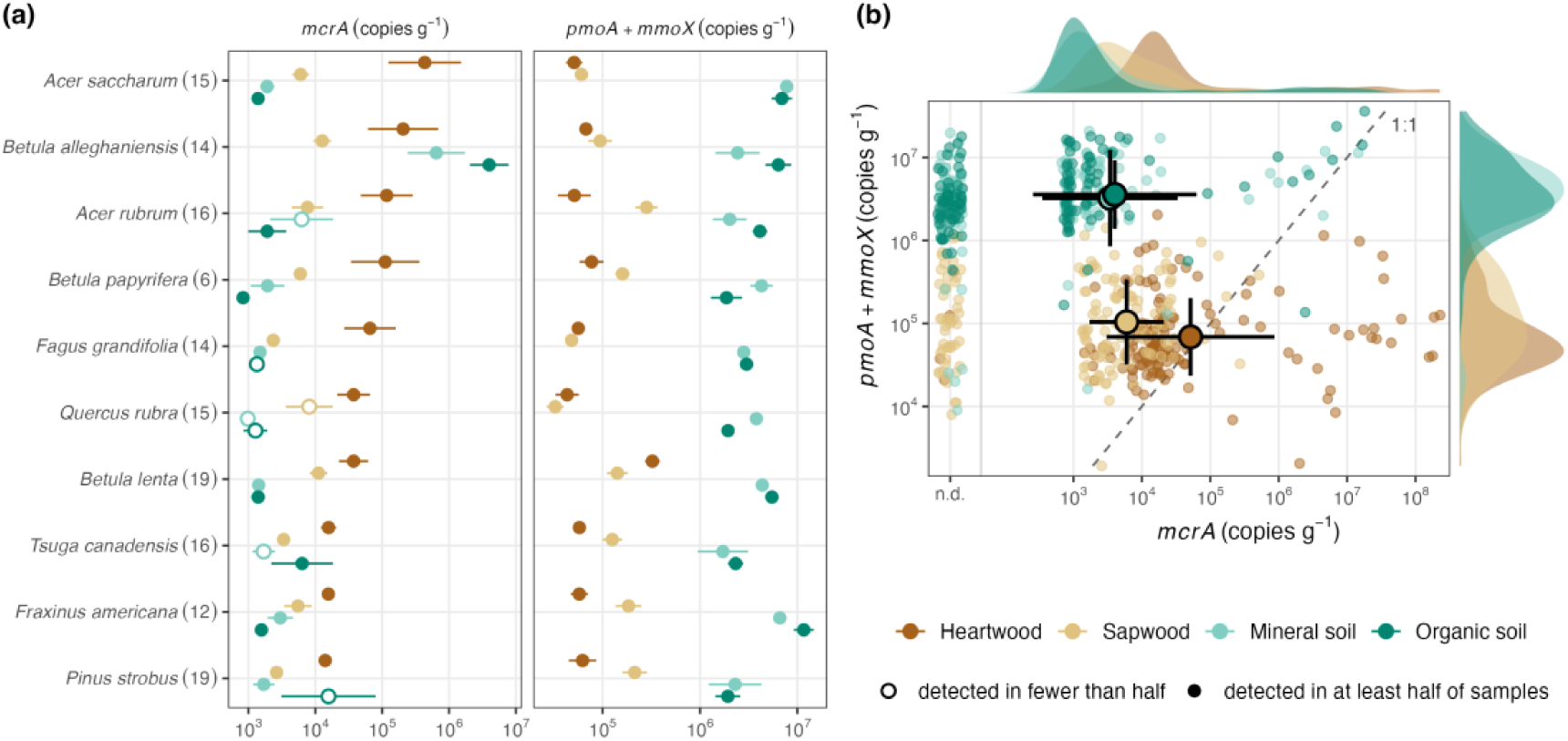
Methanogen and methanotroph gene abundance by species and compartment. (a) *mcrA* and *pmoA* + *mmoX* (copies g⁻¹; dry mass for wood, fresh mass for soil) in heartwood, sapwood, mineral soil and organic soil for the ten species in the species-level gene–flux analyses (at least five trees), ranked by heartwood *mcrA*; trees sampled in parentheses. Points, mean of log₁₀ among positive samples ± SE; hollow points, compartments where fewer than half of samples were positive. All 16 species, with *pmoA* and *mmoX* separately, are in Fig. S7. (b) *mcrA* against *pmoA* + *mmoX* for every sample, with marginal densities of positive samples; samples without detectable *mcrA* are plotted at n.d., and the one sample without detectable *pmoA* or *mmoX* is omitted. Large points and bars, compartment mean ± SD of log₁₀ among positive samples; dashed line, 1:1.

Methanogens were less abundant in soils surrounding trees. Only 59% of mineral and 53% of organic soil samples contained detectable methanogens, and 17–23% were at or above the limit of detection (10³·³ copies g⁻¹ fresh soil), against 92% of heartwood. Among positive samples, mineral soils averaged 10³·⁵ copies g⁻¹ fresh soil and organic horizons 10³·⁶ (both medians 10³·²), about an order of magnitude below heartwood. Exceptions occurred at the wettest positions, particularly around *B. alleghaniensis*, where soil methanogens reached 10⁷·² (mineral) and 10⁷·³ (organic) copies g⁻¹.

Methanotrophs showed the inverse pattern. Combined *pmoA* and *mmoX* abundances were lowest in heartwood (10⁴·⁸ copies g⁻¹), intermediate in sapwood (10⁵·⁰) and highest in soils (10⁶·⁵–10⁶·⁶), and both genes were detected in at least 93% of samples in every compartment. The two genes were present at similar levels in heartwood, whereas *pmoA* exceeded *mmoX* about threefold in sapwood and by about two orders of magnitude in soils; they covaried in soils but not in wood (Fig. S8).

### Community composition of methanogens and methanotrophs

Sequencing of 16S rRNA genes also found methanogens throughout the wood microbiome. In heartwood, methanogens made up 0–54.7% of the community (mean 3.2 ± 9.1%), and in sapwood 0–5.9% (mean 0.10 ± 0.59%; Fig. 5a). Methanogen abundance varied by tree species (ANOVA: 16S p < 0.01; ddPCR p = 0.01 among the ten species with at least five trees). *Acer saccharum* heartwood contained the highest methanogen concentrations both in absolute terms (Fig. 4a; mean log₁₀ *mcrA* 5.6 ± 2.1 copies g⁻¹, SD) and as a proportion of the total microbiome (15.8 ± 13.1%, p < 0.001).

**Figure 5.**
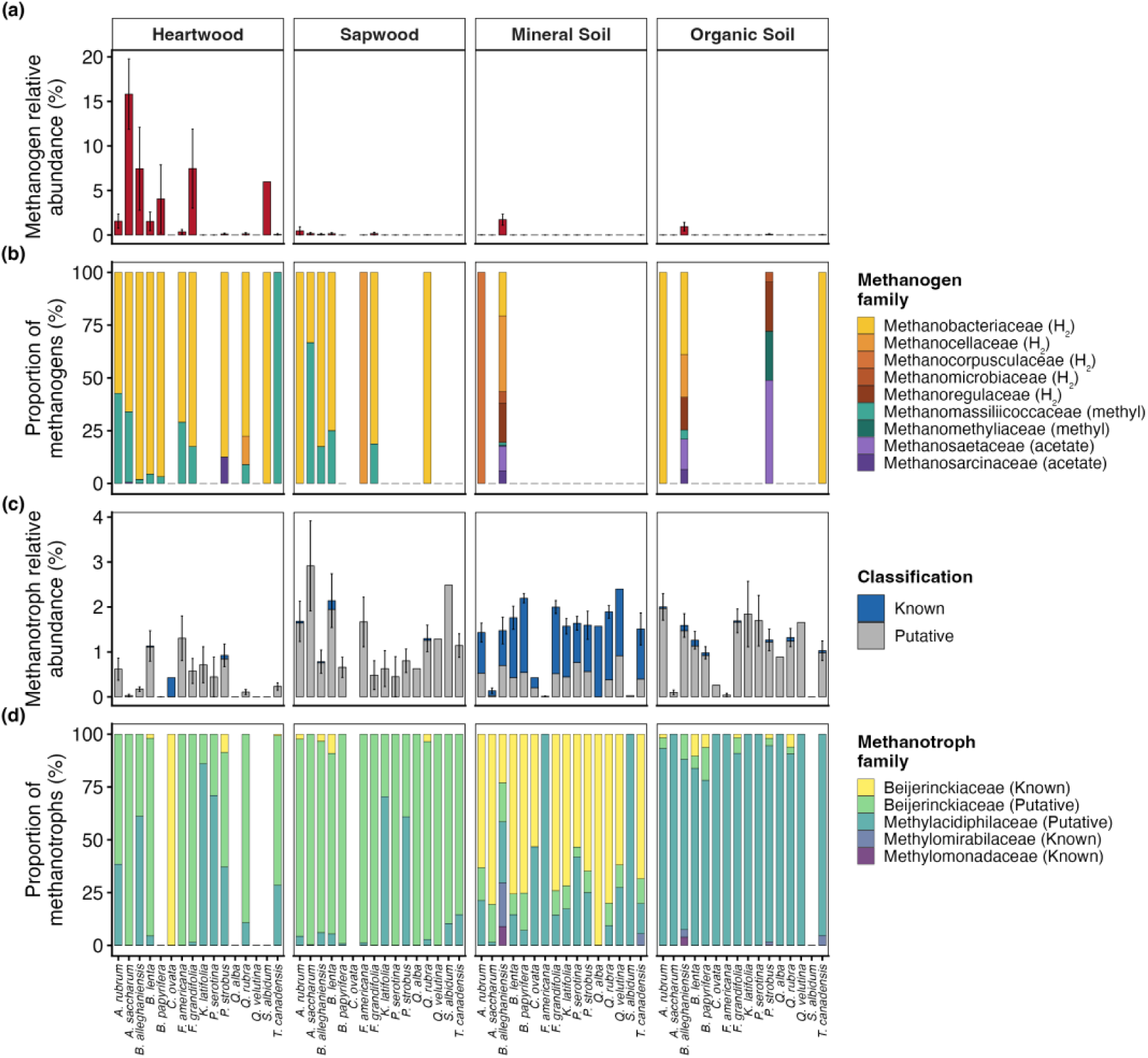
Community composition of methane-cycling microorganisms across tree species and compartments. Based on 16S rRNA amplicon sequencing, faceted by compartment. (a) Methanogen relative abundance (% of total community) by species, mean ± SE. (b) Proportional family-level composition of methanogens by species, with families shaded by methanogenic pathway (yellows to browns, hydrogenotrophic; teals, methylotrophic; purples, acetoclastic). (c) Methanotroph relative abundance by species, stacked by tier: Known (genus carries methane monooxygenase) and Putative (genus-unresolved members of methanotroph-containing families; an upper bound). (d) Proportional family-level composition of methanotrophs by species, with families labeled by tier.

The wood methanogenic community was dominated by two taxonomic families (Fig. 5b). Methanobacteriaceae comprised 2.55 ± 7.92% of heartwood communities on average but reached up to 54.7% in some samples, while in sapwood they ranged from 0 to 5.9%. Methanomassiliicoccaceae were less abundant, averaging 0.65 ± 2.43% in heartwood with a maximum of 17.5%, and 0–0.63% in sapwood. Both families were significantly more abundant in heartwood than sapwood (p < 0.001). Other methanogenic groups occurred only at trace amounts, with the third most abundant family, Methanosarcinaceae, peaking at just 1.00% of the total community. No anaerobic methane-oxidizing archaea (ANME), which also carry *mcrA*, were detected, so *mcrA* is read here as a methanogen marker.

Soil communities contained few methanogens. Methanogens comprised only 0.14 ± 0.70% of mineral soil and 0.09 ± 0.52% of organic soil communities by 16S relative abundance (Fig. 5a). Only trace amounts of the wood-dominant families were detected in soils (Methanobacteriaceae: mineral 0.028 ± 0.16%, organic 0.034 ± 0.20%; Methanomassiliicoccaceae virtually absent). Bathyarchaeia (0.23% of soil communities), a group with debated methanogenic potential (Evans *et al*., 2015), were also present but are not included in these methanogen totals.

Known methanotrophs were scarce in wood, 0.02–0.03% of heartwood and sapwood communities (Fig. 5c). Counting Putative methanotrophs raised wood to 0.55% of heartwood and 1.29% of sapwood communities (Fig. 5c–d), mostly from Beijerinckiaceae unresolved at genus level. In soils, Known methanotrophs (1.00% of the mineral-soil community) were mostly *Methylocapsa*, and the Putative tier in organic soil was mostly Methylacidiphilaceae (Methods S3).

Relative abundance from 16S and absolute abundance from ddPCR (Figs 4, 5) agreed only in part, and their detection limits differ (Methods S3). Organic soil carried as much *pmoA* as mineral soil but few methanotrophs that 16S resolved to genus, so its *pmoA* carriers are present but unresolved (Methods S3; Table S4; Fig. S4).

### Taxonomic correlations and inferred metabolic pathways

Family-level 16S relative abundance was tested against *mcrA* copy number in the same samples (Fig. 6a; Fig. S9). Beyond the methanogen families themselves (Methanobacteriaceae, Methanomassiliicoccaceae), two families were positively associated with *mcrA* at FDR < 0.05, Eggerthellaceae and Dysgonomonadaceae. Christensenellaceae and Ethanoligenenaceae, both H₂-producing fermenters, were nominally associated (p < 0.01) but did not pass correction (Fig. 6a; Methods S7). No family was associated with *pmoA* at FDR < 0.05 (Fig. S10). Predicted functions also pointed to anaerobic metabolism. With methanogen ASVs removed, fermentation and anaerobic chemoheterotrophy were enriched in heartwood relative to sapwood (FAPROTAX; paired Wilcoxon, 108 trees, FDR ≤ 0.003; Fig. 6b; Fig. S11), and samples with more *mcrA* were enriched in fermentation and carbon-fixation pathways and depleted in aerobic respiration (PICRUSt2; Fig. 6c; Figs S12–S14; Methods S7).

**Figure 6.**
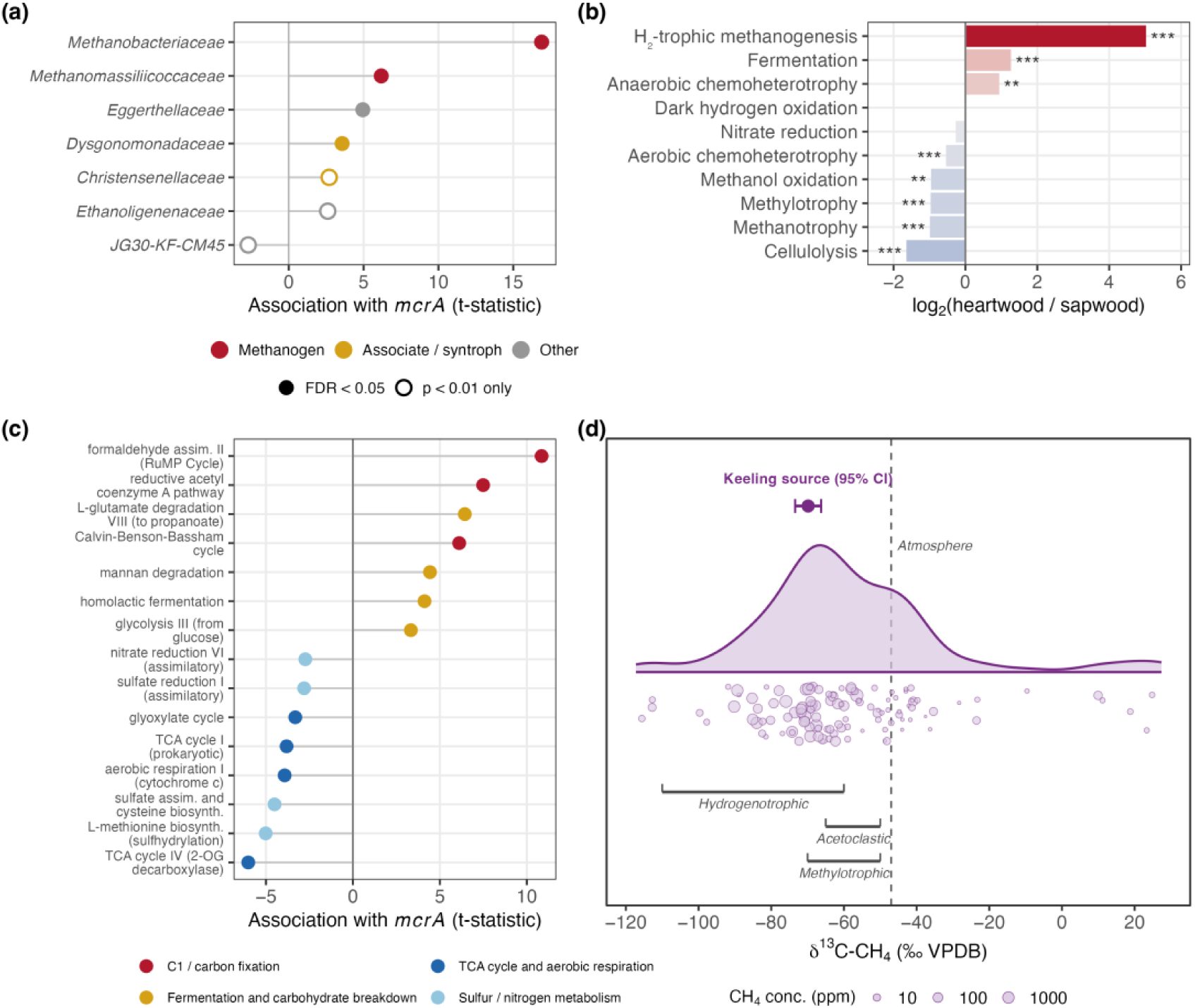
Taxonomic, functional and isotopic evidence on the methanogenic pathway in heartwood. (a) Families whose relative abundance is associated with *mcrA* in wood (heartwood and sapwood; mixed models with tree as a random intercept, controlling for compartment and 16S load; families with p < 0.01): methanogens (red), fermenters with evidence of supporting methanogens (gold) and other families (gray); filled circles, FDR < 0.05; open circles, p < 0.01 but not significant after FDR correction. (b) FAPROTAX functions as log₂(heartwood/sapwood), computed without methanogen ASVs; hydrogenotrophic methanogenesis is shown for reference. Asterisks, paired Wilcoxon signed-rank tests between heartwood and sapwood of the same tree (n = 108 trees), BH-corrected across functions: * FDR < 0.05, ** < 0.01, *** < 0.001. (c) MetaCyc pathways (PICRUSt2) associated with *mcrA* (mixed-model t-statistic; FDR < 0.05), excluding contributions from methanogen ASVs; up to four pathways per category. (d) δ¹³CH₄ of internal stem gas (n = 128 trees; points sized by CH₄ concentration; 12 values beyond the axis are included in all statistics). Brackets, ranges for each methanogenic pathway (Whiticar, 1999); point and bar, tree-produced source from the robust Keeling intercept with 95% CI; dashed line, atmospheric δ¹³CH₄ (−47‰). Panels (a)–(c) share the same 16S and *mcrA* data; the isotopes (d) are independent.

### Internal gas concentrations and isotopic composition

Internal stem CH₄ ranged from 1.3 ppm to at least 60,340 ppm (median 9.7 ppm; mean 2,544 ppm; Fig. S15), and 13% of trees exceeded 1,000 ppm. It rose with internal CO₂ and fell with internal O₂, consistent with anaerobic production. It was only weakly related to heartwood *mcrA* and to surface flux, and the relationship with flux strengthened with measurement height (Fig. S16; Methods S1).

Stable carbon isotope ratios of internal stem CH₄ (n = 128 trees across 14 species; Fig. 6d; Fig. S17) had a median δ¹³CH₄ of −63.0‰ VPDB (IQR −72.6 to −46.5‰). Taken alone, this bulk value is not diagnostic of pathway: it falls where the ranges for CO₂ reduction (about −110 to −60‰) and acetate fermentation (about −65 to −50‰) overlap (Whiticar, 1999), and samples with low internal CH₄ are diluted toward atmospheric CH₄ (−47‰). Correcting for that dilution placed the tree-produced source at −70‰, both as the robust Keeling-plot intercept (95% CI −73 to −66‰) and by two-endmember mixing with atmosphere. That is more ¹³C-depleted than the bulk median and on the CO₂-reduction side of the overlap. Sensitivity checks gave −69‰ and −79‰ (Methods S1). Internal δ¹³CO₂ (median −20.4‰) was not correlated with δ¹³CH₄ among samples with at least 10 ppm CH₄ (Spearman ρ = 0.01, p = 0.94, n = 59), consistent with a CO₂ pool dominated by respiration.

### Wood decay and within-tree heterogeneity

Stem flux varied non-monotonically with visible stem deterioration in the 2023 survey (Fig. 7a; Fig. S18): it was highest on stems with moderate bark loss and lower on healthy and on severely deteriorated or dead stems (marginal R² = 0.05, p < 0.001; a quadratic was preferred over a linear term, ΔAIC = 10.3). Wood *mcrA* abundance also peaked at intermediate fungal ITS load, a proxy for fungal colonization (Fig. 7b; R² = 0.08, p < 0.001), while soil *mcrA* did not. We treat this relationship as exploratory.

**Figure 7.**
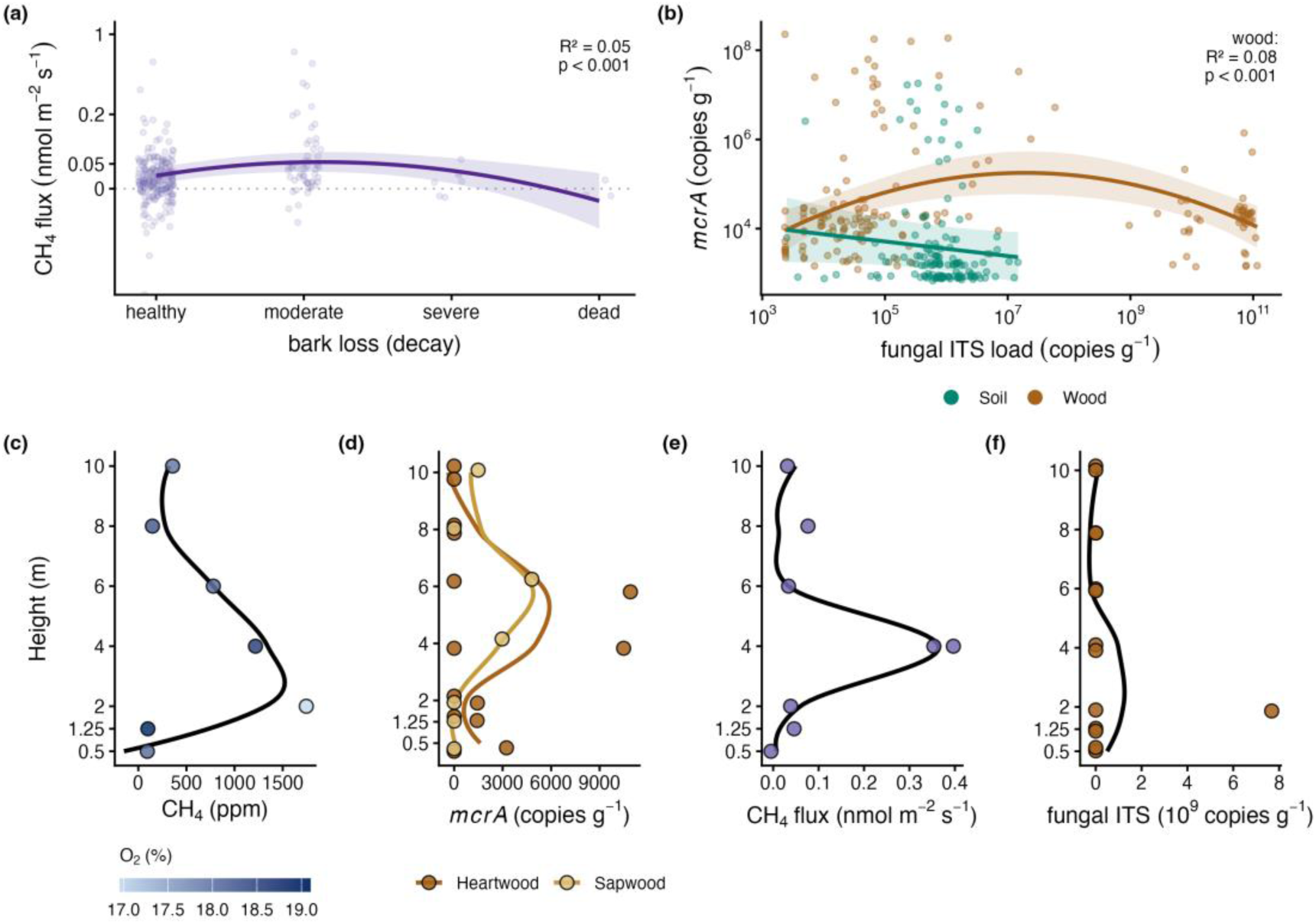
Wood decay, fungal colonization and the vertical distribution of methanogenesis. (a) Stem CH₄ flux (nmol m⁻² s⁻¹, arcsinh axis) against visible bark loss in the 2023 survey (healthy, moderate, severe, dead); line and band, quadratic mixed model with species as a random effect, ±95% CI. Dotted line, zero flux. (b) *mcrA* abundance against fungal ITS load (copies g⁻¹, log axes; wood, dry mass; soil, fresh mass) in wood (brown) and soil (teal) from the 2021 survey; lines and bands, mixed-model fits ± 95% CI (quadratic for wood, linear for soil). (c–f) Vertical profiles in the felled black oak (*Ǫuercus velutina*): (c) internal CH₄ (ppm), points colored by internal O₂ (%); (d) heartwood (brown) and sapwood (tan) *mcrA* (copies g⁻¹ dry wood); (e) stem CH₄ flux (nmol m⁻² s⁻¹); (f) heartwood fungal ITS load (copies g⁻¹ dry wood). Curves in (c–f) are smoothers drawn as guides to the eye.

Within the felled black oak (Fig. 7c–f), internal CH₄ was highest at 2 m (1,745 ppm) and 4 m (1,217 ppm). Emission peaked at 4 m (0.35 nmol m⁻² s⁻¹, four to eleven times that at other heights), near the upper limit of the decay column, which ran from a basal cavity through stained wood to clean wood above about 6 m (Fig. S19); it was not detected at 0.5 m and was still 0.03 nmol m⁻² s⁻¹ at 10 m. *mcrA* was detected in 8 of 21 cores, at up to about 11,000 copies g⁻¹ dry heartwood at 4–6 m, and at least one core was below detection at every height, including heights with elevated internal CH₄.

Methane-cycling taxa occurred in every tissue sampled, from heartwood to foliage, roots and the surrounding soil (Fig. S20). Known methanotrophs, almost all *Methylocella*, were concentrated in the outer and aerial tissues: 0.50% of bark, 0.60% of branch and 0.62% of foliage communities, against none in sapwood and 0.02% in heartwood (one tree; detected in two to three of four libraries per tissue; Table S4).

### Gene–flux relationships across scales

At the level of individual trees, gene abundances explained little of the variation in flux (Fig. S21). Area-weighted *mcrA* was essentially unrelated to flux across 125 trees (R² < 0.001), and methanotroph genes and the ratio explained little more (R² ≤ 0.04; within species, only *mmoX* was associated with flux, Methods S4). Methanogens were patchy within stems (Fig. 7c–f; Fig. S22). A random forest using the three genes predicted held-out trees no better than their mean (R² = −0.01), and adding them to species, diameter and soil moisture did not improve prediction (held-out R² 0.18 with and without; Methods S4).

Aggregated to species, the same data were informative. Across ten species with at least five trees, median area-weighted *mcrA* was positively related to median flux (R² = 0.30, p = 0.10), methanotroph abundance was negatively related (R² = 0.20, p = 0.19), and the ratio was the strongest predictor (R² = 0.46, r = 0.67 on the log ratio, p = 0.032; Fig. 8). The ratio outperformed *mcrA* alone (AIC −44.5 against −41.9). Methanogen and methanotroph abundances were not correlated with each other, either among trees (r = 0.01, p = 0.94) or among species (r = −0.04, p = 0.90; Fig. S23).

**Figure 8.**
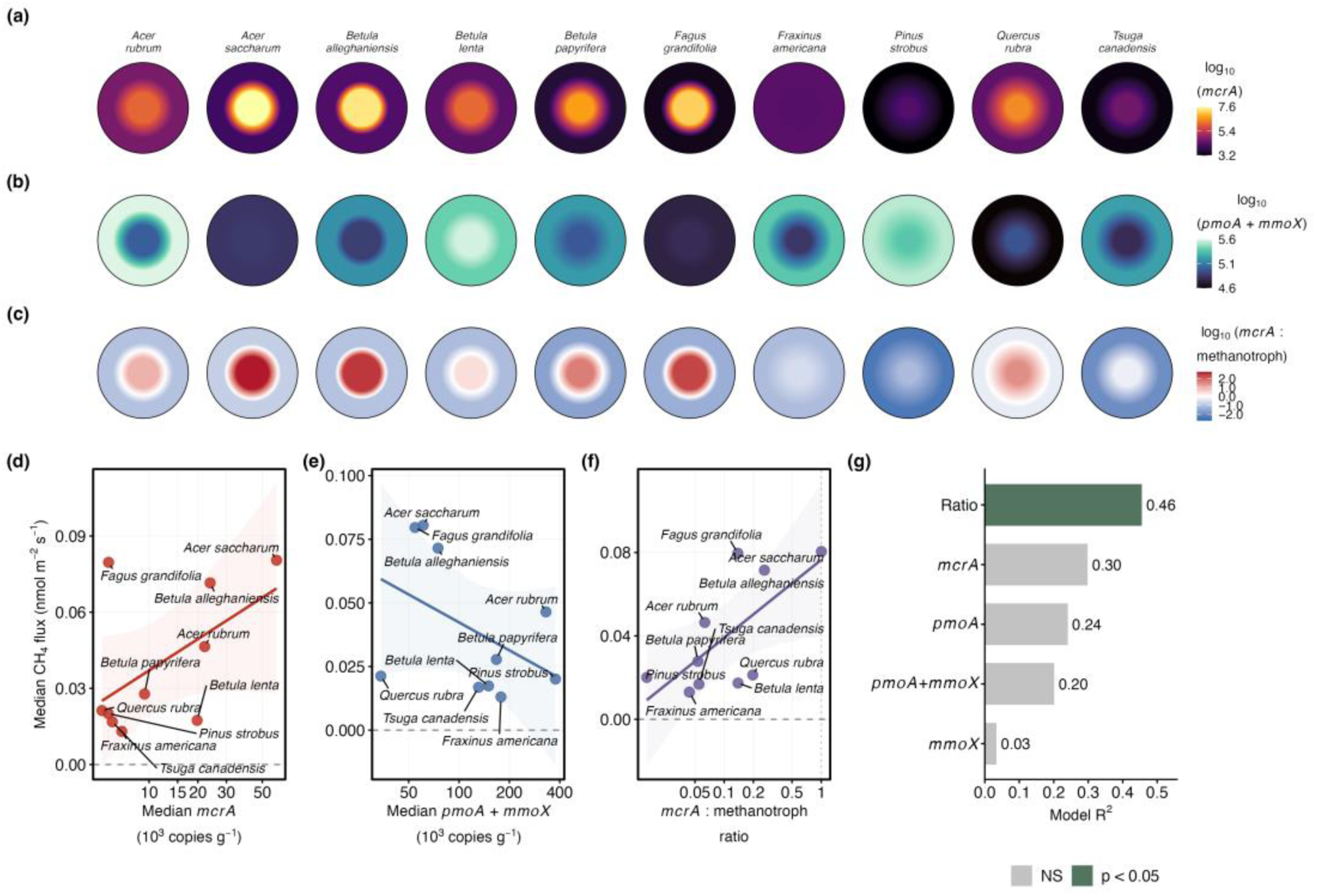
Species-level relationships between functional gene abundance and CH₄ flux. (a–c) Radial cross-sections showing (a) log₁₀(*mcrA*), (b) log₁₀(*pmoA* + *mmoX*) and (c) log₁₀ methanogen:methanotroph ratio (*mcrA*:(*pmoA* + *mmoX*)) in heartwood (center) and sapwood (outer ring) for ten tree species. Cross-sections are drawn at equal size; species in alphabetical order. In (c) the diverging scale separates methanogen-dominated (red) from methanotroph-dominated (blue) wood. (d) Species-median area-weighted *mcrA* against median CH₄ flux. (e) Species-median methanotroph abundance against median CH₄ flux. (f) Species-median methanogen:methanotroph ratio against median CH₄ flux (R² = 0.46, p = 0.032). Lines and bands in (d–f), linear regression ± 95% CI. (g) R² across five gene-abundance metrics; significant predictors (p < 0.05) in green.

Three species properties were associated with species-level gene abundances, each in the same direction across all species and among broadleaf species alone (Fig. S24; Methods S5). Species with denser bark had higher methanogen:methanotroph ratios (all species with data ρ = 0.85, p = 0.006; broadleaf ρ = 0.79, p = 0.048), species of wetter soils carried more methanogens (ρ = 0.62, p = 0.060; ρ = 0.76, p = 0.037), and longer-lived species carried fewer methanotrophs (ρ = −0.31, p = 0.38; ρ = −0.85, p = 0.007). In each case the two conifers differed from each other in the same direction as the broadleaf trend. None passed false-discovery correction across the 52 trait tests (q ≥ 0.32), and no trait was related to net flux.

### Stand-scale bounds

Random forests reproduced the seasonal pattern of stem and soil fluxes (Fig. S25). With whole trees or collars held out, R² was 0.10 for stems and 0.39 for soils (out-of-bag 0.23 and 0.41), so individual measurements were poorly predicted; held-out predictions averaged about 5% above observations (mean predicted:observed 1.05), and stand totals use a per-species calibration (Fig. S26). Soil moisture dominated the soil model; in the stem model, species and soil moisture contributed most, and stem flux was not explained by environment alone (Methods S6).

Per unit of bark surface, stems in the measured 0–2 m band emitted 0.090 nmol m⁻² s⁻¹ averaged over the year (monthly 0.079–0.099), with breast-height fluxes peaking in August; soils took up 0.87 nmol m⁻² s⁻¹ per unit ground (monthly 0.44–1.30). Per unit area, stem emission was therefore about a tenth of soil uptake, but the band’s bark area is only 12.0% of the stand area.

Integrated over that band, stem emissions totaled 5.47 mg CH₄ m⁻² yr⁻¹, offsetting 1.24% of soil uptake (440 mg CH₄ m⁻² yr⁻¹; both per m² of ground). This band, the only part of the budget with both flux and surface area measured, covers live stems; the inventory did not record standing dead stems or coarse woody debris. Extended to the whole woody surface (assumptions in Fig. S27), stem emission ranged from −16 to 101 mg CH₄ m⁻² yr⁻¹ across all 240 combinations, and from 6.0 to 101 mg (1.4–23% of soil uptake) without the linear-into-uptake form (Fig. S28). How flux changes above 2 m (staying constant, declining, or turning into uptake) moved the median estimate across a range about ten times that of woody area (93 against 10 mg), and leave-one-height-out cross-validation gave the lowest error for a constant when extrapolating upward (root-mean-square error 0.41 against 6.7 for exponential decay; Fig. S27). The stand remained a net CH₄ sink in all 240 combinations (net −455 to −339 mg CH₄ m⁻² yr⁻¹; Fig. 9d).

**Figure 9.**
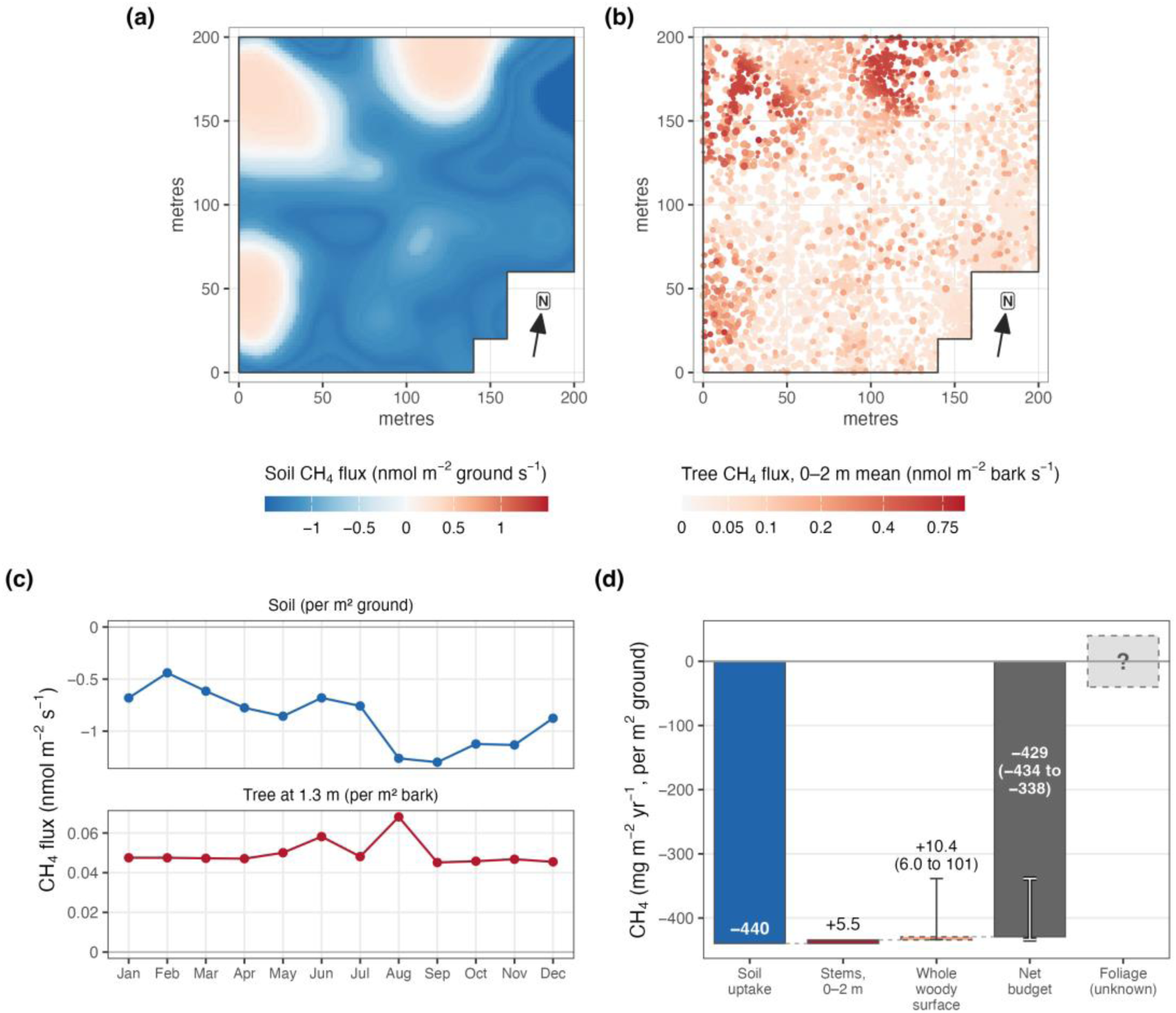
Stand-scale CH₄ exchange in the upland inventory plot: measured band and bounds. (a) Annual mean soil CH₄ flux (nmol m⁻² ground s⁻¹) predicted across the censused stand; blue, uptake; red, emission. (b) Annual mean stem CH₄ flux (nmol m⁻² bark s⁻¹, 0–2 m band mean) for each inventoried stem, point size proportional to diameter at breast height. Both maps are random-forest predictions (held-out R² 0.39 soil, 0.10 stem). Arrow points north. (c) Monthly soil flux (per m² ground) and stem flux at breast height (per m² bark). (d) Annual budget (mg CH₄ m⁻² ground yr⁻¹): soil uptake; stem emission over the measured 0–2 m band; the whole woody surface if flux declines above 2 m as it did between 0.5 and 2 m (dashed; the bar shows the increment above the measured band and the label the total; 84 if flux instead holds at its 2 m value); the net budget under the same case; and foliage, not measured (box size arbitrary). Whiskers and bracketed values are scenario ranges across the 200 scaling combinations that exclude the linear-into-uptake form, not standard errors or confidence intervals; with that form included, whole-surface emission spans −16 to 101 and the net budget −455 to −339.

Two readings of the height measurements bracket the whole surface at the woody area index estimated from this stand’s inventory (2.93; range 2.32–3.53, Fig. S27). If flux declines above 2 m as it did between 0.5 and 2 m, a decline carried by *B. alleghaniensis* (Fig. 2), stems emit 10.4 mg CH₄ m⁻² yr⁻¹ (2.4% of soil uptake); if it holds at its 2 m value, as in most species and in the one tree measured to 10 m (Fig. 7e), they emit 84 mg (19%; Methods S6).

## Discussion

### Upland stems as net methane sources

Upland trees emitted methane where the surrounding soils consistently consumed it (Fig. 1), and they remained net sources although methanotroph genes were present in nearly every sample (Fig. 4). Over the lowest 2 m, fluxes were small, net positive and roughly uniform with height in most species (Fig. 2), and the one tree we profiled to 10 m was still emitting at the top (Fig. 7e). Upland trunk profiles to 12 m, where most emission came from above 3 m (Mochidome *et al*., 2025), and a survey of eleven species (Epron *et al*., 2025) also found emission well above 2 m, whereas upland woody surfaces measured by Gauci *et al*. (2024) switched from net emission near the stem base to net uptake by about 2 m. All of these studies measure net flux, which is gross production minus consumption, and none measures the two terms separately. Net emission from lower stems is the common finding, and the sign higher on stems remains unresolved for most forests (Covey & Megonigal, 2019; Gauci *et al*., 2024). Because the net flux is small, a modest shift in the balance, such as drier wood, more active bark methanotrophs or lower internal production, could reverse its sign elsewhere. In our data, the few apparent stem uptakes track instrument precision, consistent with measurement noise (Methods S2).

### Where stem methane is produced

Our data constrain production more directly than consumption. Methanogens occurred in 97% of heartwood samples, concentrated in heartwood (Fig. 4), and wood from this forest produces methane in anaerobic incubation (Arnold *et al*., 2025). Hydrogenotrophic Methanobacteriaceae were about four times as abundant as Methanomassiliicoccaceae, which reduce methyl compounds with H₂ (Fig. 5b) (Borrel *et al*., 2014; Boone *et al*., 2015; Buan, 2018; Cozannet *et al*., 2021), and the dilution-corrected source δ¹³CH₄ of about −70‰ (Fig. 6d) points the same way: CO₂ reduction with H₂ is likely the primary pathway in wood, with H₂-dependent methyl reduction secondary. Isotopic interpretation is complicated by co-occurring pathways and by fractionation during oxidation, which enriches residual δ¹³CH₄ (Jeffrey *et al*., 2021b), but values this depleted are difficult to explain without substantial hydrogenotrophic production. Wood can also supply methyl groups directly: in anoxic peat, the methoxy groups of lignin monomers were converted to CH₄ (Liu *et al*., 2025).

Fermenters likely supply the H₂. Two families of anaerobic fermenters, Eggerthellaceae and Dysgonomonadaceae, co-varied with *mcrA*, and two H₂-producing fermenters, Christensenellaceae and Ethanoligenenaceae, did so nominally (Fig. 6a; Methods S7); samples with more *mcrA* were predicted to be richer in fermentation and poorer in aerobic respiration (Fig. 6c), consistent with the higher methane and more variable oxygen of heartwood than sapwood in this forest (Arnold *et al*., 2025).

In these upland stems, internal production appears to be the main source, with soil-derived methane added where soils are methanogenic. Only in *B. alleghaniensis* did flux decline strongly and consistently with height, the pattern expected from soil-to-stem transport (Fig. 2). This species grew on wetter microsites, where soil methanogen abundance was highest, and also carried high heartwood *mcrA* (Fig. 4a), so both sources likely contribute there. Soil-to-stem transport is the dominant pathway in wetland forests (Pangala *et al*., 2013, 2017; Barba *et al*., 2019a; Jeffrey *et al*., 2019), and stem emissions rise on wetter soils (Ranniku *et al*., 2024), so its share likely grows as sites become wetter and soils shift from net sinks to net sources. Internal production, by contrast, should operate on wet and dry soils alike and give low, uniform profiles. Such profiles were also found in birch and spruce on drained peat, whose stem emissions were unrelated to soil flux (Ranniku *et al*., 2023), and in upland trunks where methane was produced at height (Mochidome *et al*., 2025).

Nonetheless, we cannot fully rule out soil-origin contributions on net-methanotrophic soil. Deep soil layers below the oxidation zone harbor methanogens whose products may be able to bypass oxidation through root uptake and xylem transport, a route shown for wetland plants and rice but not yet confirmed for upland trees (Nouchi *et al*., 1990; Schimel, 1995; Watson *et al*., 1997; Bradford *et al*., 2001; Bhullar *et al*., 2013), and anoxic microsites within aerobic soils (aggregate interiors, root detrituspheres, or temporarily saturated zones) act as localized sources (Zausig *et al*., 1993; Keiluweit *et al*., 2018; Lacroix *et al*., 2023). Tracers, metatranscriptomics and root-severing experiments could separate internal from external sources directly.

### Methane-cycling communities inside and outside the stem

The methane-cycling community of wood differed from that of the surrounding soil. Wood methanogens (Methanobacteriaceae, Methanomassiliicoccaceae; Fig. 5b; Fig. S9) were virtually absent from soils, consistent with heartwood microbiomes that are taxonomically distinct from surrounding substrates in living (Cregger *et al*., 2018; Arnold *et al*., 2025) and dead wood (Moll *et al*., 2018), and suggesting selection for lineages adapted to wood. Within the stem, methanogens were more abundant in heartwood and methanotrophs relatively more so in sapwood (Figs 4, 5), with production concentrated where oxygen is depleted and consumption capacity where it remains available. These are plant tissues with different physiology. Sapwood conducts sap and keeps living parenchyma, so it stays connected to the atmosphere through the transpiration stream, whereas heartwood formation ends sap flow and cell activity and deposits species-specific extractives, isolating wood from that supply. Because heartwood and sapwood were defined here operationally, by depth in the core, separating the effects of tissue and depth is a question for future study. Methanogens were likewise concentrated in wood, rather than in bark or leaves, in five floodplain tree species (Moisan *et al*., 2026), and in heartwood rather than sapwood in eastern cottonwood, whose wood communities differed from those of soil (Yip *et al*., 2019).

Methanotrophs showed the opposite pattern: their signal in wood was weak. Known methanotrophs made up only 0.02–0.03% of heartwood and sapwood communities (Fig. 5c). Counting Putative ones raises this but probably overcounts. The largest Putative group in wood was genus-unresolved Beijerinckiaceae, and most wood Beijerinckiaceae that could be resolved belonged to genera with no annotated methane monooxygenase, such as *Lichenibacterium* (Table S3). Phylogenetic placement put only 9–14% of the wood Putative signal close to a methanotroph genus (Methods S3; Table S4). The 16S census may also miss wood methanotrophs: at our read depth it cannot detect taxa below about 0.03% of a sample, reference genomes from wood are scarce, and *pmoA* was detected by ddPCR in 95–99% of wood samples although 16S named few methanotrophs there (Fig. S8; Methods S3). Soils carried different taxa: *Methylocapsa* in mineral soil and, in organic soil, Putative, genus-unresolved Methylacidiphilaceae, whose methane monooxygenase is known only from geothermal acidophiles (van Teeseling *et al*., 2014) (Fig. 5d).

The survey cored only wood; in the one tree sampled in every tissue, Known methanotrophs were concentrated in bark, branches and foliage (Fig. S20; Table S4), as reported for bark communities elsewhere (Jeffrey *et al*., 2021a; Leung *et al*., 2026). They were almost all *Methylocella*, a genus also found in the bark of an upland eucalypt (Leung *et al*., 2026), whereas wetland bark is dominated by *Methylomonas* (Jeffrey *et al*., 2021a; Leung *et al*., 2026). The wood survey therefore gives a lower bound on methanotrophs at the stem surface. Such results depend greatly on how methanotrophs are defined from taxonomic data, in the absence of metagenomes or direct activity measurements: a recent report of a methane-oxidizer signal in floodplain bark and leaves, for example, was dominated by *Methylobacterium*–*Methylorubrum* (Moisan *et al*., 2026), which may lack methane monooxygenase (Methods S3; Table S7).

The *mmoX* data point to which consumers matter in wood. *Methylocella* carries sMMO (*mmoX*) but not pMMO. In soil the two genes rose together, whereas in wood they varied independently (Fig. S8), and sapwood *mmoX* was the only gene associated with flux within species, negatively (Methods S4). Cells express sMMO when they cannot acquire enough copper, and its lower affinity for CH₄ suits elevated rather than atmospheric concentrations (Semrau *et al*., 2010). sMMO-bearing methanotrophs in wood are therefore better placed to consume internally produced methane, which reached thousands of ppm in some stems (Fig. S15), than atmospheric methane, whose uptake depends on high-affinity enzymes and on redox and trace-metal conditions (Gauci *et al*., 2024; Davidson *et al*., 2024; Leung *et al*., 2026).

Methanotrophy overall remains the less understood process in trees. Bark methanotrophs act on stem fluxes: sterilizing bark increased stem emissions by about 36% (Jeffrey *et al*., 2021a), and gross bark methanotrophy has been confirmed independently (Leung *et al*., 2026). Measured fluxes therefore already include some consumption, of internally produced and possibly of atmospheric CH₄, so gross production in stems exceeds net emission. Which organisms oxidize CH₄ in and on stems, and at what rates, can be settled only by activity measurements (rates, transcripts, stable-isotope probing) and by reaction–transport models that couple microbial physiology with transport in wood (Megonigal *et al*., 2020).

### Scale dependence in predicting net flux

Between individual trees, neither methanogen nor methanotroph abundance explained much of the variation in flux (R² ≤ 0.04; Fig. S21). Gene counts are a census of potential producers and consumers: they cannot distinguish active from dormant cells, and gene abundance often decouples from process rates (Rocca *et al*., 2015). Production is also patchy within stems. In the felled oak, methanogenic hotspots lay at 4–6 m (Fig. 7c–f), yet increment cores, each sampling a tiny fraction of the stem beneath a single chamber, missed methanogens in at least one core at every height, including the emission peak. A single core is therefore a noisy measure of a tree’s methanogen load, and such measurement error weakens tree-level correlations through regression dilution (Frost & Thompson, 2000). The patchy detections suggest localized production sites such as decay pockets, wound responses, or other anaerobic niches (Waring *et al*., 2016; Nunan *et al*., 2020), from which methane spreads well beyond its source.

Decay is a plausible example of one such niche: in our survey, flux peaked on moderately deteriorated stems and wood *mcrA* at intermediate fungal load (Fig. 7a–b), and in the felled oak emission peaked near the top of the decay column (Fig. S19); in sugar maple, stems with more extensive heart rot, mapped by sonic tomography, emitted more CH₄ (Senevirathne *et al*., 2026), and in a cool-temperate upland forest emission peaked at 4–6 m in some trunks, where production was linked to wood degradation (Mochidome *et al*., 2025).

Transport and oxidation inside the stem further separate production from surface flux. In our survey, internal CH₄ was only weakly related to surface flux (Fig. S16), and heartwood CH₄ monitored monthly for 15 months likewise predicted emission only weakly, with apparent diffusion coefficients that differed among trees (Epron & Mochidome, 2024; Pangala, 2025). Methane produced in one place may emerge in another, and oxidation during diffusion through aerobic zones complicates inferring production from surface flux. Wood porosity, vessel anatomy (ring-porous vs. diffuse-porous), sapflow (Barba *et al*., 2019b, 2021; Anttila *et al*., 2024; Bréchet *et al*., 2025) and radial cell architecture (Côté, 1963; Sorz & Hietz, 2006) create species- and tree-specific transport regimes. Transport also decides whether lower-stem measurements represent whole trees: in wetlands, soil-transported methane declines exponentially with height (Pangala *et al*., 2013; Barba *et al*., 2019a; Jeffrey *et al*., 2024), whereas internally produced methane may sustain emission higher on the stem. How long methane resides in wood, and whether it follows preferential paths such as vessels, rays or decay channels, remain to be measured; multi-point coring at several depths and heights, imaging of wood anatomy, and gas sampling with isotopic tracers could locate sources and paths.

Yet, averaged over within-tree heterogeneity, the genes became informative. Across species, the methanogen:methanotroph ratio was the strongest predictor of net flux (Fig. 8) and is the closest our data come to the production–consumption balance itself: species whose wood is tilted toward methanogens emit more, and the ratio accounted for 46% of the variation among species. Aggregation and ratios carry statistical risks (Jasienski & Bazzaz, 1999; Bradford *et al*., 2017), but the species pattern held in direction for the numerator and denominator separately (Fig. 8d–e). Species means are also the scale at which flux integrates over within-tree heterogeneity (Levin, 1992; Polussa *et al*., 2021), and the usual unit for relating function to traits in comparative ecology, from tree effects on litter decomposition (Hobbie *et al*., 2006) and tree growth and mortality (Poorter *et al*., 2008) to mammalian metabolic rate (White & Seymour, 2003).

### Plant traits and species differences

Tree species differed in flux and in the balance between methanogens and methanotrophs, and properties of the trees themselves may explain part of that difference. Methanogen loads were highest in *A. saccharum* heartwood and low in conifers, particularly in sapwood (Fig. 4; Methods S1), pointing to wood chemistry or heartwood formation. Species differences may also reflect ecology more than taxonomy. Two birch species bracketed the range: *B. alleghaniensis* emitted most and *B. lenta* least (Fig. 3a). *B. alleghaniensis* grows on the wettest, most methanogenic soils at our study site, whereas *B. lenta* is found on drier sites than *B. alleghaniensis* and is regenerating widely in the study forest and region as hemlock and oak decline, so its trees are likely younger, with less of the heart rot that supports methanogens (Methods S4).

The trait associations (Fig. S24; Table S8) differ by group: methanogens track hydrology (species of wetter soils carry more), methanotrophs track longevity (fewer in long-lived broadleaf species), and the ratio is higher in species with dense bark. Species of wet soils may hold wetter wood, in which oxygen diffuses slowly and anoxic microsites persist, or may draw in methanogens with saturated soil water. Dense bark may restrict gas exchange with the outer wood, and long-lived, slow-growing species build denser, nitrogen-poor and more chemically defended wood; both could limit aerobic methanotrophs. None of these associations passed correction for multiple tests (Methods S5), so we offer them as exploratory hypotheses implicating plant tissue density, life history and hydrology in the stem methane balance, to be tested across more species.

### Implications for forest methane budgets

At the upland study site, stem emissions over the measured 0–2 m band offset 1.24% of soil uptake (5.47 against 440 mg CH₄ m⁻² yr⁻¹; Fig. 9d); in a young oak plantation the figure was 0.1% (Plain *et al*., 2019). Beyond that band, our estimates are bounds rather than a budget, and conservative ones, because no scenario lets flux rise above 2 m as it did in the felled oak (Fig. 7e): across every combination of woody area, its vertical distribution and how flux changes above 2 m, whole-surface emission spans −16 to 101 mg CH₄ m⁻² yr⁻¹, up to almost a quarter of soil uptake. How flux changes above 2 m is the main uncertainty: it moves the total about ten times as much as woody area does, and the temperate woody area index of 3.07 used by Gauci *et al*. (2024) falls inside our range for this stand (2.32–3.53). The stand stays a net sink in every combination, but these bounds omit the canopy, where foliage can emit methane, as Scots pine shoots do (Machacova *et al*., 2016; Tenhovirta *et al*., 2022), or possibly take it up, as the leaves of several temperate tree species have recently been reported to do (Gorgolewski *et al*., 2023; Karim & Thomas, 2026).

Extrapolating to stand- and ecosystem-scale budgets needs three advances. First, vertical profiling across trees and species must show whether fluxes measured below 2 m scale to the whole stem. Second, total woody surface area can be better constrained with direct three-dimensional measurement rather than allometric estimates, which terrestrial laser scanning can provide (Calders *et al*., 2020). Third, and perhaps most importantly, tower-based or airborne eddy covariance can constrain tree contributions by comparing net ecosystem and soil fluxes (Pangala *et al*., 2017). In a California oak savanna, chambers combined with eddy covariance and laser-scanned stem area showed stems to be net sources all year and the ecosystem a net CH₄ source despite soil uptake (Kasak *et al*., 2026). However, few upland towers measure methane, tree signals are small against soil uptake, and wetland patches make tree contributions hard to isolate (Delwiche *et al*., 2021).

### Conclusions

Upland temperate trees harbor abundant, taxonomically distinct methane-cycling communities. Methanogens were detected in nearly all heartwood samples, about an order of magnitude above surrounding soils, and isotopic and taxonomic evidence points to hydrogenotrophic methanogenesis supported by fermenters. Stem emissions in most species were consistent with internal production, and stems remained net sources despite methanotroph genes in nearly every sample. At our site, stem emissions below 2 m offset about 1% of soil uptake and the whole woody surface 2.4–19%, depending on whether flux declines or holds above 2 m, and the stand remained a net sink under every assumption we tested. Net flux reflects the balance between production and consumption, and across species the methanogen:methanotroph ratio accounted for 46% of the variance in emission, so changes in either process could alter the contribution of trees to forest methane budgets.

## Supporting information

Notes S1: ddPCR protocol for methane-cycling genes

## Acknowledgements

We thank Josep Barba, Makenzie Birkey, Cade Brown, Claire Butler, Ari Gewirtzman, Ben Girgenti, Thomas Harris, Naomi Hegwood, Marsh Hlavka, Luke Jeffrey, Fiona Jevon, Ellie Jose, Jonas Karosas, Talia Kolodkin, Camila Ledezma, Laura Logozzo, Taylor Maavara, Jackie Matthes, Naomi Norbraten, Joseph Orefice, Alex Polussa, Andrew Reinmann, Judith Rosentreter, Adriana Rubenstein, Michelle Spicer, Cyrena Thibodeau, Les Welker, and Qespi Wood for their contributions to this work. We acknowledge our additional collaborators on the forest inventory plot: Liza Comita, Mark Ashton, Simon Queenborough, Stuart Davies and Sean McMahon. We thank Yale School Forests, University of Minnesota Genomics Center, Yale Analytical and Stable Isotope Center, W.M. Keck Biotechnology Resource Laboratory, Yale Center for Genetic Analyses of Biodiversity, and Yale Chemistry Glass Shop for technical support and facilities access. We thank the TRY initiative on plant traits (http://www.try-db.org) and the contributors of the datasets used. Claude (Anthropic) was used to assist with editing and formatting the manuscript; all scientific content, analyses, and interpretations are the work of the authors.

## Author Contributions

J.G., W.A., M.T., P.A.R., J.P., and M.A.B. conceptualized the research. J.G., W.A., M.T., D.J.W., C.R.B., M.C.D., P.A.R., J.P., and M.A.B. designed the methodology. J.G., W.A., M.T., H.B., D.J.W., N.W., K.K., and L.G. conducted the investigation. J.G., W.A., M.T., H.B., D.J.W., N.W., K.K., and L.G. curated the data. J.G., W.A., and C.M. conducted the formal data analysis and developed the software. J.G. and W.A. validated the results. J.G., W.A., and C.M. prepared the visualizations. J.G., W.A., C.R.B., M.C.D., P.A.R., J.P., and M.A.B. acquired funding and provided resources. J.G., W.A., J.P., and M.A.B. administered the project. J.G., W.A., P.A.R., J.P., and M.A.B. supervised the research. J.G. wrote the original draft of the manuscript. All authors reviewed and edited the manuscript. J.G. and W.A. contributed equally to this work.

## Data Availability

The 16S rRNA amplicon sequencing data generated in this study have been deposited in the NCBI Sequence Read Archive (SRA) under BioProject accession number PRJNA1124946 (https://www.ncbi.nlm.nih.gov/bioproject/PRJNA1124946). All methane flux measurements, methanogen and methanotroph abundance data (ddPCR), internal gas concentration data, tree and soil characterization data, and associated metadata are archived in Zenodo (https://doi.org/10.5281/zenodo.24092176). R code for statistical analyses, random forest models, variance partitioning, and figure generation is available on GitHub: https://github.com/jgewirtzman/tree-methanogens. Supporting Information containing additional figures, methods, and results is provided with this manuscript.

## Supporting Information

The following Supporting Information is available for this article:

**Methods S1.** Field and laboratory procedures

**Methods S2.** Detection limits and apparent stem uptake

**Methods S3.** Methanotrophs at several taxonomic resolutions and phylogenetic placement

**Methods S4.** Gene–flux relationships at the tree and species levels

**Methods S5.** Plant traits

**Methods S6.** Random forests and stand-scale bounds

**Methods S7.** Inferred functions (FAPROTAX and PICRUSt2)

**Notes S1.** Protocol: quantifying methane-cycling genes in living wood and soil by droplet digital PCR (separate file)

**Figure S1.** Study site, soil moisture and tree species.

**Figure S2.** Stem flux chambers.

**Figure S3.** Probe-based *mcrA* assay compared with the EvaGreen assay on the same extracts.

**Figure S4.** 16S group abundance against the gene measured by ddPCR in the same sample.

**Figure S5.** Height slope of stem flux against soil moisture.

**Figure S6.** Realized soil-moisture niche of each species.

**Figure S7.** Gene abundance by species for all 16 species, with *pmoA* and *mmoX* shown separately.

**Figure S8.** *pmoA* and *mmoX* by compartment, and their coupling in wood and soil.

**Figure S9.** Family-level 16S taxonomy associated with *mcrA* abundance.

**Figure S10.** Family-level 16S taxonomy associated with *pmoA* abundance.

**Figure S11.** FAPROTAX functional predictions in tree wood.

**Figure S12.** MetaCyc pathways associated with *mcrA* abundance, excluding methanogen contributions.

**Figure S13.** MetaCyc pathways associated with *mcrA* abundance, all ASVs.

**Figure S14.** MetaCyc pathways associated with *pmoA* abundance.

**Figure S15.** Internal CH₄ concentrations by species.

**Figure S16.** Internal gases, *mcrA* and flux.

**Figure S17.** Source signatures of internal CH₄ and CO₂.

**Figure S18.** Stem flux against visible deterioration (2023 survey).

**Figure S19.** Cross-sections of the felled black oak at each sampled height.

**Figure S20.** Methane-cycling taxa across tissues of the felled black oak.

**Figure S21.** Gene–flux relationships across scales of aggregation.

**Figure S22.** Radial *mcrA* distribution in individual trees of ten species.

**Figure S23.** Independence of methanogen and methanotroph abundance.

**Figure S24.** Plant traits and the stem methane-cycling community.

**Figure S25.** Random-forest models of stem and soil flux.

**Figure S26.** Calibration of random-forest predictions.

**Figure S27.** The scaling assumptions.

**Figure S28.** Whole-woody-surface stem emission across the scaling grid.

**Table S1.** ddPCR assays: primers, probe and thermocycling.

**Table S2.** Methane-cycling taxa in the 16S data and their classification.

**Table S3.** Genome screen for methane monooxygenase in genera within mixed methanotroph families.

**Table S4.** Methanotrophs in the 16S data at several taxonomic resolutions, their agreement with ddPCR, and the felled oak by tissue.

**Table S5.** Agreement between ddPCR and 16S detection in the same samples, by compartment.

**Table S6.** Diameter at breast height of measured trees by campaign, location, species and status.

**Table S7.** Methanotroph identifications reported from trees and plant tissues, and the evidence behind them.

**Table S8.** Sources of the species trait values.

