## Supplemental Data 1 for "Methane emissions from upland tree stems reflect a balance of microbial production and consumption"

Each section gives the methods for one topic together with its supporting results.

- Methods S1. Field and laboratory procedures
- Methods S2. Detection limits and apparent stem uptake
- Methods S3. Methanotrophs at several taxonomic resolutions and phylogenetic placement
- Methods S4. Gene–flux relationships at the tree and species levels
- Methods S5. Plant traits
- Methods S6. Random forests and stand-scale bounds
- Methods S7. Inferred functions (FAPROTAX and PICRUSt2)

Notes S1 (a step-by-step protocol for quantifying methane-cycling genes in wood and soil by ddPCR) is supplied as a separate file. All data and code are archived as described in the main text (Data Availability).

#### Methods S1. Field and laboratory procedures

##### Campaigns

The monthly survey measured 48 trees (23 upland, 12 intermediate, 13 wetland-margin), including eight standing dead stems, and 30 soil collars in 11 campaigns from June 2020 to May 2021. The 2021 survey measured 150 trees of 16 species at 0.5, 1.25 and 2 m on the southern aspect, and the 2023 survey 338 trees at 1.25 m. The 2021 survey included *Acer rubrum*, *A. saccharum*, *Betula alleghaniensis*, *B. lenta*, *B. papyrifera*, *Carya ovata*, *Fagus grandifolia*, *Fraxinus americana*, *Kalmia latifolia*, *Pinus strobus*, *Prunus serotina*, *Quercus alba*, *Q. rubra*, *Q. velutina*, *Sassafras albidum* and *Tsuga canadensis*.

##### Stem chambers

Semi-rigid chambers, used in the 2020–2021 monthly survey, followed Siegenthaler *et al.* (2016). Each was a transparent polyethylene terephthalate sheet framed with 1.5 cm × 3 cm adhesive-backed expanded neoprene strips, with two vertical neoprene wedges holding the sheet at a constant distance from the stem. Chamber volume was $V_c = (HL/(D + 2T)) \times [((D + 2T)/2)^2 - (D/2)^2] - V_{wedges}$, where H is chamber height, L periphery length, D stem diameter and T chamber thickness. Chambers were strapped to stems and gaps sealed with modeling clay tested for $CH_4$ and $CO_2$ off-gassing (Fig. S2a).

Rigid chambers, used from 2021 to 2023, were transparent plastic containers (Rubbermaid) with arcs cut to fit the stem. Their volumes (0.5–2.3 L) were measured by gas-standard dilution in the laboratory, and the enclosed area was taken as the planar area multiplied by the arc length. They were sealed with potting clay (Amaco) and secured with lashing straps. Seals were checked by inspection, by blowing on the seal while watching $CO_2$, and by the linearity of the concentration record (Fig. S2b).

#### Analyzer and protocol

Chambers were connected by 3.7 m (12 ft) of 1/8-inch (3.2 mm) internal-diameter tubing (Bev-a-Line) to the analyzer, which recorded $CO_2$, $CH_4$ and $H_2O$ at 1 Hz (specified $CH_4$ precision < 0.9 ppb; measured field precision in Methods S2). The 5 L $min^{-1}$ flow turned over the chamber volume within about 30 s.

#### Soil

Soil chambers were PVC caps on 25-cm collars inserted 5 cm into the soil at least 24 h before measurement: 12 collars in the upland plots (2 × 6), 6 in the intermediate plots (2 × 3) and 12 in the wetland-margin plots (2 × 6), measured under both dry and saturated conditions. Soil VWC was mapped in December 2020 along transects perpendicular to the moisture gradient, at 79 points about 20 m apart, with a handheld sensor (HydroSense, Campbell Scientific) at 12 cm depth, and was measured at the collars during campaigns (285 measurements).

#### Flux calculation

Flux was $F = (dC/dt)(V_c/A_c)(P/RT)(1 - X_{H_2O})$, where $V_c$ is the total system volume (chamber, 29 $cm^3$ of tubing and the 28 $cm^3$ analyzer cell) and $A_c$ the enclosed surface area. T was the air temperature at the deployment. In the monthly survey (stems and soils) it was the site weather station's hourly mean for the hour containing the start; in the 2021 survey, air temperature measured at the chamber (the station value where there was no reading); and in the 2023 survey, air temperature recorded in the field (the stem temperature where it was missing). These values were also the air temperatures used in the variance partitioning. Fluxes are in nmol $m^{-2}$ $s^{-1}$ for $CH_4$ and µmol $m^{-2}$ $s^{-1}$ for $CO_2$, with P = 101.325 kPa and R = 8.314 L kPa $K^{-1}$ $mol^{-1}$. For each measurement, goFlux's best.flux() chose between linear and Hutchinson–Mosier fits using goodness of fit (MAE and RMSE relative to the analyzer precision of the field day, Methods S2; AICc), physical constraints (a g-factor, the ratio of the non-linear to the linear flux, below 2; $\kappa/\kappa_{max} \leq 1$) and diagnostic flags (regression p-value, comparison with the minimum detectable flux, number of observations). Because stem respiration always produces a positive $CO_2$ flux, the quality of the $CO_2$ fit served as a check on the chamber seal.

#### Internal gas

Holes were sealed with gas-impermeable tape. After 5 min, 20 mL of gas was drawn with gas-tight syringes (Hamilton) into pre-evacuated 12-mL Exetainer vials (Labco) at overpressure. The Yale Analytical and Stable Isotope Center measured $CH_4$ and $CO_2$ by flame ionization detection and $N_2O$ and $O_2$ by electron capture detection. Calibration used certified standards up to 60,340 ppm $CH_4$, with fits weighted to minimize relative error: linear for $CH_4$ up to 5,029 ppm, with interpolation between the two highest standards above that, and quadratic for $CO_2$, $N_2O$ and $O_2$. Predictions were not truncated at zero. No sample fell below the $CH_4$ detection limit (0.18 ppm), and the one sample above the highest standard is reported at that value. Ambient-air blanks were run between samples.

Across 157 trees of 16 species, internal $CH_4$ was positively correlated with internal $CO_2$ ($R^2 = 0.22$, $p < 0.001$) and negatively with internal $O_2$ ($R^2 = 0.10$, $p < 0.001$), and weakly with heartwood *mcrA* ($R^2 = 0.04$, $p = 0.021$) and with surface flux, the relationship strengthening with measurement height ($R^2 = 0.01$ at 50 cm, 0.04 at 125 cm, 0.07 at 200 cm; Fig. S16).

#### Stable isotopes

Isotope precision was <1.15‰ at 10 ppm $CH_4$, and samples below 1.5 ppm $CH_4$ were excluded. The apparent fractionation factor was $\varepsilon_C = (\alpha_C - 1) \times 1000$, with $\alpha_C = (\delta^{13}CO_2 + 1000)/(\delta^{13}CH_4 + 1000)$ (Whiticar, 1999; Conrad, 2005). It was 47‰ when computed within each sample and 53‰ when computed from the dilution-corrected source. Both values lie where the ranges for $CO_2$ reduction and acetate fermentation overlap (Whiticar, 1999), and internal $CO_2$ is mostly respiratory rather than the methanogens' own substrate pool, so $\varepsilon_C$ does not discriminate the pathway. The Keeling-plot intercept was fitted by MM regression (MASS::rlm, method "MM"), which assigns each sample a weight according to its distance from the fitted mixing line. Of 128 samples, 17 received zero weight and 26 a weight below 0.5. Two sensitivity checks, ordinary least squares and exclusion of values more than three interquartile ranges beyond the quartiles (Tukey, 1977), gave:

- ordinary least squares on all samples: −79‰ (95% CI −92 to −66‰);

- after excluding values beyond three interquartile ranges from the quartiles (−151 to +32‰; 117 samples): −69‰ (95% CI −75 to −63‰).

The two-endmember mixing estimate (−70‰) changed by less than 1‰ when the atmospheric endmember was varied from −47 to −35‰. The association between $\delta^{13}CO_2$ and $\delta^{13}CH_4$ was tested by Spearman correlation.

#### Felled black oak

The tree had a basal seam indicating heart rot but a full canopy and was felled on 4 October 2022, before leaf senescence. At each height, heartwood and sapwood cores were taken and a cross-section photographed; leaves, bark, branches, litter, heart rot, roots and soil were also sampled. Tissue samples were taken only from surfaces that had not touched the ground (Arnold *et al.*, 2025), and root, leaf, bark, litter and heart-rot extracts were also treated with a PCR-inhibitor removal kit.

#### Sample collection and DNA extraction

Cores were wrapped in sterile foil, frozen on dry ice and stored at −80 °C, then lyophilized for 72 h and cryo-ground (Spex 6775 Freezer/Mill; 10 min pre-cooling, two cycles of 2 min grinding and 2 min cooling at 10 cycles $s^{-1}$). Equipment was flame-sterilized or bleached between samples (Arnold *et al.*, 2024). Soils were collected at the four cardinal directions around each tree, at a distance equal to its circumference or 1 m, whichever was greater. They were separated into organic and mineral horizons (mineral to 30 cm or refusal), composited by horizon, sieved to 4 mm in the field and frozen on dry ice. Wood was weighed after freeze-drying and soil field-moist, so gene abundances are per gram of dry wood and of fresh soil. Gravimetric water content was not measured on the survey soils; with that of the felled-oak soils (33% mineral, 68% organic), survey soil values per gram of dry mass would be 1.3–1.7 times higher, and the contrast with heartwood remains.

DNA was extracted with the MagMAX Microbiome Ultra Nucleic Acid Isolation Kit on a KingFisher Apex, eluting into 75 µL (wood) or 100 µL (soil). Samples with potential inhibitors were cleaned with the Zymo OneStep-96 PCR Inhibitor Removal Kit.

#### Gene abundance

The probe-based *mcrA* assay (Arnold *et al.*, 2024) used ddPCR Supermix for Probes (No dUTP; Bio-Rad) with 900 nM primers and 250 nM probe. Primers, probe and cycling for all three genes are given in Table S1. Primers were synthesized by the Keck Oligonucleotide Synthesis facility (Yale University) and probes by IDT (PrimeTime).

By species, the two conifers, *Pinus strobus* and *Tsuga canadensis*, carried among the lowest *mcrA* abundances: clearly so in sapwood, where their median was less than a third of the angiosperms' (Wilcoxon $p < 0.001$), and marginally in heartwood ($p = 0.08$), where most species had similar medians and *Acer saccharum* was highest (Fig. S7).

Every plate carried a positive standard. Negative controls were characterized in the method study, with the same assays, sample types and reagent and oligonucleotide lots (Arnold *et al.*, 2024): no-template controls gave no quantifiable signal, none in the probe-based *mcrA* assay and none above background for *pmoA* and *mmoX*. Including extraction losses, the method detects about 500 cells per 100 mg of dry wood (Arnold *et al.*, 2024). Copies are scaled from the eluate to the whole lysate (250 of 800 µL processed; a lower bound, since no transfer exceeded 250 µL). They are not corrected for the remaining extraction losses, about 18% of target DNA spiked onto ground wood, or for losses during freeze-drying and grinding (Arnold *et al.*, 2024), and are therefore conservative; a uniform underestimate does not change species ranks or methanogen:methanotroph ratios. The full protocol is given in Notes S1 (separate file).

#### 16S sequencing

16S libraries were sequenced at the University of Minnesota Genomics Center with dual indexing (Arnold *et al.*, 2025), and sequences with fewer than 10 reads were removed before rarefaction. Bacterial 16S rRNA and fungal ITS copies per µL of extract were quantified by qPCR at the sequencing facility. ITS was converted to copies per gram as for the ddPCR genes, and extracts in which ITS was reported undetected were left out of the ITS analyses.

*Plastid removal and sample dropout.* Removing chloroplast and mitochondrial reads (17.8% of all reads; median 22.7% of a wood library and 2.2% of a soil library) left 36 wood samples (11.8%) below the 3,500-read rarefaction depth. These are well-sequenced libraries dominated by host DNA (median 80% plastid). Because the dropped libraries have, by definition, fewer than 3,500 non-organelle reads, dropped and retained wood were compared at an equal depth of 1,000 reads; they did not differ in the abundance or detection of methanogens or methanotrophs (34 dropped, 259 retained; smallest $q = 0.42$). Within heartwood, however, none of the 11 dropped samples contained methanogens, where 5.0 were expected from retained samples of the same species at the same depth ($q = 0.0014$). The 16S estimate of methanogen prevalence in heartwood is therefore slightly biased upward. Heartwood methanogen prevalence by ddPCR (97%) is unaffected.

### Methods S2. Detection limits and apparent stem uptake

#### Instrument precision

Analyzer precision (σ) was estimated from the second differences of each closure's concentration record and taken as the median per field day; one analyzer made every measurement. A second difference cancels any linear trend, so the estimate does not depend on the flux being measured. The median daily σ was 1.08 ppb in the 2021 height and molecular survey (range 0.92–1.37 ppb over 18 days), 1.59 ppb in the 2023 cross-species survey (1.44–1.78 ppb, 19 days) and 2.02 ppb in the monthly survey (1.44–2.42 ppb, 34 days).

#### Detection limits and status

A fit was treated as noisy when its residual scatter exceeded σ, and the curvature of the non-linear fit was bounded by the minimum detectable flux (Hüppi *et al.*, 2018). The minimum detectable flux (MDF) of each closure was 1.96σ/t multiplied by the flux term, where t is the closure time in seconds (from the record's own timestamps) and the flux term the chamber geometry and temperature factor (Parkin *et al.*, 2012); 1.96 is used as a conventional benchmark rather than a calibrated 95% test (Cowan *et al.*, 2025). This form does not credit the averaging gained by fitting a slope to many readings, so it is conservative. The regression test (slope $p < 0.05$) is reported alongside. Median stem MDF was 0.007 nmol $m^{-2}$ $s^{-1}$ in the 2021 height and molecular survey, 0.023 in the 2023 cross-species survey and 0.007 in the monthly survey (soil, 0.097).

Of 1,191 stem measurements, 824 (69.2%) exceeded their MDF: 748 emissions and 76 uptakes. Of 286 soil measurements, 259 (90.6%) were detected, 224 of them uptakes. By the regression test, 1,072 stem measurements (90.0%) had a slope distinguishable from zero. The median detected stem uptake (−0.024 nmol $m^{-2}$ $s^{-1}$) is 2.2% of the median detected soil uptake (−1.06 nmol $m^{-2}$ $s^{-1}$).

#### Screening criteria

Common screening criteria differ up to 26-fold in what they retain:

| Criterion | Stem measurements retained | Retained (%) | Retained that are uptake (%) | Change in mean flux |
|---|---|---|---|---|
| Retain all | 1,191 | 100 | 18.5 | – |
| Slope $p < 0.05$ | 1,072 | 90.0 | 15.2 | +11% |
| Above MDF | 824 | 69.2 | 9.2 | +44% |
| $r^2 > 0.7$ | 458 | 38.5 | 1.3 | +137% |

| Criterion | Stem measurements retained | Retained (%) | Retained that are uptake (%) | Change in mean flux |
|---|---|---|---|---|
| $r^2 > 0.9$ | 270 | 22.7 | 0.7 | +251% |

An $r^2$ threshold rewards a smooth fit regardless of the size of the flux. Applied here, it removes nearly all apparent uptake and more than doubles the mean. We therefore retained every measurement and report detection status separately. The only exclusions were for procedural failure, when a chamber was not sampling ambient air at closure. One screen was applied to the soil and 2021-survey deployments, the only ones with a recorded starting concentration: deployments whose starting $CH_4$ exceeded 1.5 times the campaign median were flagged and excluded from model training. It removed two soil deployments from one session and no stem deployment.

#### Apparent stem uptake

Four observations indicate that apparent stem uptake reflects measurement resolution rather than a sink. First, its prevalence follows instrument precision: detected uptake made up 1.1% of measurements in the quietest survey (the 2021 survey; deepest −0.026 nmol $m^{-2}$ $s^{-1}$), against 10.4% and 8.7% in the two noisier surveys. Second, in the cross-species survey, apparent uptake tracked how far a chamber's starting concentration sat above local ambient (the 10th percentile of the trace in the 3 min either side of the enclosure). Detected uptakes opened a median 27 ppb above local ambient, against 7 ppb for both undetected measurements and detected emissions (uptake against the rest, Wilcoxon $p < 0.001$; emissions against undetected, $p = 0.75$; 337 measurements). That is the pattern expected when an enclosure that opens above ambient decays toward it. Third, the most negative stem measurement (−0.38 nmol $m^{-2}$ $s^{-1}$) opened 152 ppb above local ambient, after a $CH_4$ excursion to 3.8 ppm (1.9 times ambient) in the preceding 15 min. Fourth, the only measurements above 2 m, on the climbed black oak at 4, 6, 8 and 10 m, were all positive.

The linear-into-uptake form in the scaling grid (Methods S6) is therefore included only as a lower bound; the climbed-tree profile does not support it.

#### Moss and lichen cover

In the 2023 survey, moss and lichen cover within each chamber footprint was assessed visually on a three-point scale of relative cover (1, least; 3, most); taxa and functional groups were not identified, and their contribution to stem flux was not isolated. Of 330 live-stem measurements, 278 had a score (113 scored 1, 117 scored 2, 48 scored 3). Across species, flux rose slightly with cover (Spearman $\rho = 0.15$, $p = 0.010$), because species with more cover, such as *B. alleghaniensis*, also emitted more. Within species, cover was unrelated to flux (linear model of arcsinh-transformed flux on species and score, p = 0.47), and no species with at least ten scored trees showed significantly lower flux with more cover.

##### Bark loss and wounding

In the 2023 survey, bark loss and wounding were each scored visually at each chamber as healthy, moderate or severe (1–3), or dead stem; in the models the dead class was scored 4 (Fig. 7a; Fig. S18). Of 338 measurements, 282 had a bark-loss score and 282 a wounding score. Flux against each score, and wood *mcrA* against ITS load (Fig. 7b), were fitted by linear and quadratic mixed models with species as a random intercept and compared by AIC.

#### Methods S3. Methanotrophs at several taxonomic resolutions and phylogenetic placement

To test how much the genus-level rule under- or over-counts methanotrophs, their share of each sample's 16S community (libraries of at least 1,000 reads, not rarefied) was computed at order, family and genus level and with phylogenetic placement, and each grouping was correlated with ddPCR *pmoA* and *mmoX* in the same extracts (Spearman, all samples; Table S4).

*Tiers by compartment.* Of 733 methanotroph ASVs, 77 were Known and 656 Putative. Known methanotrophs were substantial only in mineral soil (1.00% of the community), almost all *Methylocapsa*, a genus that includes the atmospheric-methane oxidizer *M. gorgona* (Tveit *et al.*, 2019); they made up 0.02–0.06% of heartwood, sapwood and organic soil communities. Adding Putative ASVs raises heartwood to 0.55%, sapwood to 1.29%, organic soil to 1.23% and mineral soil to 1.41%. In wood, the three most abundant resolved Beijerinckiaceae genera with no annotated methane monooxygenase (Table S3) were each more abundant than all Known methanotrophs together (*Lichenibacterium* 0.7% of heartwood and 2.4% of sapwood communities; *Methylobacterium–Methylorubrum* 0.3% and 0.5%).

In organic soil the Putative increment is Methylacidiphilaceae. Genus-unresolved Methylacidiphilaceae are Putative under the same rule as other mixed families, but the bound is loose for them: methane monooxygenase is confirmed only in *Methylacidiphilum* and *Methylacidimicrobium*, acidophiles from geothermal and volcanic habitats (van Teeseling *et al.*, 2014); mesophilic relatives from peat (Dedysh *et al.*, 2021) and related lineages of the order from tree bark (Leung *et al.*, 2026) lack it; and no ASV here resolves to either genus. The *pmoA* primers do not target verrucomicrobial *pmoA* (Knief, 2015), so these taxa do not contribute to the ddPCR methanotroph measures.

*Phylogenetic placement.* Placement used the ASV tree from sequence processing. For each Beijerinckiaceae ASV unresolved at genus level (all in the Putative tier), the nearest genus-resolved Beijerinckiaceae ASV was found by patristic distance, and the ASV was counted as placed with methanotrophs if that neighbor belonged to a methanotroph genus (*Methylocapsa*, *Methylocella* or *Methyloferula*) within 0.03 substitutions per site, about 97% identity over V4. Of 273 such ASVs, 74 were placed with methanotroph genera, 100 with genera that lack methane monooxygenase, 59 with other genera, and 40 had no genus-resolved relative within the threshold. Thresholds of 0.02 and 0.04 placed 59 and 91 ASVs, changing the Known + tree-placed share of sapwood from 0.11% to 0.20% and leaving soils almost unchanged (Table S4). The felled oak's 16S data are not on the ASV tree, so its tissues are summarized at order, family and genus level only. Placed ASVs make up 9–14% of the wood Putative signal (0.08% of heartwood and 0.12% of sapwood communities), three to five times the Known share.

*Genes by compartment.* In heartwood *pmoA* and *mmoX* were present at similar medians ($2.4 \times 10^4$ and $2.5 \times 10^4$ copies $g^{-1}$ dry wood); *pmoA* exceeded *mmoX* about threefold in sapwood, and in soils by roughly two orders of magnitude ($3.3–3.9 \times 10^6$ against $3.0–4.1 \times 10^4$ copies $g^{-1}$ fresh soil) (Fig. S8a). *pmoA* was detected in 95.2% of heartwood, 99.4% of sapwood and 100% of soil samples; *mmoX* in 93.2% of heartwood, 97.4% of sapwood and 100% of soil samples.

*Comparison with ddPCR.* At 3,500 reads, 16S misses a taxon below about 0.03% of a sample; the *pmoA* primers also miss NC10 *pmoA* and under-detect the USCα lineage (Knief, 2015). The *pmoA* primers exclude ammonia monooxygenase, and *Methylocella*, which lacks pMMO, is detected by *mmoX* only. Organic and mineral soil carried *pmoA* at similar density by ddPCR (median ratio 0.84) and in the same proportion to total bacterial 16S, although 16S assigned Known methanotrophs almost only to mineral soil. In sapwood, 16S methanotroph groups were at most weakly related to *pmoA* (Spearman $\rho$ 0.06–0.29 across group definitions; Table S4) and not to *mmoX*; in soils, 16S methanotroph groups were negatively related to the genes, mainly through Methylacidiphilaceae and, in mineral soil, possibly USCα (Table S4).

*Defining tree methanotrophs.* Accounts of tree methanotrophs depend heavily on how methanotrophs are defined. The choice matters: in our wood, Beijerinckiaceae genera without the enzyme outnumbered Known methanotrophs 70–100-fold. Surveys of tree tissues have counted methylotrophs such as *Methylobacterium–Methylorubrum*, which may lack methane monooxygenase (Dedysh *et al.*, 2004; Semrau *et al.*, 2011) but dominated the methane-oxidizer signal of leaves and bark (Moisan *et al.*, 2026), and bark Methylacidiphilaceae have been counted as methanotrophs from 16S family membership (Jeffrey *et al.*, 2021), although genomes of related lineages of the same order from tree bark lack methane monooxygenase (Leung *et al.*, 2026) (Table S7). A strict genus rule errs the other way, missing methanotrophs that 16S cannot name, such as the *pmoA* carriers of organic soil and the tree-placed Putative ASVs. Metagenomes from conifers contained few canonical methanotroph genes and instead monooxygenases related to those of propane and butane oxidizers, whose role in $CH_4$ oxidation is unknown (Putkinen *et al.*, 2021). Lifestyles are as uncertain as identities. *Methylocella*, the Known methanotroph of the felled oak's bark and canopy, and some *Methylocapsa* and *Methylocystis* also grow on multicarbon substrates (Dedysh *et al.*, 2005; Semrau *et al.*, 2011), and the methanotrophs isolated from conifer needles also grew on methanol, which plant tissues release (Putkinen *et al.*, 2021). Which of these organisms oxidize $CH_4$ in situ, at what concentrations, and whether any oxidize atmospheric $CH_4$ on woody surfaces (Gauci *et al.*, 2024) can be settled only by activity measurements (rates, transcripts, stable-isotope probing), not by census.

#### Methods S4. Gene–flux relationships at the tree and species levels

##### Species, environment and natural history

Species identity and its interactions with environment explained more of the among-tree variance in flux than environment alone (Fig. 3b), and natural history matters as much as taxonomy or site. The two birches had the highest and lowest mean fluxes (Fig. 3a). *B. alleghaniensis* occupied the wettest soils of any species (Fig. S6), where soil methanogens were most abundant. *B. lenta* is regenerating vigorously in this stand: 84% of its 876 inventoried stems are under 10 cm DBH. Across southern New England it is replacing hemlock killed by the hemlock woolly adelgid (Orwig & Foster, 1998; Kizlinski *et al.*, 2002) and filling gaps left by declining oak (McEwan *et al.*, 2011), so its canopy trees are likely younger than co-occurring trees of similar size and may carry less of the heart rot that supports methanogens; the trees we measured were of ordinary size for the survey (median DBH 29 cm). Neither taxonomy, with two congeners at the extremes, nor environment alone (1.9% of among-tree variance) predicts a species' flux, which is why we examined plant traits and life history (Methods S5).

##### Area weighting

Heartwood and sapwood abundances (copies $g^{-1}$ dry wood) were combined into one value per tree, the mean over the breast-height cross-section of a radial profile that takes the sapwood value in the outer 5 cm (the outer core segment), the heartwood value within 5 cm of the pith, and changes linearly with radius between them:

$C = C_s + (C_h - C_s)(r_1^2 + r_1 r_2 + r_2^2)/(3R^2)$,

where $C_h$ and $C_s$ are the heartwood and sapwood abundances, $R = DBH/2$, $r_1 = \min(5\ \text{cm}, R - 5\ \text{cm})$ and $r_2 = \max(R - 5\ \text{cm}, r_1)$. In trees with $R \leq 10$ cm the heartwood value applies only inside the outer 5 cm ($r_1 = r_2 = R - 5$ cm), and a tree with $R \leq 5$ cm takes the sapwood value. Trees without a measured diameter have no area-weighted value. With logistic transitions (moderate or steep) or steps in place of the linear transition, including the two bounds (the heartwood value everywhere inside the outer 5 cm, or only within $r_1$ of the pith), the species-level ratio $R^2$ ranged 0.44–0.47 ($p \leq 0.036$), *mcrA* $R^2$ 0.29–0.31 and *pmoA* + *mmoX* $R^2$ 0.18–0.21, and within species only *mmoX* remained associated with flux.

#### Tree level: linear models

Gene–flux models used the 125 trees with *mcrA*, *pmoA*, *mmoX* and flux all measured. Each gene was added to a linear model that already contained species, so its coefficient measures the association within species. Adding *mcrA*, *pmoA*, *pmoA* + *mmoX*, or the methanogen:methanotroph ratio did not improve on species alone (gene $p = 0.31$, 0.39, 0.32 and 0.47). Area-weighted *mmoX* was the only gene associated with flux within species: species plus *mmoX* explained 39.2% of variance (adjusted $R^2 = 0.308$; AIC = 7.0), 3.7 percentage points more than species alone ($\beta = -0.318$, SE = 0.123, $p = 0.011$).

With tissue-specific gene abundances in place of area-weighted ones (129 trees), the full model (species and six tissue-specific gene terms) explained 40.5% (adjusted $R^2 = 0.295$; AIC = 9.7), and only sapwood *mmoX* was significant ($\beta = -0.271$, SE = 0.097, $p = 0.006$; all other gene terms $p \geq 0.07$; species $F = 4.79$, $p < 0.001$). Soil gene abundances added little beyond tree *mmoX* ($\Delta R^2 \leq 0.02$; all $p \geq 0.12$). A negative within-species association of sapwood *mmoX* with flux is consistent with consumption capacity in the sapwood, but it is one significant term among many tested.

#### Tree level: random forests

To allow for non-linear and interacting gene effects, random forests were compared on the 123 trees with complete gene, flux, diameter and soil-moisture data. The response was tree-mean flux on the arcsinh scale, and skill was measured on held-out trees (30 repeats of 5-fold cross-validation, with the same folds for every model). Genes alone (log *mcrA*, *pmoA* and *mmoX*) predicted held-out trees no better than their mean ($R^2 = -0.01 \pm 0.04$). Species alone gave $R^2 = 0.20$, and species with diameter and soil moisture 0.18; adding the three genes left it at 0.18 (higher in about half of the repeats). Soil moisture and species carried most of the permutation importance.

#### Species level

Species-level relationships used the ten species with at least five trees. Species medians of area-weighted gene abundance were regressed against median flux (Fig. 8). Area-weighted *mcrA* was positively related to flux ($R^2$ = 0.299, r = 0.546, p = 0.10), *pmoA* + *mmoX* negatively ($R^2$ = 0.201, r = −0.448, p = 0.19), and *pmoA* and *mmoX* separately only weakly ($R^2$ = 0.24, p = 0.15; $R^2$ = 0.03, p = 0.61). The methanogen:methanotroph ratio was the strongest predictor ($R^2$ = 0.455, r = 0.675 on the log ratio, p = 0.032) and was preferred over *mcrA* alone (AIC −44.5 against −41.9). Heartwood *mcrA* concentration without area weighting gave $R^2$ = 0.391 (p = 0.053).

Fig. S21 shows how these relationships change with the scale of aggregation. Gene–flux associations are negligible between individual trees, appear weakly when a species-level gene value is used to predict individual fluxes, and strengthen for species medians and means. In that figure flux is arcsinh-transformed before aggregation, so that a species mean is not dominated by a few large fluxes. On that scale the ratio is significant for both species medians and means, *mcrA* for means (and marginally for medians), and the methanotroph genes for neither. Aggregation and ratios both carry statistical risks (Jasienski & Bazzaz, 1999; Bradford *et al.*, 2017); the species pattern holds in direction for the numerator and denominator separately (Fig. 8d–e), and methanogen and methanotroph abundances are uncorrelated (Fig. S23).

### Methods S5. Plant traits

We compiled per-species traits from public databases and each species' realized soil-moisture niche, the mean volumetric water content across its measured stems at our site (Fig. S6). Wood and bark density are species means from the Global Wood Density Database v2.2 (Fischer *et al.*, 2026). Wood and bark carbon, wood C:N, bark volume, coarse-wood decomposition rate, waterlogging tolerance, rooting depth, fine-root diameter and tissue density, mycorrhizal type and maximum longevity are from the TRY Plant Trait Database (Kattge *et al.*, 2020), from the datasets listed in Table S8. TRY records flagged as duplicates or probable errors (error risk > 4) and values republished from another dataset were removed; each trait was restricted to one quantity (maximum rooting depth, maximum age); species values are means of dataset means. Maximum age serves as a ranking of life-history pace, from short- to long-lived species, not as a typical age, and the analyses are rank-based (Spearman); longevity taken as the mean of all records, or of records reported as means, gave the same reported associations. The trait table and the source of every value are archived with the data (Data Availability). Wood pH was not used: published species-level values exist for fewer than eight of the ten species.

Traits were included by rules set before analysis: data for at least eight of the ten species (rooting depth, with six, was excluded); at least five distinct values, or at least three species in the smallest class of a categorical trait (mycorrhizal type, scored ectomycorrhizal or arbuscular by the majority of records); one column per property (duplicates from other sources and ratios of included traits excluded); values plausible for adult trees (TRY plant height, which mixes life stages, was excluded); and a stated route by which the trait could affect stem methane cycling. Thirteen traits in five groups met these rules (Fig. S24).

Each was related to species-level flux and to area-weighted *mcrA*, *pmoA* + *mmoX* and methanogen:methanotroph abundance by Spearman correlation, for the ten species with at least five sampled trees and five flux measurements. Because the two conifers are extremes in trait space, each correlation was repeated among the eight broadleaf species alone. An association is reported when it holds ($p < 0.05$, same sign) in both analyses, or when it holds among broadleaf species, has the same sign across all ten, and the two conifers differ from each other in the same direction as the broadleaf trend, recognizing that the two analyses share eight species and so are not independent. No association passed Benjamini–Hochberg correction across the 52 tests (minimum $q = 0.32$ across all ten species, 0.36 among broadleaf species), as expected for effects of this size with ten species; the associations are exploratory.

Bark density and the ratio also held when any single species was left out (largest $p = 0.028$). A joint multivariate analysis (redundancy analysis on trait principal components; partial least squares judged by leave-one-out cross-validation) found no structure that was robust to the choice of traits or species, and no trait or combination predicted flux.

### Methods S6. Random forests and stand-scale bounds

#### Training data and predictors

*Training population.* Of the 1,191 stem deployments in the three survey campaigns, 64 on 16 standing dead stems and one root-crown measurement were excluded, leaving 1,126 measurements from 488 live trees. Dead stems were excluded because the inventory to which the model is applied contains none; they remain in every descriptive analysis.

*Responses and quality control.* Both models were fitted to flux on its measured scale (nmol $m^{-2}$ $s^{-1}$), without transformation, so stand totals need no back-transformation. No measurement was removed for its flux value or detection status. The two soil deployments that failed the chamber-closure screen were excluded (Methods S2), and the soil model was not trained on the 20 measurements from two collars in open wetland outside the censused stand (no stem within 10 m), because it is applied only within the stand; these two collars produced all five soil fluxes above 5 nmol $m^{-2}$ $s^{-1}$.

Differences between chamber types were removed from the flux measurements before modeling, because inventory stems have no chamber type.

*Seasonal coverage.* The training data are weighted toward summer: June–August hold 881 of the 1,126 stem measurements, January has no stem or soil measurements, and no stem measurement in December has an observed soil temperature. The drivers are monthly climatologies rather than campaign values, so outside the summer the model interpolates between sparse months.

*Predictors.* The stem model used species, diameter, soil moisture at the tree, soil temperature and measurement height. Measurement height entered the model for 431 of the 435 training measurements from the 2021 multi-height survey; all others were taken at breast height and assigned 125 cm, as were the four 2021 *Kalmia latifolia* deployments made at 75 cm. Diameter was not measured on six of the 488 training trees (18 measurements), which enter the model with diameter missing. The soil model used soil moisture, soil temperature and month. Training used the soil temperature and moisture measured at each deployment. For prediction, the drivers are monthly climatologies for 2019–2022 from the site weather station; soil temperature comes from the station's air temperature over the preceding 30 days, through a relation fitted to the soil readings. Soil moisture at a location is the product of a monthly plot-wide level (a climatology) and a relative spatial pattern: a thin-plate spline through the 71 points of the December survey, normalized to a mean of one over the stand. Every located stem takes its value from this surface: 432 of the 8,160 located stems (5.3%; 8.7% of basal area) lie outside the area spanned by the survey points, where the spline extrapolates up to 34 m from the nearest point, and the 34 stems without coordinates take the stand mean. The pattern supports only large-scale wet–dry structure (leave-one-out RMSE about 17 percentage points against a survey mean of 22.6%).

*Daytime measurements.* Fluxes were measured in daytime, and the budget assumes that daytime chamber flux represents the daily mean; there were no night measurements. Adding air temperature as a predictor raised grouped $R^2$ to 0.13 for stems and 0.43 for soils, but made the soil sink depend on whether daytime or 24-h air temperature drove the scaling (482 or 519 mg $CH_4$ $m^{-2}$ $yr^{-1}$); without it the sink is 440 mg. Driving the models with daily instead of monthly values for 2019–2022 gave 0.95 times the soil sink and 1.02 times the stem flux (0–2 m band, 300 sampled stems).

#### Skill and calibration

Both random forests used ranger (Wright & Ziegler, 2017) with 10,000 trees, a minimum node size of 5 and $\sqrt{p}$ predictors tried at each split; tuning these settings, with selection nested inside cross-validation, changed grouped $R^2$ by less than 0.02. Skill was assessed by grouped cross-validation (100 repeats of 5 folds) with whole trees or soil collars held out, since a tree contributes up to 13 rows. Grouped $R^2$ was 0.10 (SD 0.04) for stems and 0.39 (SD 0.03) for soils, against out-of-bag 0.23 and 0.41; Fig. S25 also gives the concordance correlation coefficient (Lin, 1989). Under grouped cross-validation, held-out predictions averaged about 5% above observations (mean predicted/observed 1.05 across the 1,126 stem training measurements, unweighted, on the measured flux scale); this checks sample-level bias, not the inventory-weighted stand budget, which uses the per-species calibration below. The chamber-type offset was estimated once from all stem data and was not re-estimated within folds. Importance was measured by permutation, which, unlike impurity gain, is not biased toward many-valued predictors. Soil moisture carried 94% of the soil model's permutation importance. In the stem model soil moisture (about 32%) and species (about 28%) contributed most, measurement height (recorded for 431 measurements, the rest taken at breast height) and diameter about 13–17% each, and soil temperature about 10%; the ordering varied with the importance estimator.

*Calibration.* Because random-forest predictions shrink toward the mean, which biases a sum, stand totals apply a per-species multiplicative calibration, the ratio of observed to out-of-bag predicted mean flux, for species with at least five training measurements; species with fewer, or with a non-positive ratio, are left uncalibrated (ratio 1) (Fig. S26). Across reasonable calibration choices the measured-band total ranges from 5.1 to 5.5 mg $CH_4$ $m^{-2}$ $yr^{-1}$.

#### Stand geometry and the measured band

The censused stand is 3.72 ha (a 4-ha square less seven uncensused 20-m quadrats) with 8,194 stems (172.5 $m^2$ basal area). Each stem's lateral surface between 0 and 2 m was taken as $\pi \cdot DBH \cdot 2$ m ($\pi \cdot DBH \cdot 0.75$ m for the shrub *Kalmia latifolia*), giving 4,482 $m^2$ of bark, 12.0% of the stand area. Each stem's flux in that band is the calibrated model prediction for its species, size, moisture and month, evaluated at four heights within the band; below 0.5 m the 0.5 m value is held. The model predicts higher flux below about 0.9 m than at breast height in every species; holding each stem's breast-height prediction across the band instead gives 3.0 rather than 5.5 mg $CH_4$ $m^{-2}$ $yr^{-1}$. Soil flux was predicted on a grid across the stand and scaled by the soil fraction of the area.

#### Above 2 m: the scaling grid

Tree heights were estimated as $H = 1.37 + a \cdot DBH^b$ with exponents from Hulshof *et al.* (2015) ($b = 0.53$ for angiosperms, 0.60 for gymnosperms) and a set so that a stem at the 95th diameter percentile reaches the observed canopy height (25 m). Woody area index took five values: 1.50 (Whittaker &

Woodwell, 1967), 3.07 (Gauci *et al.*, 2024), and 2.32, 2.93 and 3.53 from this stand's inventory, using Whittaker and Woodwell's conic stem surface, their taper correction (1.20–1.35) and branch-to-stem ratios (2.7–4.0). Woody area was split into bole and branch components. The bole was a cone or a paraboloid, and branch area was placed uniformly over the stem, uniformly over its upper half, or as a Gaussian centered at 50% or 75% of height (Fig. S27b).

Six forms of flux above 2 m were applied to each stem's own 2-m flux (Fig. S27a): constant; exponential, power and linear declines fitted to the campaign means at the three heights, with the linear decline floored at zero; a linear decline continuing through zero into net uptake, bounded at the median detected stem uptake ($-0.024$ nmol $m^{-2}$ $s^{-1}$; Methods S2); and a per-stem exponential decay fitted to each stem's modeled 0–2 m profile. A random forest returns a constant above its training range, so the model itself cannot supply a shape above 2 m. All forms hold flux constant or declining above 2 m (per-stem increases were capped), so we consider the set conservative: a mid-stem maximum like the felled oak's (Fig. 7e) would raise whole-surface emission.

*Choosing among the forms.* On the trees measured at all three heights, each form was fitted to two heights and used to predict the third (Fig. S27d). For upward extrapolation (0.5 and 1.25 m predicting 2 m), a constant was most accurate (RMSE 0.41), followed by linear (0.69), power (1.02) and exponential decay (6.69). The constant beat the linear form (paired Wilcoxon on absolute error, $p = 0.0001$) but not decisively the exponential ($p = 0.11$). Within the band, a power law interpolated 1.25 m best (RMSE 0.15). Pooled across trees, flux declined within the lowest 2 m, a decline carried by *B. alleghaniensis* (without it the pooled slope was near zero); for extrapolating upward, a constant was most accurate. This cross-validation tests 0.75 m of extrapolation, whereas the scaling projects more than 20 m.

*Results across the grid.* The grid and the leverage of each assumption are shown in Fig. S28. Flux form dominates: the median whole-surface total ranges over 93 mg $CH_4$ $m^{-2}$ $yr^{-1}$ across flux forms, against 10 mg across woody area index, 7 mg across branch placement and less than 1 mg across bole shape. The stand is a net sink in all 240 combinations.

*The two cases quoted in the text.* At the central bottom-up woody area index (2.93), Gaussian-75 branch placement and a conic bole, exponential decay fitted to the campaign means gives 10.4 mg $CH_4$ $m^{-2}$ $yr^{-1}$ (2.4% of soil uptake; 48% extrapolated above 2 m), and constant flux gives 84 mg (19%; 94% extrapolated; Fig. S27c). Both shapes come from the measured heights. The constant extrapolated upward best in cross-validation and is consistent with the only measurements above 2 m (Fig. 7e); the exponential, which carries the pooled decline upward, extrapolated worst. The per-stem exponential form stays in the grid but is not quoted, because its decline comes from the model rather than the measurements: it is fitted to each stem's modeled 0–2 m profile over 1.13 m and projected to about 25 m, and without the modeled values below 88 cm the median fitted slope changes sign, from −0.53 to +0.11 per m.

#### Methods S7. Inferred functions (FAPROTAX and PICRUSt2)

FAPROTAX assigns functions from genus names (Fig. 6b; Fig. S11). It annotated 47% of heartwood and 51% of sapwood abundance (32% of ASVs resolved to genus). Its percentages are therefore lower bounds on shares of the whole community and, because categories overlap, not additive (Louca *et al.*, 2016). Several categories also recount the methanogens: FAPROTAX lists Methanobacteriaceae under dark hydrogen oxidation and Methanomassiliicoccaceae under methylotrophy. With methanogen ASVs removed, fermentation (16.2% of heartwood against 6.7% of sapwood) and anaerobic chemoheterotrophy (13.7% against 7.1%) remained enriched in heartwood, whereas dark hydrogen oxidation did not differ (1.05% against 1.08%), and methylotrophy, methanotrophy and aerobic chemoheterotrophy were enriched in sapwood (each FDR ≤ 0.003; dark hydrogen oxidation FDR = 0.32). These shares were computed per library and compared between heartwood and sapwood of the same tree (paired Wilcoxon signed-rank tests, 108 trees, replicate libraries averaged, Benjamini–Hochberg correction across the functions of Fig. 6b). Tissue-level FAPROTAX predictions for this forest were reported by Arnold *et al.* (2025); Fig. S11 extends them to species.

Families (Fig. 6a; Figs S9, S10) and PICRUSt2 pathways (Fig. 6c; Figs S12–S14) were tested one at a time in wood samples with at least 100 16S copies $\mu L^{-1}$ of extract, by linear mixed-effects models of gene concentration in the extract (*mcrA* or *pmoA*) on the family's or pathway's relative abundance, compartment (heartwood or sapwood) and $\log_{10}(1 + 16S \text{ copies } \mu L^{-1})$, with tree as a random intercept; gene and relative abundance were standardized. Each family or pathway present in more than 20 samples was tested by a likelihood-ratio test against the model without it, with Benjamini–Hochberg correction across families or pathways.

Two families besides the methanogens were associated with *mcrA* after Benjamini–Hochberg correction (Fig. 6a; Fig. S9), and the evidence for a role in methanogenesis differs between them. Dysgonomonadaceae are anaerobic fermenters of plant polymers (Duan *et al.*, 2016) described from methanogenic biogas reactors (Hahnke *et al.*, 2016), where *Proteiniphilum* is among the dominant acetate users (Feng *et al.*, 2023). Eggerthellaceae ferment amino acids (Noecker *et al.*, 2023) and have no established role in supplying $H_2$ to methanogens, so their co-occurrence may reflect a shared anaerobic niche rather than syntrophy. Two $H_2$-producing fermenters were associated nominally but not after correction ($p < 0.01$; FDR ≥ 0.11; Fig.

6a). Christensenellaceae (Morotomi *et al.*, 2012) support hydrogenotrophic methanogens through interspecies $H_2$ transfer in co-culture (Ruaud *et al.*, 2020) and co-occur with *Methanobrevibacter* in the human gut (Goodrich *et al.*, 2014; Waters & Ley, 2019). *Ethanoligenens*, the type genus of Ethanoligenenaceae, ferments glucose to acetate, ethanol, $H_2$ and $CO_2$ (Xing *et al.*, 2006).

PICRUSt2 pathways were tested against *mcrA* after excluding contributions from methanogen-classified ASVs (Fig. 6c; Fig. S12; without that filter, Fig. S13; *pmoA*, Fig. S14). Samples with more *mcrA* were enriched in C1 assimilation and carbon fixation (the RuMP cycle, the reductive acetyl coenzyme A pathway, the Calvin–Benson–Bassham cycle), fermentation and carbohydrate breakdown, and cell-wall synthesis; samples with less (predominantly sapwood) in the TCA cycle, aerobic respiration, heme biosynthesis and assimilatory sulfate and nitrate metabolism.

**Arnold W, Gewirtzman J, Raymond PA, Bradford MA, Butler C, Peccia J**. **2024**. A method for sampling the living wood microbiome. *Methods in Ecology and Evolution* **15**: 1084–1096.

**Arnold W, Gewirtzman J, Raymond PA, Duguid MC, Brodersen CR, Brown C, Norbraten N, Wood QTV, Bradford MA, Peccia J**. **2025**. A diverse and distinct microbiome inside living trees. *Nature* **644**: 1039–1048.

**Bradford MA, Veen GF, Bonis A, Bradford EM, Classen AT, Cornelissen JHC, Crowther TW, De Long JR, Freschet GT, Kardol P, *et al.*** **2017**. A test of the hierarchical model of litter decomposition. *Nature Ecology & Evolution* **1**: 1836–1845.

**Conrad R**. **2005**. Quantification of methanogenic pathways using stable carbon isotopic signatures: A review and a proposal. *Organic Geochemistry* **36**: 739–752.

**Cowan N, Levy P, Tigli M, Toteva G, Drewer J**. **2025**. Characterisation of analytical uncertainty in chamber soil flux measurements. *European Journal of Soil Science* **76**: e70104.

**Dedysh SN, Beletsky AV, Ivanova AA, Danilova OV, Begmatov S, Kulichevskaya IS, Mardanov AV, Ravin NV**. **2021**. Peat-inhabiting Verrucomicrobia of the order Methylacidiphilales do not possess methanotrophic capabilities. *Microorganisms* **9**: 2566.

**Dedysh SN, Dunfield PF, Trotsenko YA**. **2004**. Methane utilization by *Methylobacterium* species: New evidence but still no proof for an old controversy. *International Journal of Systematic and Evolutionary Microbiology* **54**: 1919–1920.

**Dedysh SN, Knief C, Dunfield PF**. **2005**. *Methylocella* species are facultatively methanotrophic. *Journal of Bacteriology* **187**: 4665–4670.

**Duan J, Liang J, Wang Y, Du W, Wang D**. **2016**. Kraft lignin biodegradation by *Dysgonomonas* sp. WJDL-Y1, a new anaerobic bacterial strain isolated from sludge of a pulp and paper mill. *Journal of Microbiology and Biotechnology* **26**: 1765–1773.

**Feng G, Zeng Y, Wang H-Z, Chen Y-T, Tang Y-Q**. **2023**. *Proteiniphilum* and *Methanothrix harundinacea* became dominant acetate utilizers in a methanogenic reactor operated under strong ammonia stress. *Frontiers in Microbiology* **13**: 1098814.

**Fischer FJ, Chave J, Zanne A, Jucker T, Fajardo A, Fayolle A, Lima RAF de, Vieilledent G, Beeckman H, Hubau W, *et al.*** **2026**. Beyond species means – the intraspecific contribution to global wood density variation. *New Phytologist* **249**: 2630–2651.

**Gauci V, Pangala SR, Shenkin A, Barba J, Bastviken D, Figueiredo V, Gomez C, Enrich-Prast A, Sayer E, Stauffer T, *et al.*** **2024**. Global atmospheric methane uptake by upland tree woody surfaces. *Nature* **631**: 796–800.

**Goodrich JK, Waters JL, Poole AC, Sutter JL, Koren O, Blekhman R, Beaumont M, Van Treuren W, Knight R, Bell JT, *et al.*** **2014**. Human genetics shape the gut microbiome. *Cell* **159**: 789–799.

**Hahnke S, Langer T, Koeck DE, Klocke M**. **2016**. Description of *Proteiniphilum saccharofermentans* sp. nov., *Petrimonas mucosa* sp. nov. and *Fermentimonas caenicola* gen. nov., sp. nov., isolated from mesophilic laboratory-scale biogas reactors, and emended description of the genus *Proteiniphilum*. *International Journal of Systematic and Evolutionary Microbiology* **66**: 1466–1475.

**Hulshof CM, Swenson NG, Weiser MD**. **2015**. Tree height–diameter allometry across the United States. *Ecology and Evolution* **5**: 1193–1204.

**Hüppi R, Felber R, Krauss M, Six J, Leifeld J, Fuß R**. **2018**. Restricting the nonlinearity parameter in soil greenhouse gas flux calculation for more reliable flux estimates. *PLoS ONE* **13**: e0200876.

**Jasienski M, Bazzaz FA**. **1999**. The fallacy of ratios and the testability of models in biology. *Oikos* **84**: 321.

**Jeffrey LC, Maher DT, Chiri E, Leung PM, Nauer PA, Arndt SK, Tait DR, Greening C, Johnston SG**. **2021**. Bark-dwelling methanotrophic bacteria decrease methane emissions from trees. *Nature Communications* **12**: 2127.

**Kattge J, Bönisch G, Díaz S, Lavorel S, Prentice IC, Leadley P, Tautenhahn S, Werner GDA, Aakala T, Abedi M, *et al.* 2020**. TRY plant trait database – enhanced coverage and open access. *Global Change Biology* **26**: 119–188.

**Kizlinski ML, Orwig DA, Cobb RC, Foster DR**. **2002**. Direct and indirect ecosystem consequences of an invasive pest on forests dominated by eastern hemlock. *Journal of Biogeography* **29**: 1489–1503.

**Knief C**. **2015**. Diversity and habitat preferences of cultivated and uncultivated aerobic methanotrophic bacteria evaluated based on *pmoA* as molecular marker. *Frontiers in Microbiology* **6**: 1346.

**Leung PM, Jeffrey LC, Bay SK, Gomez-Alvarez P, Hall M, Johnston SG, Dittmann J, Deschaseaux E, Hopkins B, Haskell J, *et al.* 2026**. Bark microbiota modulate climate-active gas fluxes in Australian forests. *Science* **391**: eadu2182.

**Lin LI**. **1989**. A concordance correlation coefficient to evaluate reproducibility. *Biometrics* **45**: 255–268.

**Louca S, Parfrey LW, Doebeli M**. **2016**. Decoupling function and taxonomy in the global ocean microbiome. *Science* **353**: 1272–1277.

**McEwan RW, Dyer JM, Pederson N**. **2011**. Multiple interacting ecosystem drivers: Toward an encompassing hypothesis of oak forest dynamics across eastern North America. *Ecography* **34**: 244–256.

**Moisan M-A, Maire V, Morency M-J, Martineau C**. **2026**. Tree tissues and species traits modulate the microbial methane-cycling communities of the tree phyllosphere. *Scientific Reports* **16**: 19771.

**Morotomi M, Nagai F, Watanabe Y**. **2012**. Description of *Christensenella minuta* gen. nov., sp. nov., isolated from human faeces, which forms a distinct branch in the order Clostridiales, and proposal of Christensenellaceae fam. nov. *International Journal of Systematic and Evolutionary Microbiology* **62**: 144–149.

**Noecker C, Sanchez J, Bisanz JE, Escalante V, Alexander M, Trepka K, Heinken A, Liu Y, Dodd D, Thiele I, *et al.* 2023**. Systems biology elucidates the distinctive metabolic niche filled by the human gut microbe *Eggerthella lenta*. *PLoS Biology* **21**: e3002125.

**Orwig DA, Foster DR**. **1998**. Forest response to the introduced hemlock woolly adelgid in southern New England, USA. *Journal of the Torrey Botanical Society* **125**: 60–73.

**Parkin TB, Venterea RT, Hargreaves SK. 2012**. Calculating the detection limits of chamber-based soil greenhouse gas flux measurements. *Journal of Environmental Quality* **41**: 705–715.

**Putkinen A, Siljanen HMP, Laihonen A, Paasisalo I, Porkka K, Tiirola M, Haikarainen I, Tenhovirta S, Pihlatie M. 2021**. New insight to the role of microbes in the methane exchange in trees: Evidence from metagenomic sequencing. *New Phytologist* **231**: 524–536.

**Ruaud A, Esquivel-Elizondo S, Cuesta-Zuluaga J de la, Waters JL, Angenent LT, Youngblut ND, Ley RE. 2020**. Syntrophy via interspecies $H_2$ transfer between *Christensenella* and *Methanobrevibacter* underlies their global cooccurrence in the human gut. *mBio* **11**: e03235–19.

**Semrau JD, DiSpirito AA, Vuilleumier S. 2011**. Facultative methanotrophy: False leads, true results, and suggestions for future research. *FEMS Microbiology Letters* **323**: 1–12.

**Siegenthaler A, Welch B, Pangala SR, Peacock M, Gauci V. 2016**. Technical note: Semi-rigid chambers for methane gas flux measurements on tree stems. *Biogeosciences* **13**: 1197–1207.

**Tukey JW. 1977**. *Exploratory data analysis*. Reading, MA: Addison-Wesley.

**Tveit AT, Hestnes AG, Robinson SL, Schintlmeister A, Dedysh SN, Jehmlich N, von Bergen M, Herbold C, Wagner M, Richter A, *et al.* 2019**. Widespread soil bacterium that oxidizes atmospheric methane. *Proceedings of the National Academy of Sciences, USA* **116**: 8515–8524.

**van Teeseling MCF, Pol A, Harhangi HR, van der Zwart S, Jetten MSM, Op den Camp HJM, van Niftrik L. 2014**. Expanding the verrucomicrobial methanotrophic world: description of three novel species of *Methylacidimicrobium* gen. nov. *Applied and Environmental Microbiology* **80**: 6782–6791.

**Waters JL, Ley RE. 2019**. The human gut bacteria Christensenellaceae are widespread, heritable, and associated with health. *BMC Biology* **17**: 83.

**Whiticar MJ. 1999**. Carbon and hydrogen isotope systematics of bacterial formation and oxidation of methane. *Chemical Geology* **161**: 291–314.

**Whittaker RH, Woodwell GM. 1967**. Surface area relations of woody plants and forest communities. *American Journal of Botany* **54**: 931–939.

**Wright MN, Ziegler A. 2017**. Ranger: A fast implementation of random forests for high dimensional data in C++ and R. *Journal of Statistical Software* **77**: 1–17.

**Xing D, Ren N, Li Q, Lin M, Wang A, Zhao L**. **2006**. *Ethanoligenens harbinense* gen. nov., sp. nov., isolated from molasses wastewater. *International Journal of Systematic and Evolutionary Microbiology* **56**: 755–760.

### Notes S1. Protocol: quantifying methane-cycling genes in living wood and soil by droplet digital PCR

This protocol quantifies:

- *mcrA* (methyl-coenzyme M reductase, methanogens);
- *pmoA* and *mmoX* (particulate and soluble methane monooxygenase, aerobic methanotrophs).

It applies to tree cores and soil and uses the Bio-Rad QX200 system. It follows Arnold *et al.* (2024), where the extraction, the probe-based *mcrA* assay, recovery and the detection limit were validated in wood. Abundances are expressed as gene copies per gram of dry wood or fresh soil. The protocol starts from freeze-dried, cryo-ground wood and sieved, frozen soil; sampling and wood preparation are described in Methods S1 and Arnold *et al.* (2024).

#### 1. Equipment and reagents

**Extraction**

- MagMAX Microbiome Ultra Nucleic Acid Isolation Kit (Thermo Fisher).
- KingFisher Apex with 96 deep-well head.
- Vortex mixer with a tube adapter.
- Optional: Zymo OneStep-96 PCR Inhibitor Removal Kit.

**ddPCR**

- QX200 Droplet Generator and Droplet Reader; DG8 cartridges and gaskets.
- PX1 PCR Plate Sealer and pierceable foil seals.
- Thermal cycler with a deep-well block.
- QuantaSoft software.
- ddPCR Supermix for Probes (No dUTP) and Droplet Generation Oil for Probes.
- QX200 ddPCR EvaGreen Supermix and Droplet Generation Oil for EvaGreen.
- Nuclease-free water.
- Synthetic double-stranded DNA standards (gBlocks) for each target.

**Oligonucleotides (5′→3′)**

| Target | Oligonucleotide | Sequence | Final concentration | Source |
|---|---|---|---|---|
| *mcrA* | Forward | ACGACYTRCAGGAYCAGTGY | 900 nM | Arnold *et al.* (2024) |
| | Probe (FAM) | WGGWCCWAACTAYCCBAACTACG | 250 nM | |
| | Reverse | TGGTGWCCBACGTTCATYG | 900 nM | |
| *pmoA* | 189f | GGNGACTGGGACTTCTGG | 180 nM | Bourne *et al.* (2001) |
| | mb661r | CCGGMGCAACGTCYTTACC | 180 nM | |
| *mmoX* | 536f | CGCTGTGGAAGGGCATGAAGCG | 180 nM | Fuse *et al.* (1998) |
| | 898r | GCTCGACCTTGAACTTGGAGCC | 180 nM | |

The *mcrA* amplicon is 118 bp; the *pmoA* and *mmoX* amplicons are predicted at 508 and 363 bp from the primer binding sites in *Methylococcus capsulatus* Bath (Table S1). The *pmoA* primers do not target the divergent *pmoA* of verrucomicrobial and NC10 methanotrophs and under-detect USCα (Knief, 2015). *Methylocella*, which lacks pMMO, is detected by *mmoX* only.

#### 2. DNA extraction

1. Weigh 100 mg of dry wood powder or 250 mg of fresh soil into the kit bead tube, and record the mass to 0.1 mg.
2. Add 800 µL of lysis buffer. Bead-beat for 10 min at maximum speed on the vortex mixer.
3. Centrifuge to pellet debris.
4. Transfer 250 µL of cleared lysate to the processing plate.

5. Run the KingFisher program MagMAX_Microbiome_Soil_Liquid_Buccal_v1.
6. Elute into 75 µL (wood) or 100 µL (soil).
7. Include one extraction blank (lysis buffer only) per plate.
8. For tissues rich in polyphenols (bark, leaves, roots, litter, decayed wood), or where inhibition is suspected, pass the eluate through the Zymo OneStep-96 PCR Inhibitor Removal Kit.
9. Store eluates at −80 °C.

#### 3. Reaction set-up

Prepare a master mix for n + 10% reactions on ice. Add the template last.

**Probe assay (*mcrA*), 25 µL per reaction**

| Component | µL |
|---|---|
| ddPCR Supermix for Probes (No dUTP), 2× | 12.5 |
| Forward primer, 10 µM | 2.25 |
| Reverse primer, 10 µM | 2.25 |
| Probe, 10 µM | 0.625 |
| Nuclease-free water | 4.875 |
| DNA extract | 2.5 |

**EvaGreen assays (*pmoA*, *mmoX*), 25 µL per reaction**

| Component | µL |
|---|---|
| QX200 ddPCR EvaGreen Supermix, 2× | 12.5 |
| Forward and reverse primers | to the final concentration above |
| Nuclease-free water | to 22.5 |
| DNA extract | 2.5 |

On every plate include:

- a gBlock positive standard at about $10^3$–$10^4$ copies per reaction;
- at least two no-template controls (water in place of template);
- the extraction blanks.

Mix by pipetting, seal and spin down.

#### 4. Droplet generation

1. Load 20 µL of each reaction and 70 µL of the matching droplet generation oil into a DG8 cartridge. Use oil for Probes or oil for EvaGreen to match the supermix.
2. Generate the droplets.
3. Transfer 40 µL of droplets slowly to a 96-well plate, with the pipette tip against the well wall.
4. Heat-seal the plate (180 °C, 5 s) and start cycling within 30 min.

#### 5. Thermocycling

| Assay | Program | Ramp |
|---|---|---|
| *mcrA* (probe) | 95 °C 10 min; 40 × [94 °C 30 s, 48 °C 80 s]; 98 °C 10 min; 4 °C hold | 2 °C $s^{-1}$ |
| *pmoA* | 95 °C 5 min; 40 × [95 °C 30 s, 52 °C 35 s, 72 °C 50 s]; 4 °C 5 min; 90 °C 5 min; 4 °C hold | 1 °C $s^{-1}$ |
| *mmoX* | 95 °C 5 min; 40 × [95 °C 30 s, 52 °C 30 s, 72 °C 45 s]; 4 °C 5 min; 90 °C 5 min; 4 °C hold | 1.5 °C $s^{-1}$ |

#### 6. Reading and classification

1. Read the plate on the Droplet Reader.
2. In QuantaSoft, reject wells with fewer than 5,000 accepted droplets. A typical well has 12,000–20,000.
3. Inspect the 1-D amplitude plot for each plate. Set the threshold manually in the gap between the negative and positive droplet clouds, and use the same threshold for all wells of a plate.
    - With the probe assay the clouds are well separated (positives about 1,900 and negatives about 700 amplitude units), so the call is insensitive to the exact threshold.
    - EvaGreen clouds are closer together. Export a second, more conservative threshold as a sensitivity check.
4. Export for every well:
    - concentration (copies $\mu L^{-1}$);
    - positive, negative and accepted droplets;
    - the threshold used.

#### 7. Calculation

QuantaSoft reports the concentration $C$ in copies per µL of the 1× reaction, from the fraction of positive droplets by Poisson statistics (droplet volume 0.85 nL):

$C = -\ln(1 - \text{positives / accepted droplets}) / 0.00085$

The “copies per 20 µL well” field is $C \times 20$ and is a display convention only. Correct for the template’s dilution into the reaction, then scale to the eluate, to the whole lysate and to the sample mass:

copies $g^{-1}$ = $C$ × (25 µL reaction / 2.5 µL template) × elution volume (µL) × (800 µL lysate / 250 µL processed) / sample mass (g)

The elution volume is 75 µL for wood and 100 µL for soil. 250 µL is the volume of lysate aimed for; transfers did not exceed it (127–238 µL in the method study’s records), so the lysate scaling is a lower bound, and it adds up to about two-fold variation between samples.

*Example.* One positive droplet among 16,000 gives $C$ = 0.0735 copies $\mu L^{-1}$. For 100 mg of wood this is 0.0735 × 10 × 75 × 3.2 / 0.1 ≈ 1,760 copies $g^{-1}$, the smallest non-zero value such a sample can report.

Report values without correcting for extraction efficiency (§8). Analyze abundances on a logarithmic scale.

#### 8. Quality control, detection and recovery

**No-template controls and blanks** should contain no positive droplets. Because each droplet is scored positive or negative at endpoint, a control with no positive droplets is a true zero. With these oligonucleotides, all probe-assay no-template controls were zero, and *pmoA* and *mmoX* controls were at background (Arnold *et al.*, 2024).

**Do not quantify *mcrA* in wood with intercalating-dye chemistry.** Published *mcrA* primers with EvaGreen gave low but persistent false positives in wood, varying with species and tissue (Fig. S3) (Arnold *et al.*, 2024).

**Limit of detection**

- The probe assay detects 3 copies per reaction consistently (6 of 6 replicates) and is linear over five orders of magnitude (slope 0.99, $R^2$ = 0.997) (Arnold *et al.*, 2024).
- This is the limit of detection: below it, a negative cannot be distinguished from Poisson chance. Only about two-thirds of a 20 µL well passes the reader, so 3 copies per reaction is about 2 positive droplets.

- Three copies per reaction is $C$ = 0.15 copies µL$^{-1}$. That is about 3,600 copies g$^{-1}$ for 100 mg of wood (about 360 copies in the sample, because each reaction receives about 1% of the sample's DNA) and about 1,900 copies g$^{-1}$ for 250 mg of soil (about 480 copies in the sample; 100 µL eluate).
- Including extraction losses, the method detects about 500 cells per 100 mg of dry wood.
- Samples between the single-droplet floor and this limit are real detections, but they may not be detected on repeat.

**Precision near the limit**

- Counting error in digital PCR is about 1/√(positive droplets): about 70% at the limit of detection and 35% at 8 positive droplets, a common limit of quantification (Forootan *et al.*, 2017).
- Analyze abundances on a log scale, where this error is a standard deviation of about 0.3 $\log_{10}$ units at the limit of detection (0.43 × CV).

| Threshold | Positive droplets | Counting CV | SD ($\log_{10}$) | Copies g$^{-1}$ wood (100 mg) | Copies g$^{-1}$ soil (250 mg) |
|---|---|---|---|---|---|
| Single droplet | 1 | 100% | 0.43 | 1,800 | 940 |
| Limit of detection (3 copies per reaction) | ≈2 | ≈70% | 0.30 | 3,600 | 1,900 |
| | 4 | 50% | 0.22 | 7,100 | 3,800 |
| Limit of quantification (CV ≤ 35%) | 8 | 35% | 0.15 | 14,000 | 7,500 |
| | 16 | 25% | 0.11 | 28,000 | 15,000 |

*Detection and counting precision of the probe* mcrA *assay by threshold,* for 16,000 accepted droplets.

**Recovery**

- Extraction recovered 22 ± 4% of gBlocks spiked onto ground field wood immediately before extraction, and 30 ± 4% of gBlocks added to lysis buffer alone (Arnold *et al.*, 2024).
- Both figures include processing only part of the lysate (§2, step 4): the lysis-buffer control recovers about the fraction processed, so the bead cleanup itself loses little.

- Freeze-drying and cryo-grinding lose more. In spiked sterile dowels, recovery relative to the lysis-buffer control fell from 43% when spiked just before extraction to 19% when spiked before freeze-drying (Arnold *et al.*, 2024).
- PCR inhibition in wood extracts was negligible: dilution series had slopes near 1, and an inhibitor-removal kit made no difference (Arnold *et al.*, 2024).
- Because values are scaled to the whole lysate (§7), the extraction loss that remains in reported values is about 18% (release from the wood; the cleanup loses little), plus freeze-drying and grinding.

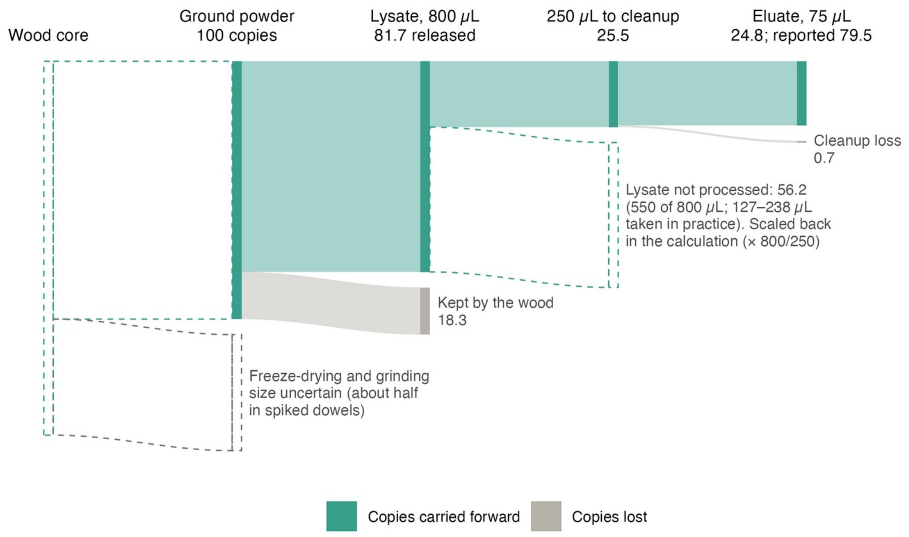


*Where target copies are lost between the core and the reaction.* Freeze-drying and cryo-grinding lose copies before extraction (dashed: about half in spiked sterile dowels, a different medium from field cores, so the size of this loss in cores is uncertain). Of 100 copies in the ground powder, the wood releases 81.7 (the mean field-wood recovery relative to the lysis-buffer control reported by Arnold *et al.* (2024)), 250 of the 800 µL of lysate is processed, and the cleanup keeps 97% (the lysis-buffer control's 30.4% divided by the 31% processed), so about 25 reach the eluate. The calculation scales the eluate back to the whole lysate (× 800/250), so about 79 are reported; the lysate not processed (dashed) is not a loss in reported values. The volume actually drawn off above the pellet varies (127–238 µL in the method study's records), so reported values can differ about two-fold between otherwise identical samples. Transferring 2.5 µL of eluate into each reaction is a subsample that the calculation scales back, not a loss. Values from Arnold *et al.* (2024).

**Record for every plate**

- sample masses;

- elution, template and reaction volumes;
- reagent lots;
- the thresholds, with an image of the amplitude plot;
- accepted droplets per well;
- standards, no-template controls and blanks, labeled as such in the export.


**Arnold W, Gewirtzman J, Raymond PA, Bradford MA, Butler C, Peccia J**. **2024**. A method for sampling the living wood microbiome. *Methods in Ecology and Evolution* **15**: 1084–1096.

**Bourne DG, McDonald IR, Murrell JC**. **2001**. Comparison of *pmoA* PCR primer sets as tools for investigating methanotroph diversity in three Danish soils. *Applied and Environmental Microbiology* **67**: 3802–3809.

**Forootan A, Sjöback R, Björkman J, Sjögreen B, Linz L, Kubista M**. **2017**. Methods to determine limit of detection and limit of quantification in quantitative real-time PCR (qPCR). *Biomolecular Detection and Quantification* **12**: 1–6.

**Fuse H, Ohta M, Takimura O, Murakami K, Inoue H, Yamaoka Y, Oclarit JM, Omori T**. **1998**. Oxidation of trichloroethylene and dimethyl sulfide by a marine *Methylomicrobium* strain containing soluble methane monooxygenase. *Bioscience, Biotechnology, and Biochemistry* **62**: 1925–1931.

**Knief C**. **2015**. Diversity and habitat preferences of cultivated and uncultivated aerobic methanotrophic bacteria evaluated based on *pmoA* as molecular marker. *Frontiers in Microbiology* **6**: 1346.

### Supporting Information: Figures S1–S28

#### Methane emissions from upland tree stems reflect a balance of microbial production and consumption

Figures are numbered in order of first citation in the main text. Methods S1–S7, Notes S1 and Tables S1–S8 are supplied separately.

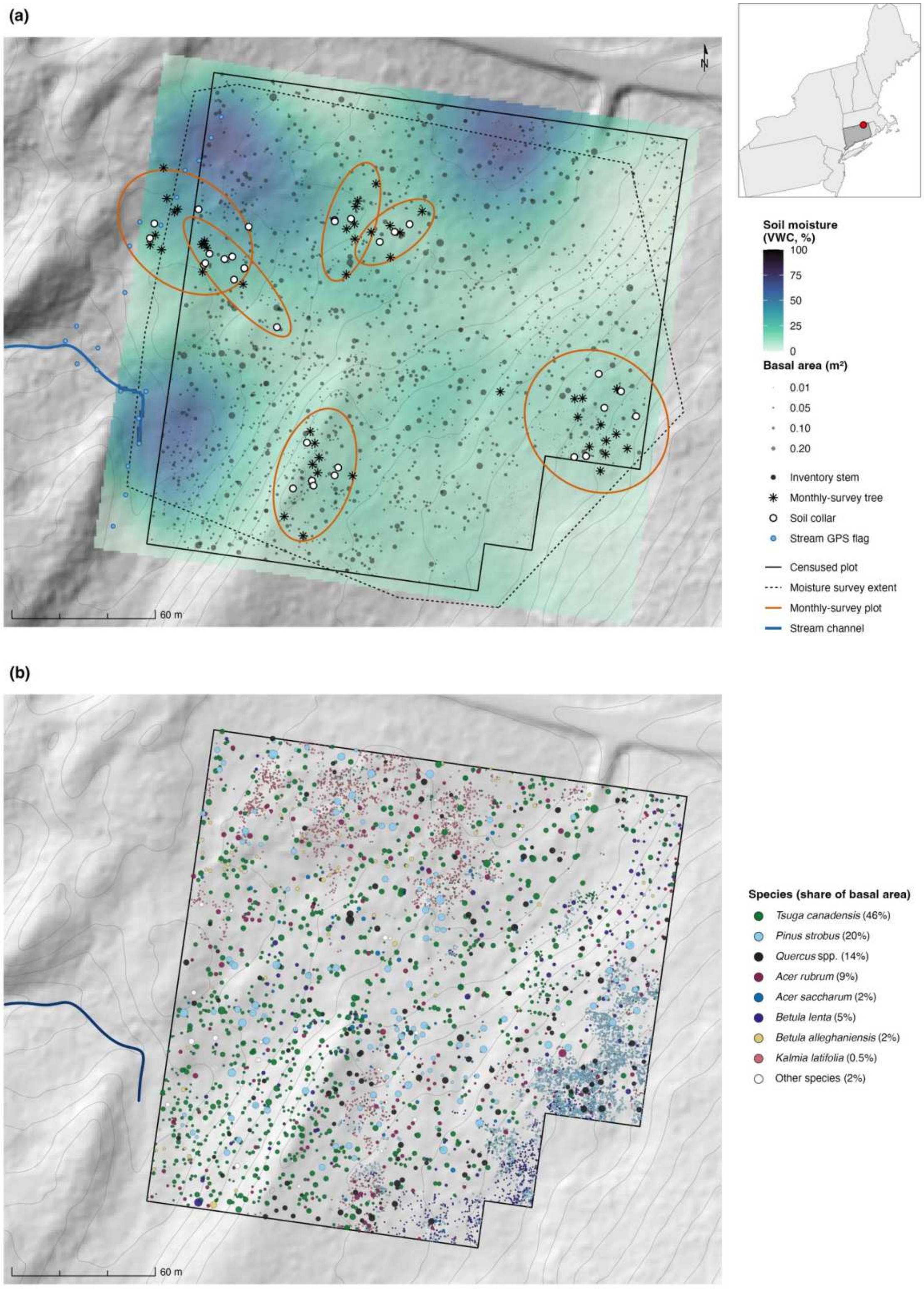


**Figure S1. Study site, soil moisture and tree species.** Both panels: hillshade and 2 m contours from 1 m lidar (2016 Connecticut statewide survey, USGS 3DEP); solid outline, the 4-ha inventory plot (3.72 ha censused); points, the 8,160 located inventory stems, sized by basal area. (a) Sampling design and soil moisture. Color, soil volumetric water content (VWC, %) from the December 2020 survey, interpolated with the thin-plate spline used in the analysis (Methods S6) and shown over the rectangle spanning all mapped trees and soil collars (plus 5 m); outside the dashed line (the extent of the survey), values are extrapolated. Orange ellipses, the monthly-survey plots; open circles, soil collars; asterisks, trees measured repeatedly in the 2020–2021 monthly survey; blue points, stream GPS flags; blue line, the defined stream channel, which lidar flow routing shows draining west from a low point midway along the flags (the flags to the north and south lie on near-flat wet ground). Inset, location in the northeastern United States (Connecticut shaded). (b) Inventory stems by species, with each group's share of basal area in the legend. *Quercus* spp. is *Q. rubra* 11.5%, *Q. velutina* 1.5% and *Q. alba* 0.9%; *Kalmia latifolia* is 24% of stems. Open circles, other species (2.3% of basal area together): unidentified or rare species 0.8%, *Prunus serotina* 0.5%, *Betula papyrifera* 0.4%, *Fagus grandifolia* 0.3%, *Fraxinus americana* 0.3% and *Acer pensylvanicum* <0.1%.

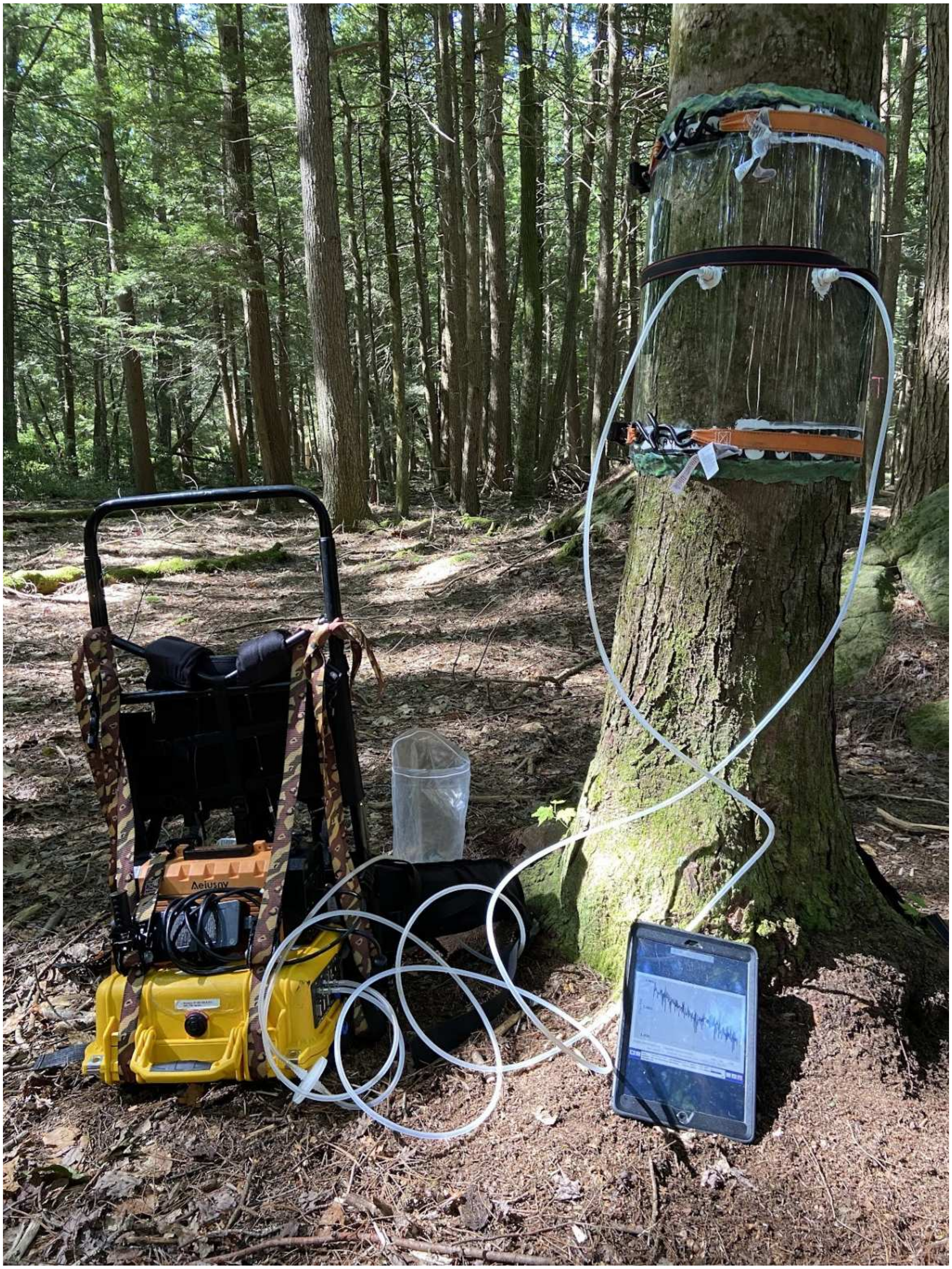
Aeiusny

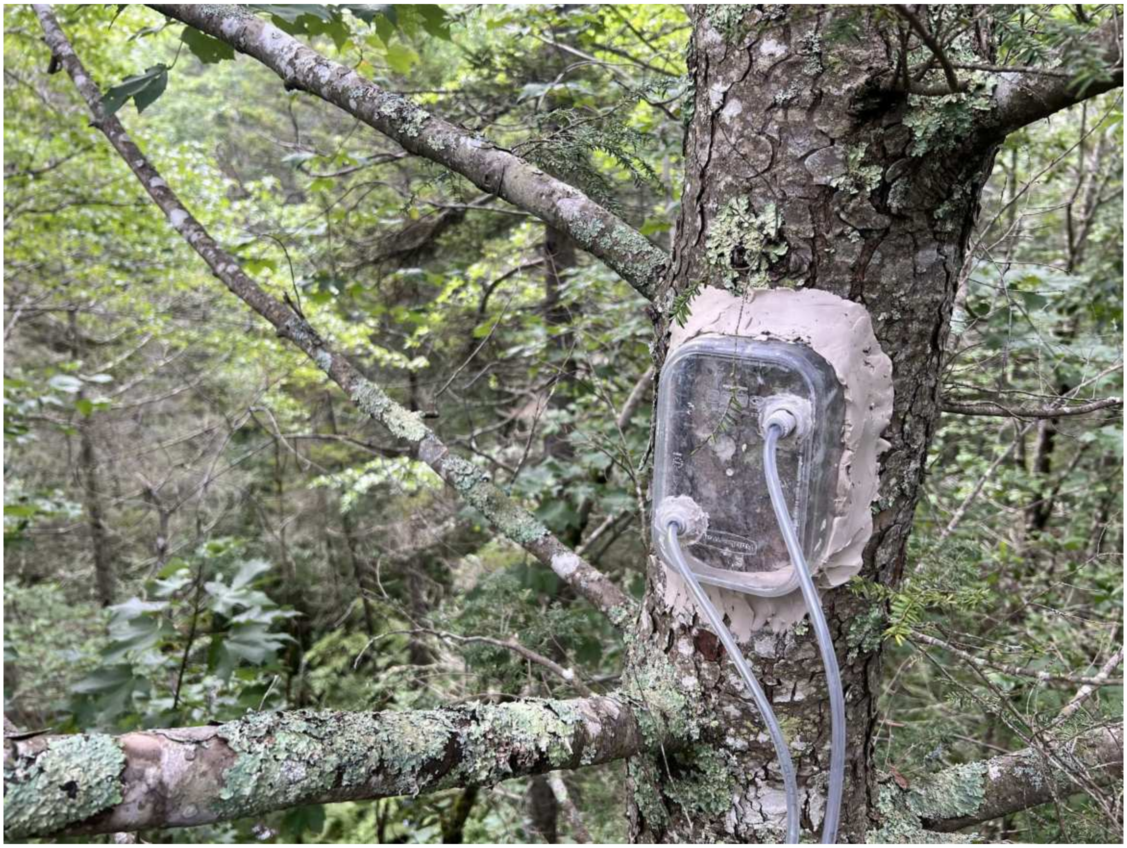

**Figure S2. Stem flux chambers.** (a) Semi-rigid chamber used in the 2020–2021 monthly survey. (b) Rigid chamber used in the 2021 height and molecular survey and the 2023 cross-species survey.

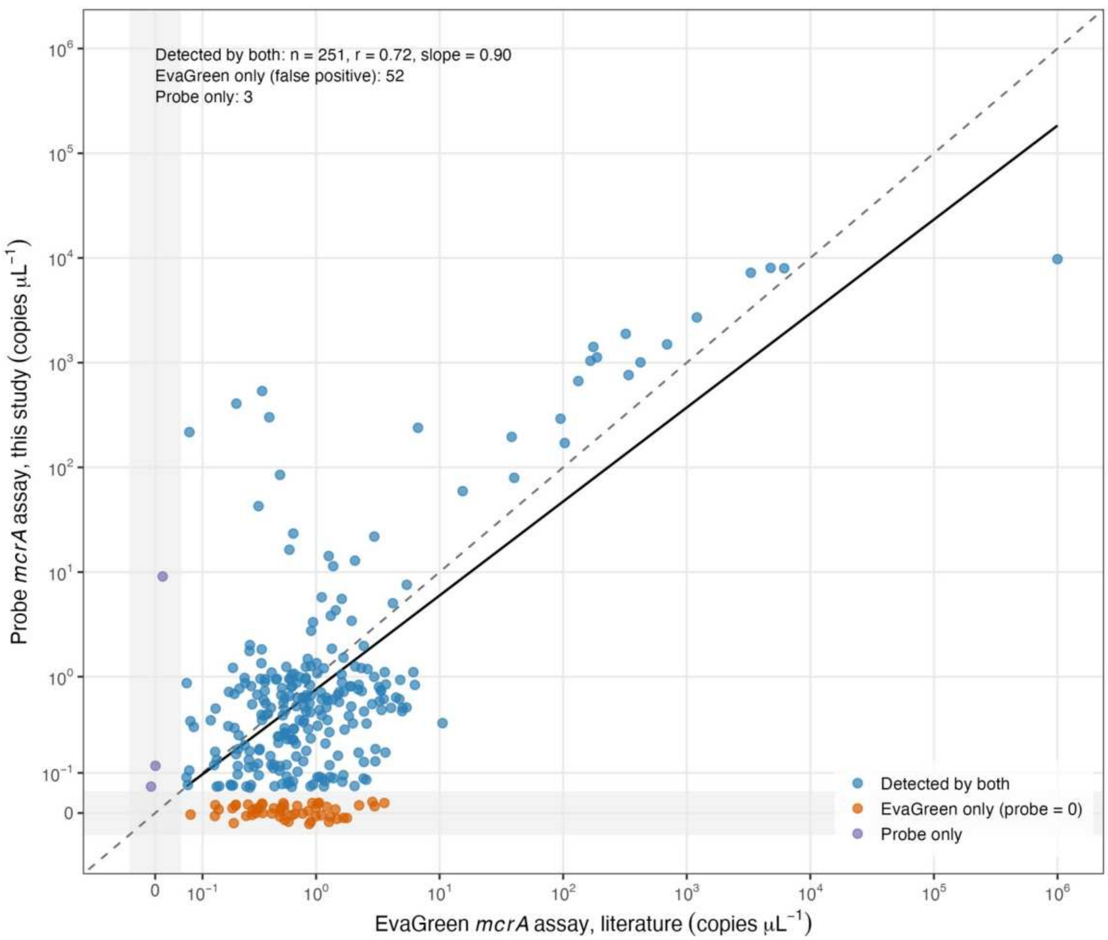


**Figure S3. Probe-based *mcrA* assay compared with the EvaGreen assay on the same extracts.** Concentrations (copies $\mu L^{-1}$) on both axes, pseudo-log scale. Blue, detected by both assays (Pearson r, slope and n annotated); orange, detected by EvaGreen only, which the probe assay treats as false positives; purple, detected by the probe only. Dashed line, 1:1; solid line, fit to samples detected by both.

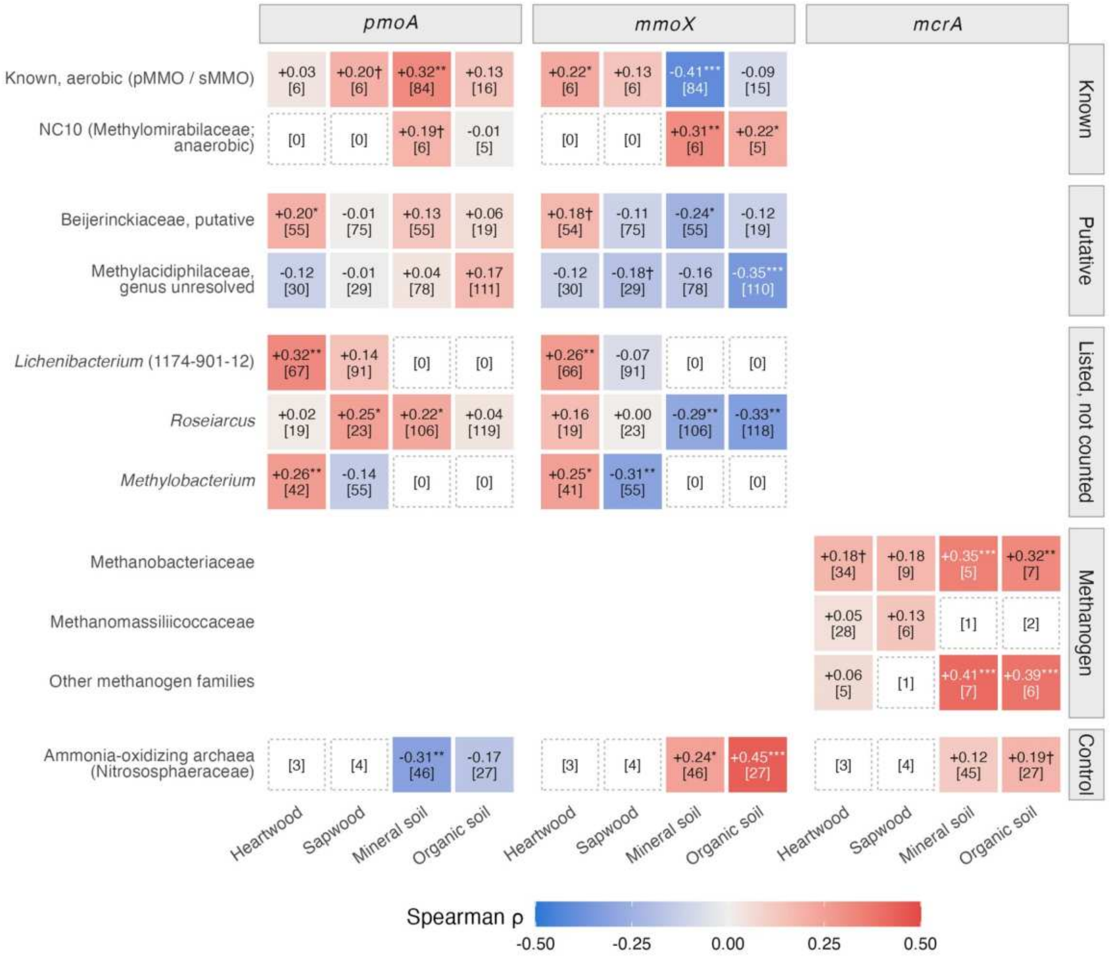


**Figure S4. 16S group abundance against the gene measured by ddPCR in the same sample.** Cells give Spearman ρ between the group's share of 16S reads and the gene's copies per bacterial 16S copy (gene by ddPCR, 16S by qPCR, in the same extract), by compartment and gene, with the number of samples in which the group is present in brackets; significance after Benjamini–Hochberg correction across all tests (* $q < 0.05$, ** $q < 0.01$, *** $q < 0.001$; † $p < 0.05$ uncorrected only). Dashed cells, group present in fewer than five samples (count in brackets), not tested. Rows follow the classification of Table S2. Table S4 gives the same comparison against gene copies per reaction (not normalized to 16S), so signs can differ. The *pmoA* primers do not target verrucomicrobial or NC10 *pmoA* and under-detect USCα.

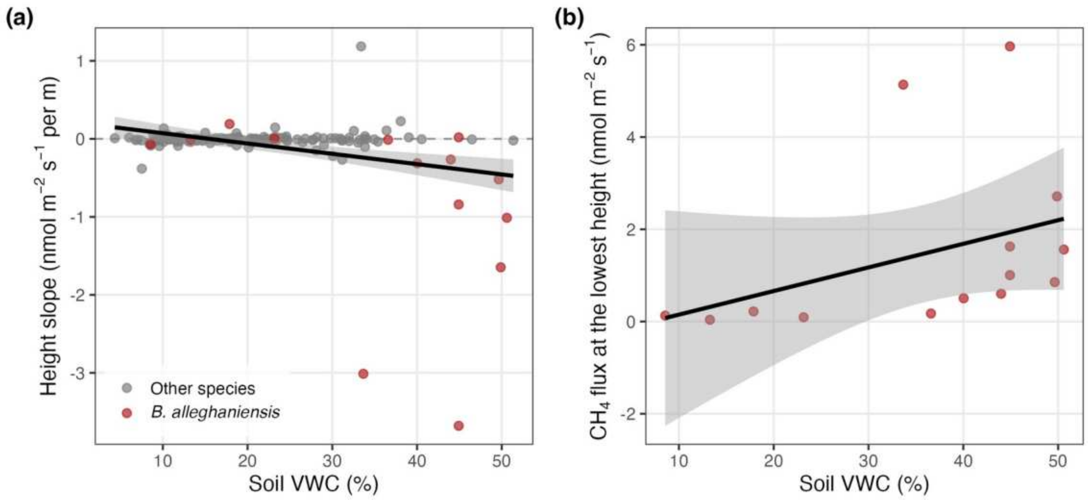


**Figure S5. Height slope of stem flux against soil moisture.** (a) Per-tree slope of flux with height against soil VWC for trees measured at three heights ($r = -0.32$, $p < 0.001$, $n = 138$ trees); *Betula alleghaniensis* in red. (b) Flux at the lowest measured height against soil VWC in *B. alleghaniensis* ($r = 0.39$, $p = 0.17$, $n = 14$ trees). Species medians of the per-tree slopes in (a): $r = -0.61$, $p = 0.047$ (11 species). Lines, linear regression with 95% CI.

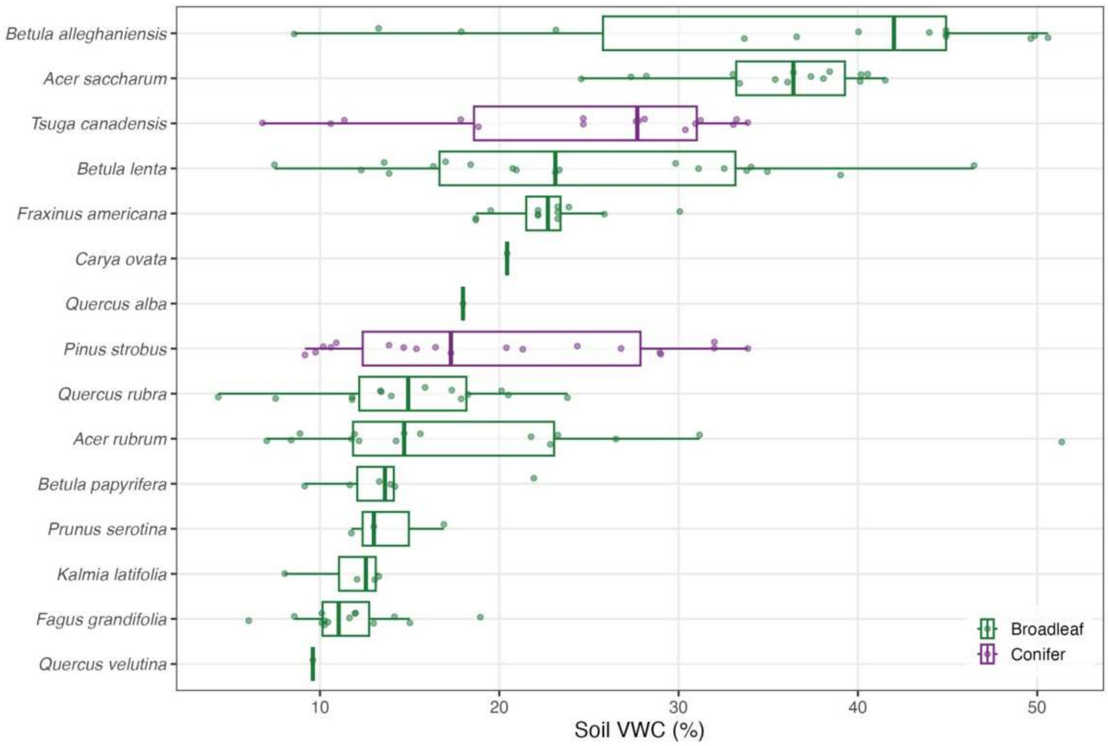


**Figure S6. Realized soil-moisture niche of each species.** Soil VWC (%) at each measured tree, by species; boxes show median and interquartile range. Green, broadleaf species; purple, conifers.

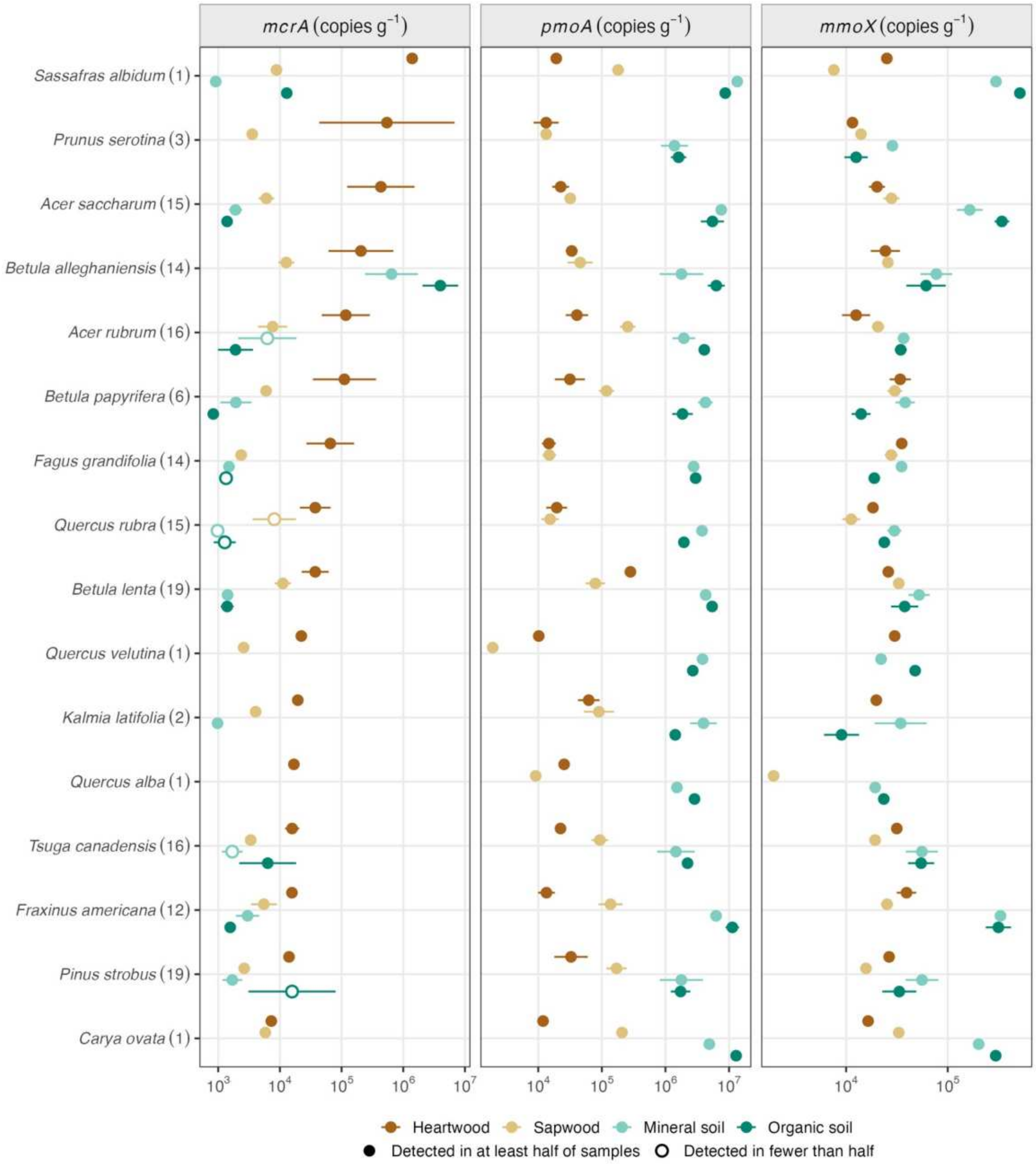


**Figure S7. Gene abundance by species for all 16 species, with *pmoA* and *mmoX* shown separately.** *mcrA*, *pmoA* and *mmoX* (copies $g^{-1}$; dry mass for wood, fresh mass for soil) by compartment, as in Fig. 4a: points, mean of $\log_{10}$ among positive samples ± SE; hollow points, fewer than half of samples positive; trees sampled in parentheses; species ranked by heartwood *mcrA*.

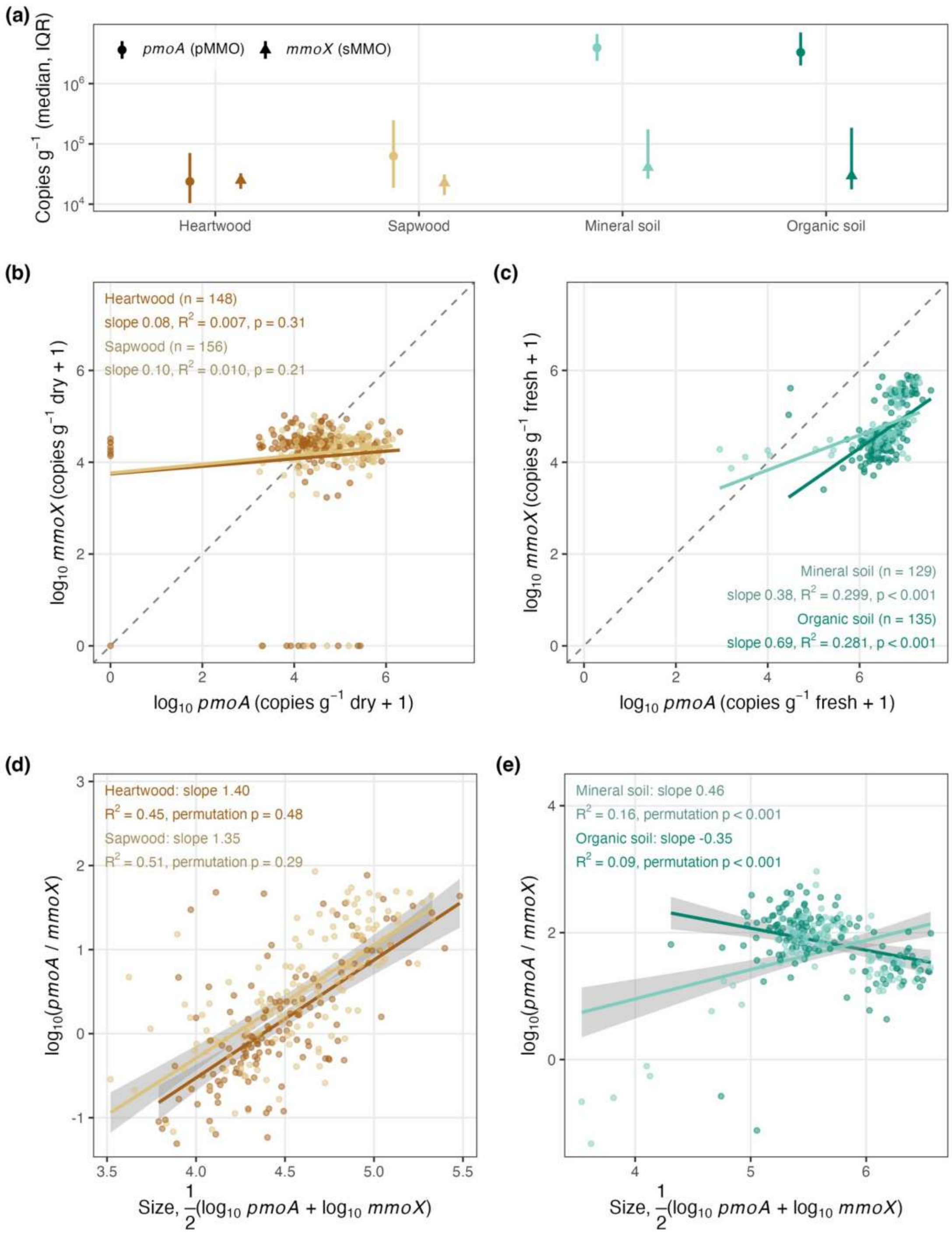


**Figure S8. *pmoA* and *mmoX* by compartment, and their coupling in wood and soil.** (a) Median and interquartile range of *pmoA* (circles) and *mmoX* (triangles) by compartment (copies $g^{-1}$; dry mass for wood, fresh mass for soil; nondetects as zero). (b, c) $\log_{10}$ *mmoX* against $\log_{10}$ *pmoA* in wood (b) and soil (c), with a fit per compartment; dashed line, 1:1. (d, e) $\log_{10}$(*pmoA*/*mmoX*) against size, the mean of the two log abundances, in wood (d) and soil (e), for samples with both genes detected. A steep ratio–size slope is expected mechanically when one gene varies little; the permutation p tests the observed slope against independent shuffling of the two genes. Wood matches that null; soil departs from it.

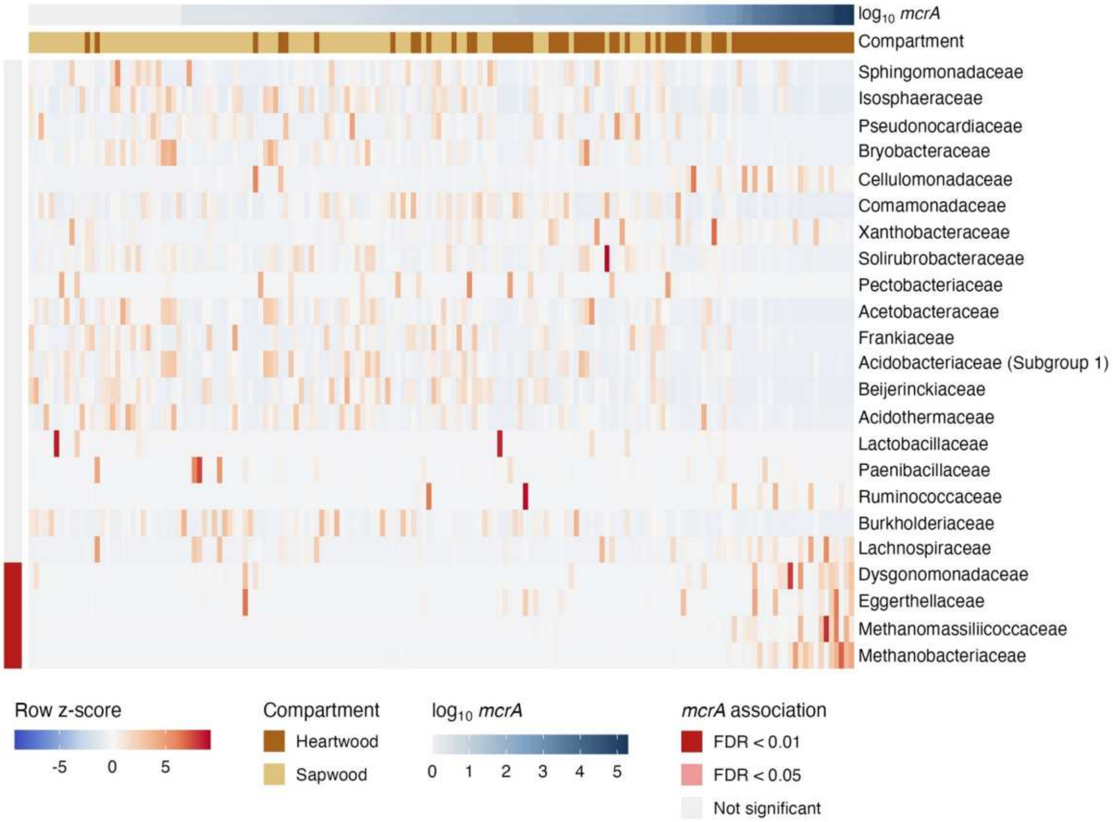


**Figure S9. Family-level 16S taxonomy associated with *mcrA* abundance.** Row-scaled relative abundance (z-score) of the 20 most abundant families and of families significantly associated with *mcrA* (FDR < 0.05). Samples (columns) are ordered by *mcrA* abundance and families (rows) by association t-statistic. Top bars, $\log_{10}$ *mcrA* and compartment; left bar, FDR significance from linear mixed-effects models with tree as a random intercept, controlling for compartment and total 16S concentration.

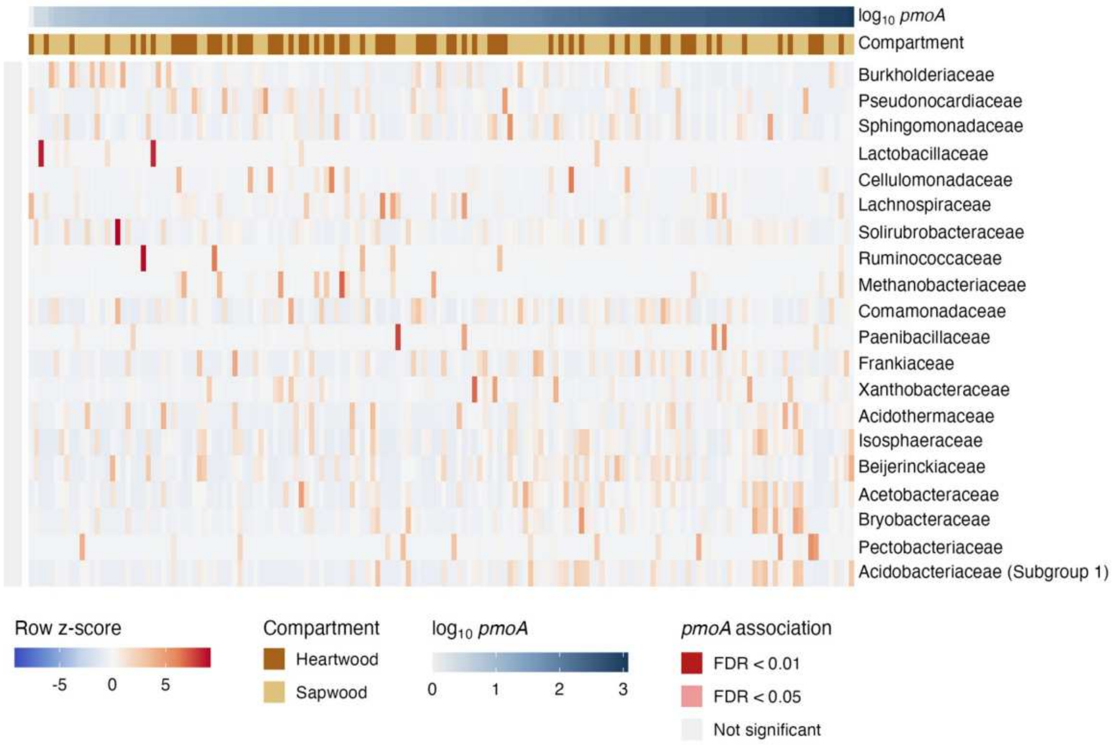


**Figure S10. Family-level 16S taxonomy associated with *pmoA* abundance.** As Fig. S9, for *pmoA*.

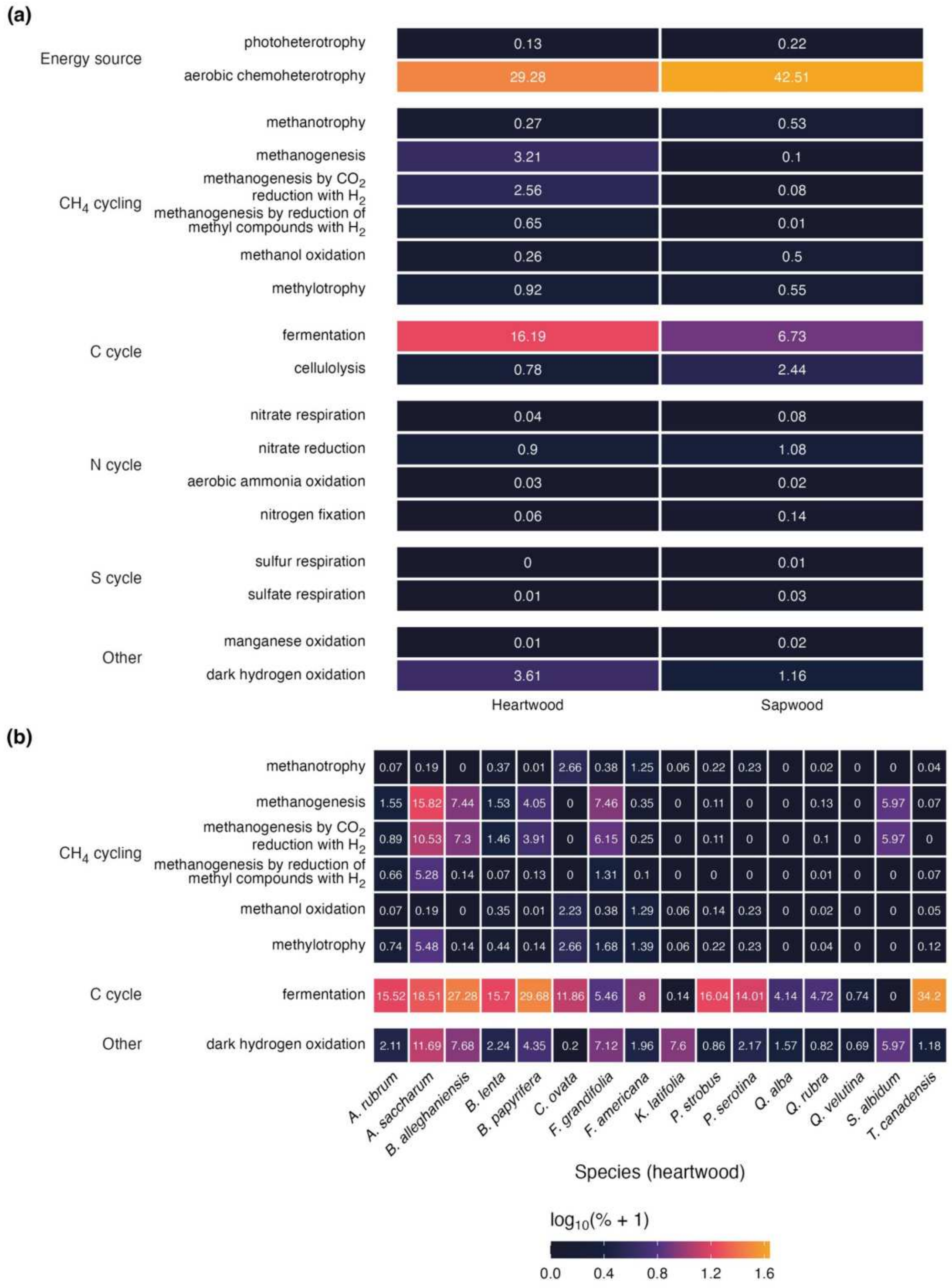


**Figure S11. FAPROTAX functional predictions in tree wood.** (a) Mean relative abundance ($\log_{10}(\% + 1)$) of predicted functions in heartwood and sapwood, grouped by category. (b) Selected $CH_4$-cycling, carbon-cycle and other functions across species (heartwood). Predicted percentages are fractions of the whole community and, because many ASVs are not annotated, lower bounds. FAPROTAX also lists methanogens under dark hydrogen oxidation and methylotrophy, so those categories here include them; Fig. 6b shows the functions without methanogens.

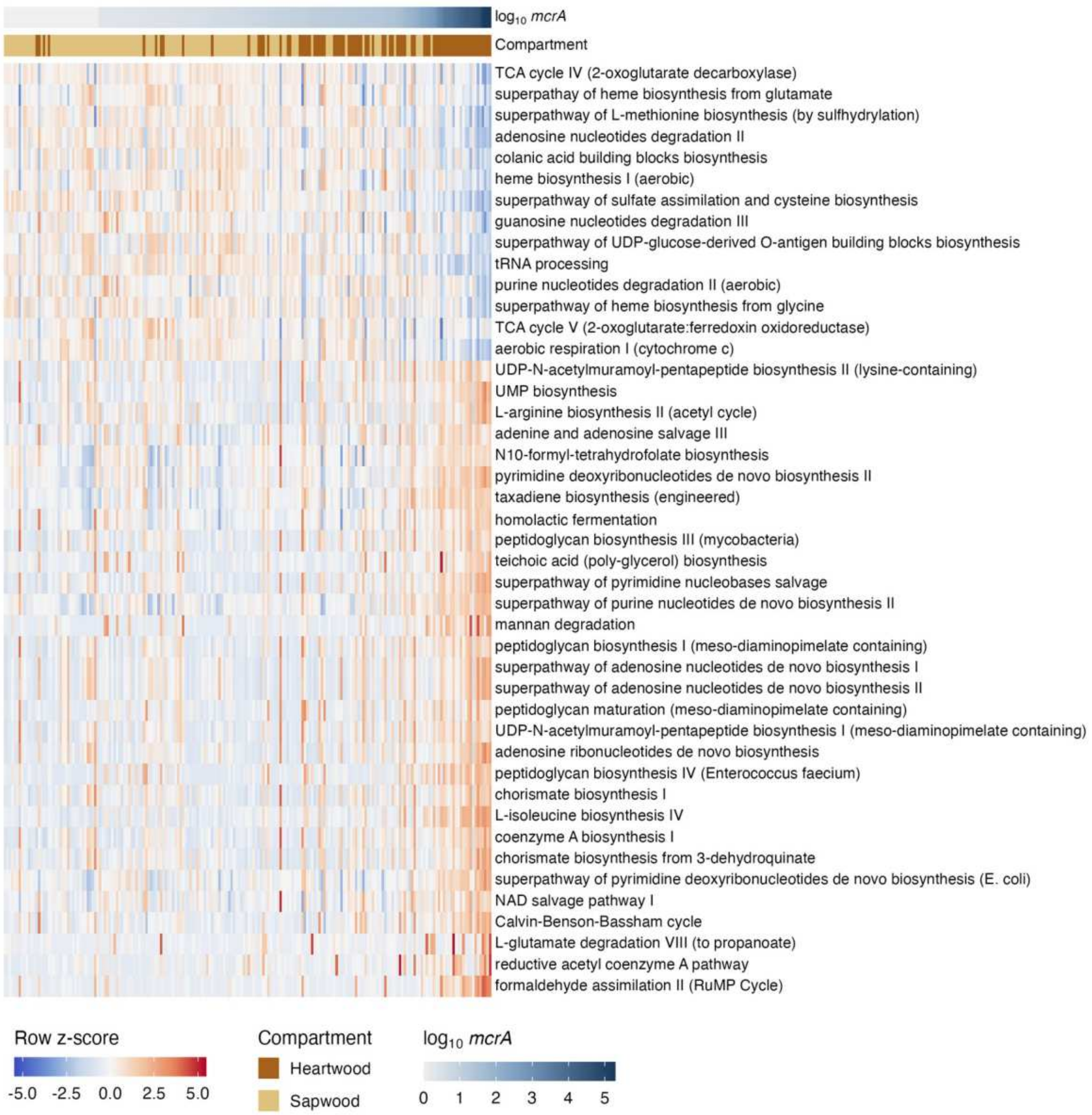


**Figure S12. MetaCyc pathways associated with *mcrA* abundance, excluding methanogen contributions.** Heatmap of the associations summarized in Fig. 6c, at a stricter threshold. PICRUSt2 pathway abundances significantly associated with *mcrA* (linear mixed-effects models with tree as a random intercept, controlling for compartment and 16S load; FDR $< 0.001$) after removing predicted contributions from methanogen-classified ASVs; pathways with more than 10% methanogen-ASV contribution are excluded; these include archaeal biosynthesis and methanogenesis pathways (tetrahydromethanopterin, archaeal mevalonate and flavin biosynthesis, methanogenesis from acetate), the expected methanogen signal. Samples are ordered by *mcrA* and pathways by association t-statistic; color, row-scaled abundance.

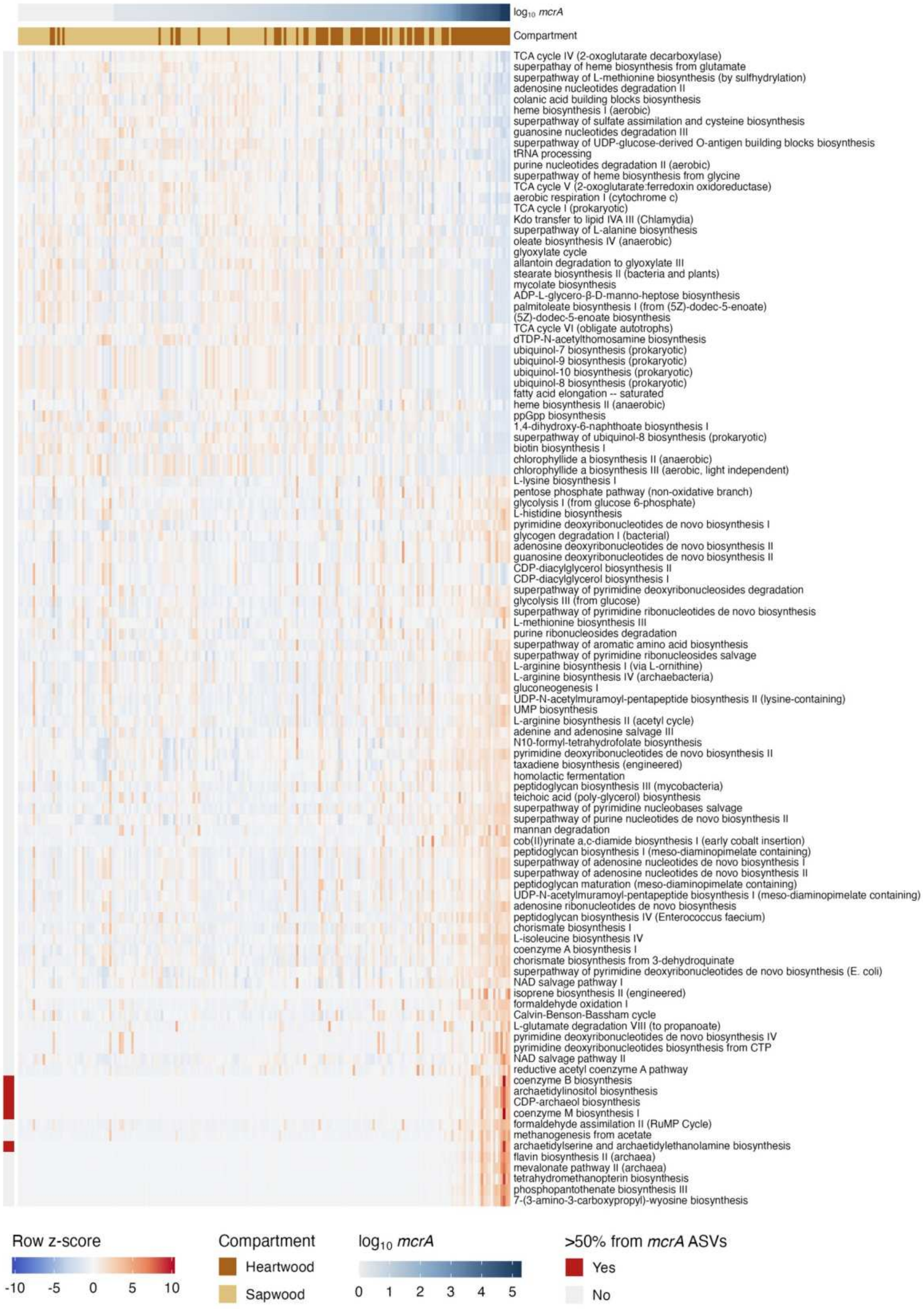

**Figure S13. MetaCyc pathways associated with *mcrA* abundance, all ASVs.** As Fig. S12 but retaining methanogen-classified ASVs and at FDR < 0.01; the row bar marks pathways with more than 50% of predicted abundance from methanogen-classified ASVs. The full association table is archived (Data Availability).

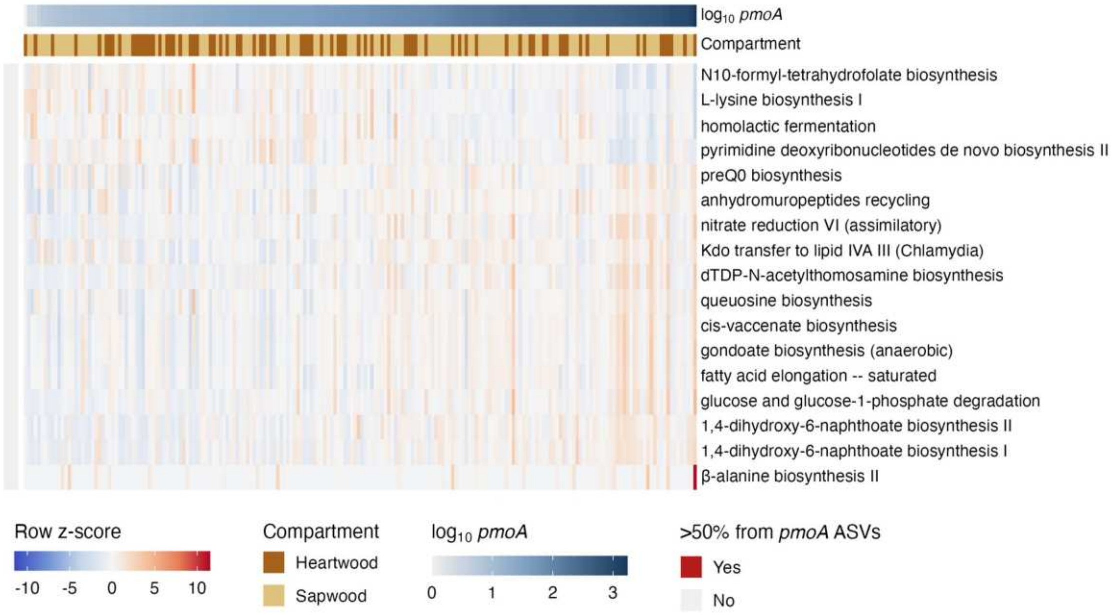


**Figure S14. MetaCyc pathways associated with *pmoA* abundance.** As Fig. S13, for *pmoA* (FDR < 0.01), retaining methanotroph-classified ASVs; the row bar marks pathways with >50% of predicted abundance from methanotroph-classified ASVs.

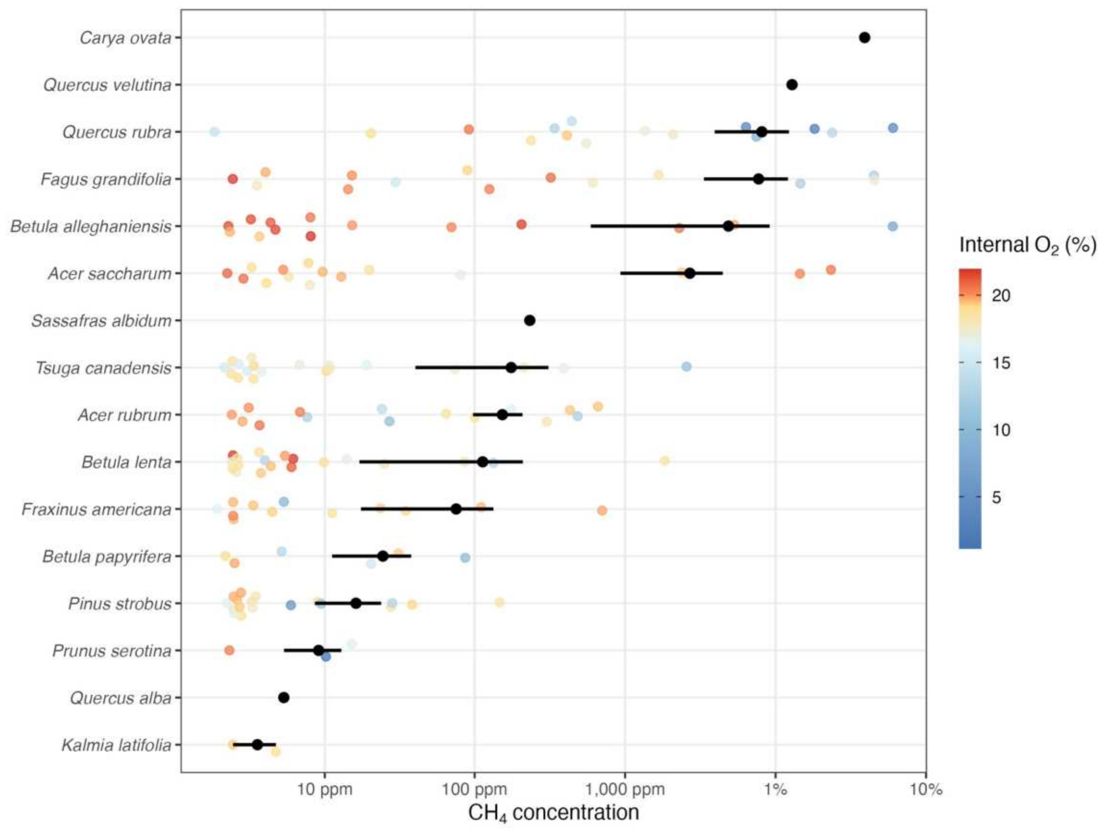


**Figure S15. Internal $CH_4$ concentrations by species.** Internal stem $CH_4$ for 16 species, ordered by mean concentration, on a pseudo-$log_{10}$ axis. Points colored by internal $O_2$ (blue, low; red, high); black points and bars, species mean ± SE.

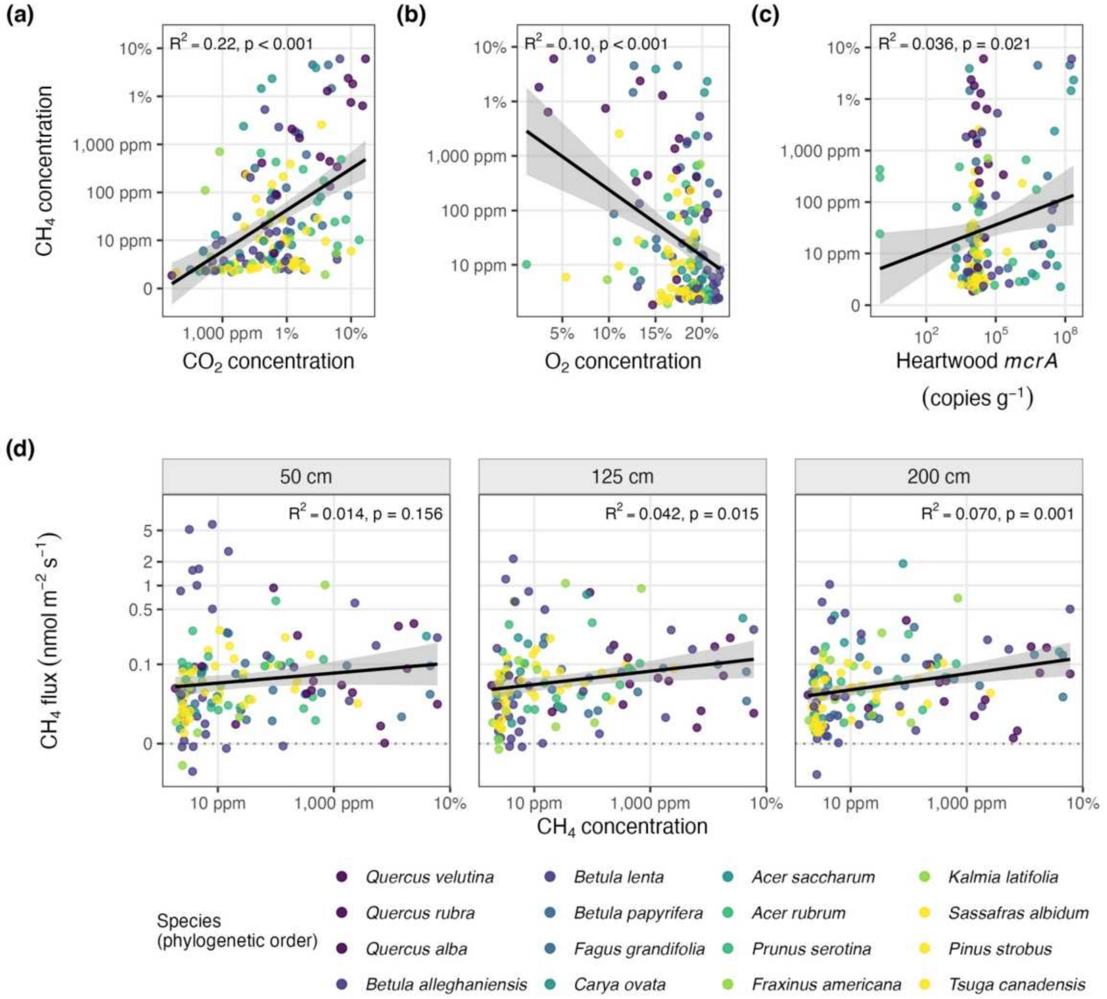


**Figure S16. Internal gases, *mcrA* and flux.** (a) Internal $CO_2$ against $CH_4$ (pseudo-$\log_{10}$ axes). (b) Internal $O_2$ (%) against $CH_4$. (c) Heartwood *mcrA* ($\log_{10}$ copies $g^{-1}$) against internal $CH_4$. (d) Internal $CH_4$ against stem flux at 50, 125 and 200 cm. Points colored by species, in phylogenetic order (colors as in Fig. S23); lines, linear regression with 95% CI; $R^2$ and p annotated.

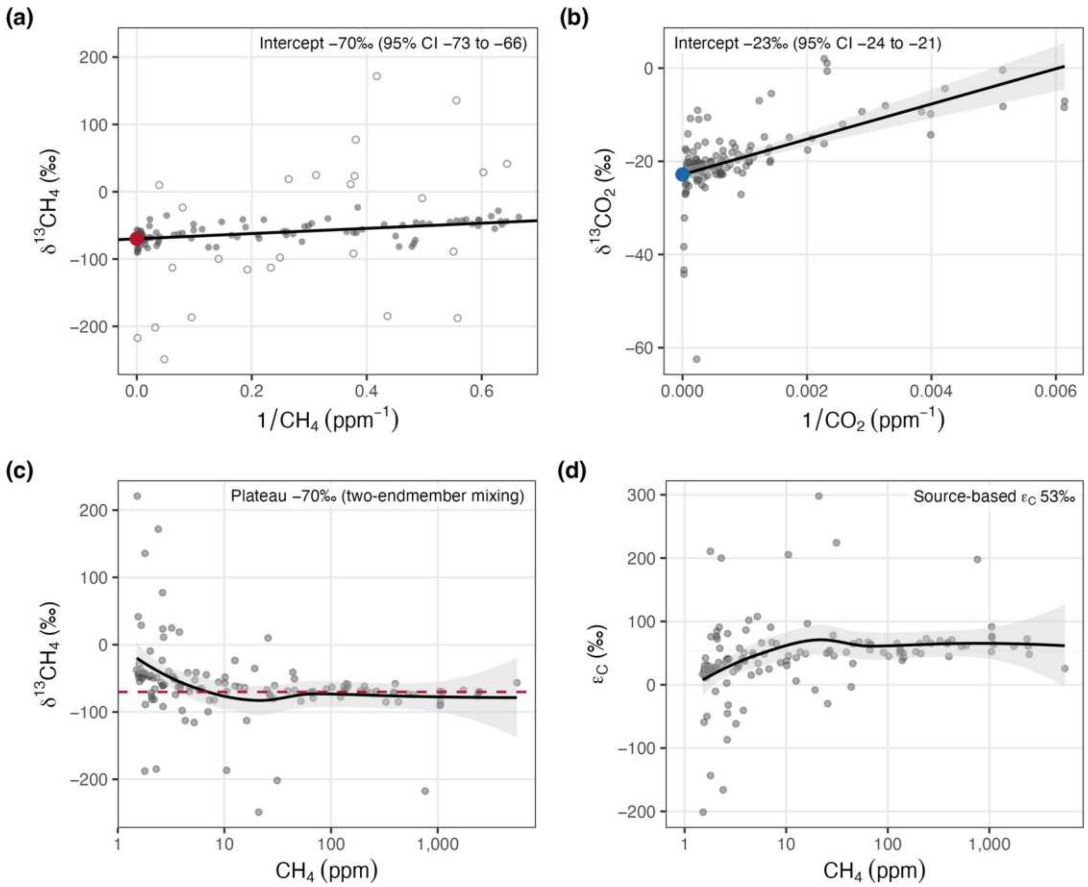


**Figure S17. Source signatures of internal $CH_4$ and $CO_2$.** (a) Keeling plot of $\delta^{13}CH_4$ against 1/[$CH_4$], fitted by MM robust regression (line); open circles, samples with weight below 0.5. The intercept (red point) estimates the tree-produced source and is given in the panel with its 95% CI. (b) The same for $CO_2$. (c) $\delta^{13}CH_4$ against internal $CH_4$ concentration; dilution by atmospheric $CH_4$ pulls low-concentration samples toward −47‰, and the dashed line marks the dilution-corrected source from two-endmember mixing. (d) Apparent $CO_2$–$CH_4$ fractionation ($\varepsilon_C$) against $CH_4$ concentration; line and band, smoother with 95% CI. Internal $CO_2$ is dominated by respiration, so $\varepsilon_C$ is apparent. n = 128 trees.

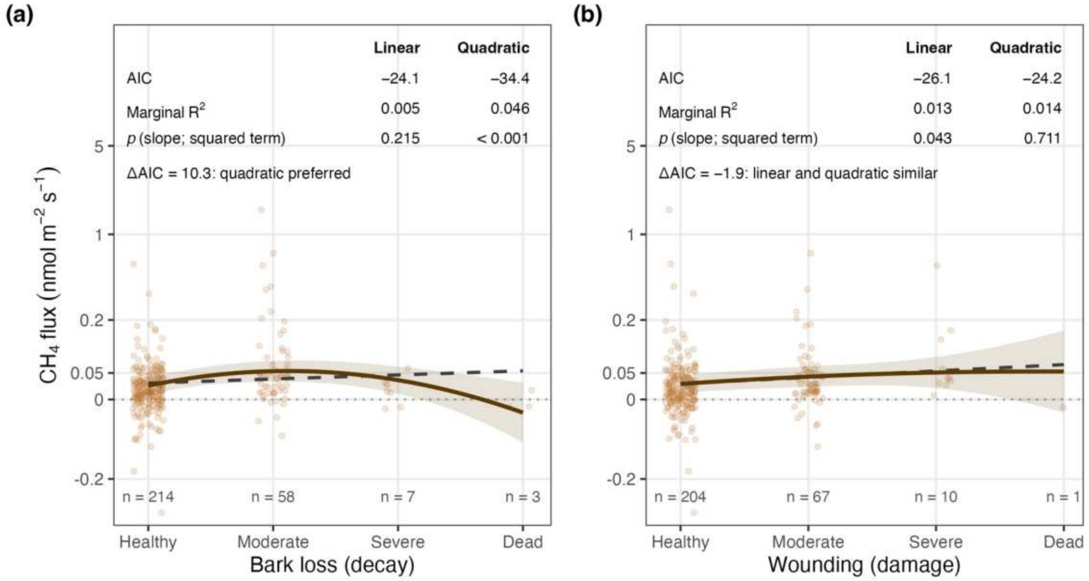


**Figure S18. Stem flux against visible deterioration (2023 survey).** (a) Bark loss; (b) wounding. Lines, linear (dashed) and quadratic (solid) mixed models with species as a random intercept (n = 282 measurements), with AIC, marginal $R^2$ and p in each panel; the number of measurements in each class is given at the foot. Bark loss favors the quadratic, a rise into moderate deterioration that holds for means, medians and ranks; wounding does not.

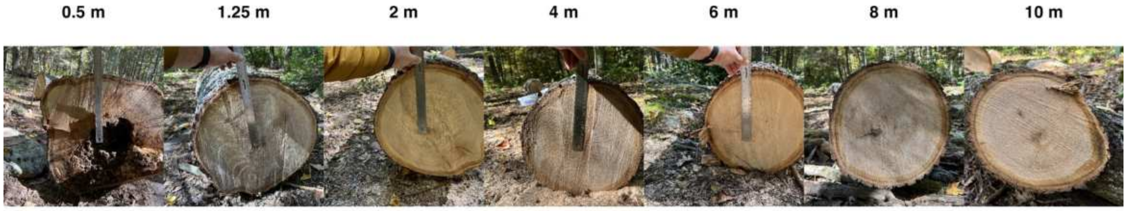


**Figure S19. Cross-sections of the felled black oak at each sampled height.** Photographs at 0.5, 1.25, 2, 4, 6, 8 and 10 m, showing a basal cavity, staining through the lower stem and clean wood above ~6 m.

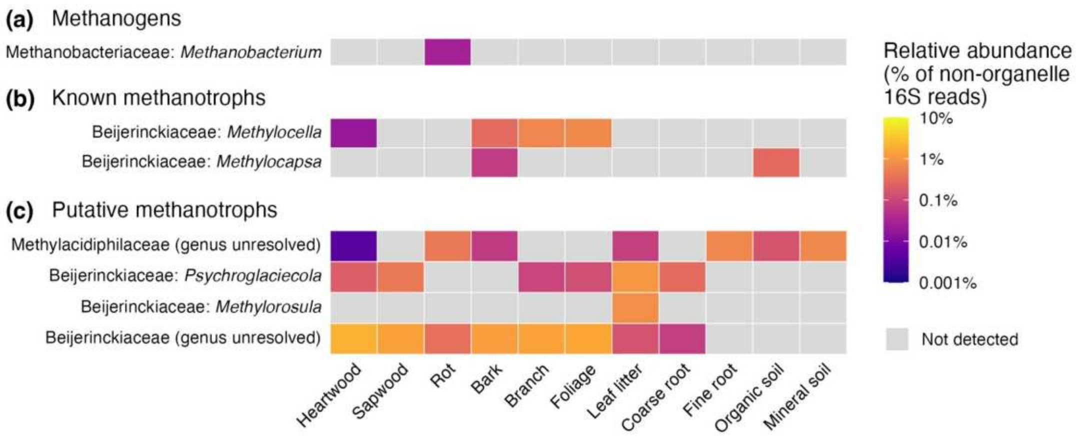


**Figure S20. Methane-cycling taxa across tissues of the felled black oak.** For each tissue, from heartwood to mineral soil, reads of each (a) methanogen, (b) Known methanotroph and (c) Putative methanotroph genus (Table S2), pooled over that tissue's libraries, as a percentage of their non-organelle 16S reads; log color scale; gray, not detected. Rows are labeled family: genus, or genus unresolved where SILVA assigns none.

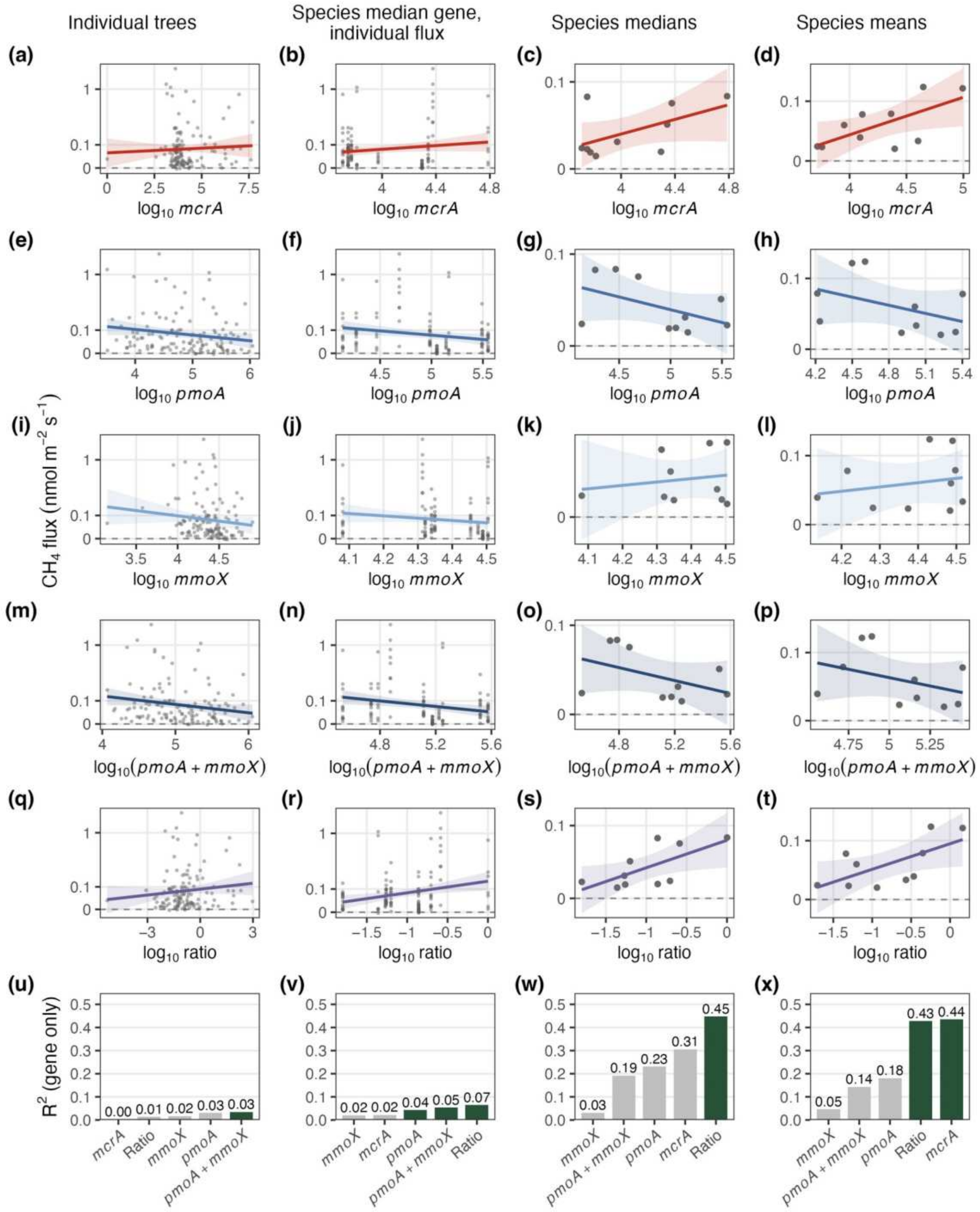


**Figure S21. Gene–flux relationships across scales of aggregation.** Rows: *mcrA*, *pmoA*, *mmoX*, *pmoA* + *mmoX*, and the methanogen:methanotroph ratio (area-weighted, $\log_{10}$). Columns: individual trees; species-median gene value against individual fluxes; species medians; species means. Flux is arcsinh-transformed before aggregation (nmol $m^{-2}$ $s^{-1}$ on that scale). Lines, linear regression with 95% CI. Bottom row (u–x), $R^2$ of each gene metric alone at each scale; dark green, $p < 0.05$. Gene-only $R^2$ is at most 0.04 among individual trees and rises to 0.45 for the ratio across species medians and to 0.44 for *mcrA* and 0.43 for the ratio across species means.

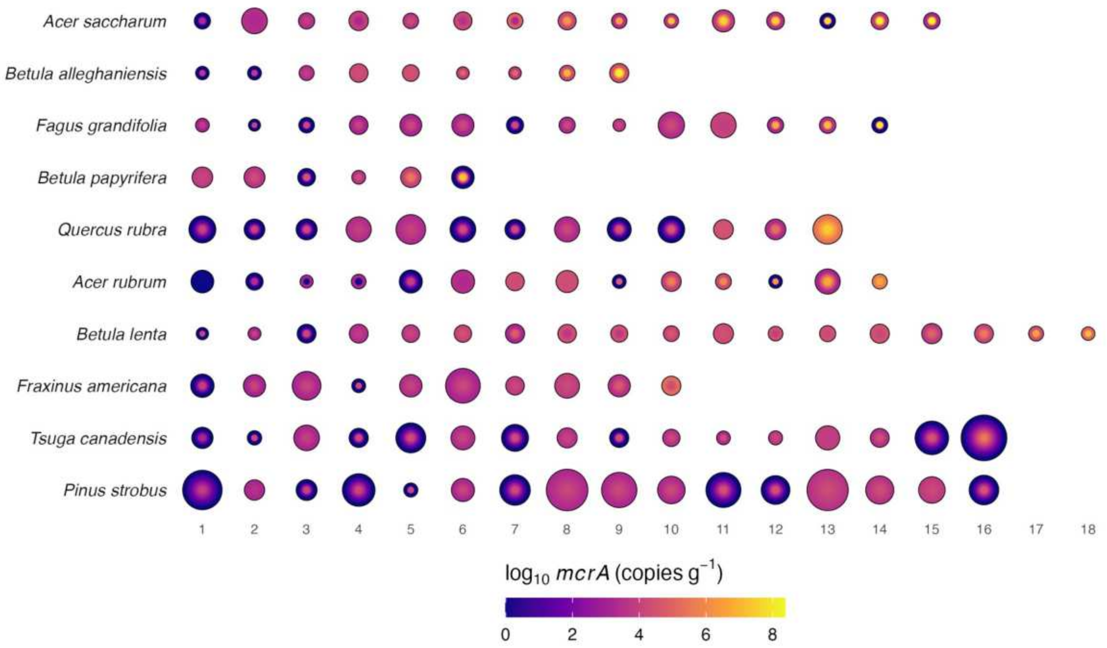


**Figure S22. Radial *mcrA* distribution in individual trees of ten species.** Each circle is one tree: $\log_{10}$ *mcrA* (copies $g^{-1}$) in heartwood (center) and sapwood (outer ring), diameter proportional to DBH. Rows are species.

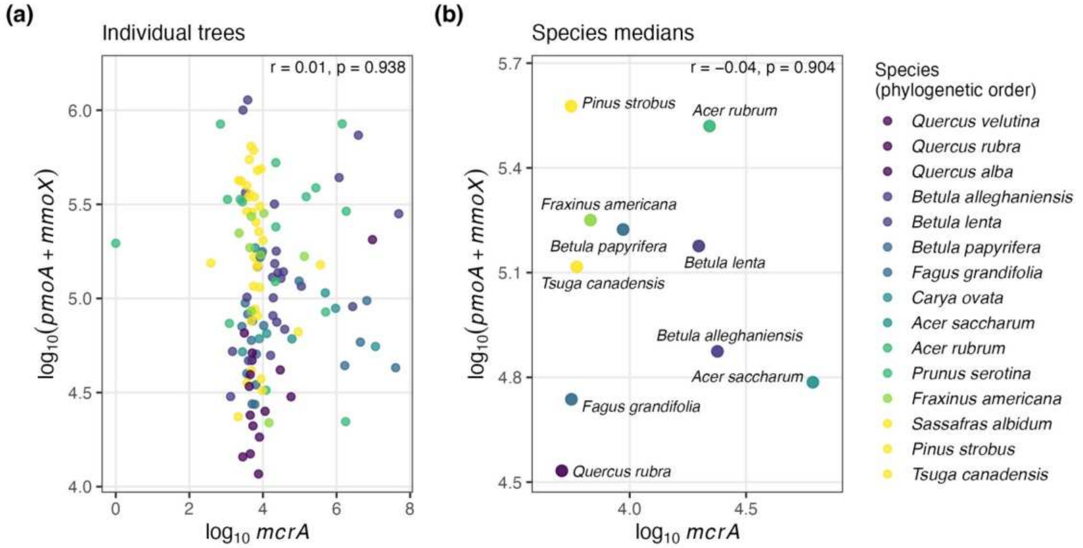


**Figure S23. Independence of methanogen and methanotroph abundance.** (a) Individual trees: $\log_{10}$ *mcrA* against $\log_{10}$(*pmoA* + *mmoX*).

(b) Species medians. Points colored by species, in phylogenetic order (colors as in Fig. S16). Pearson r and p annotated.

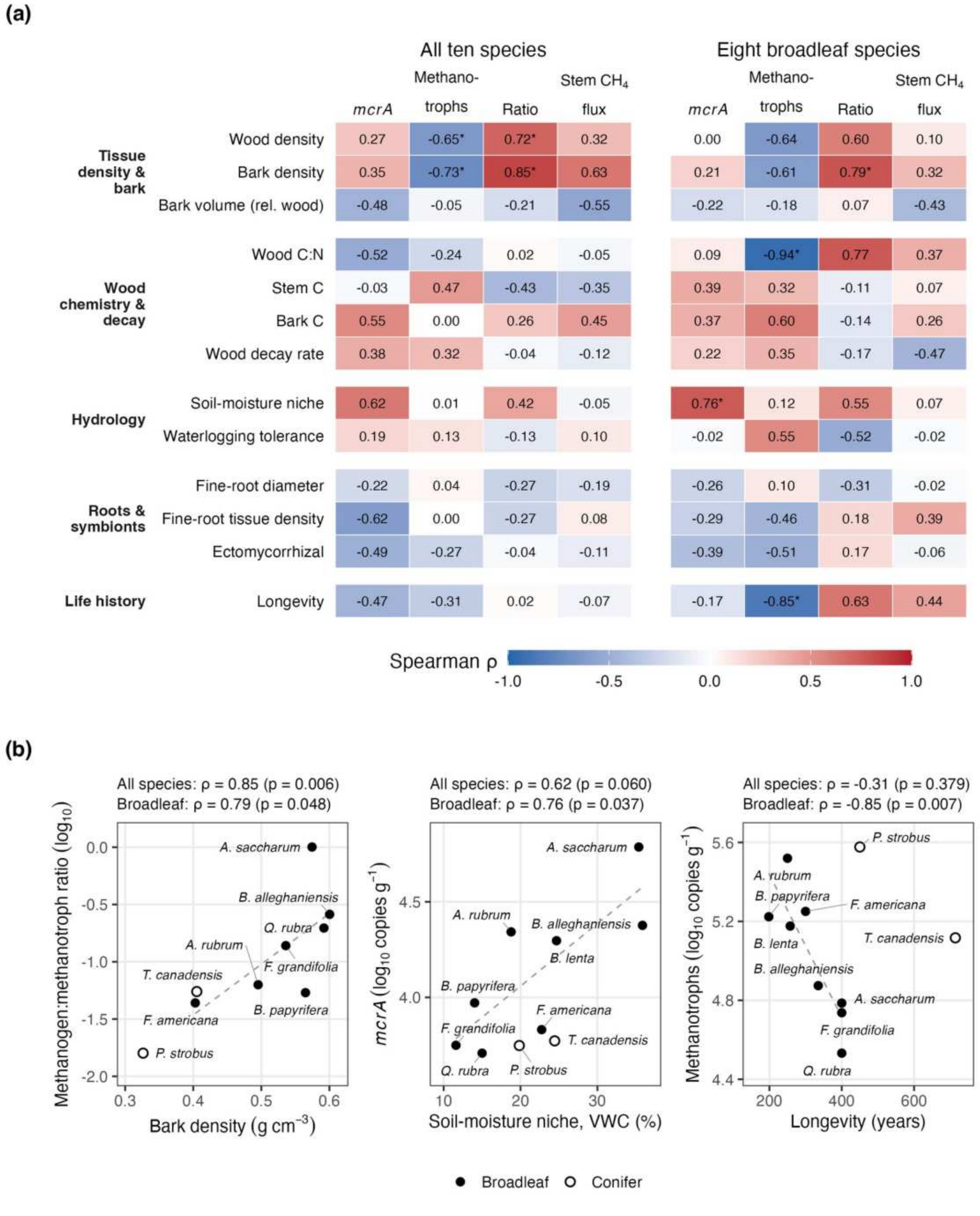


**Figure S24. Plant traits and the stem methane-cycling community.** (a) Spearman correlations between thirteen per-species traits (rows, grouped by the route by which they could act; inclusion rules in Methods S5) and four species-level quantities (*mcrA*, *pmoA* + *mmoX*, methanogen:methanotroph ratio, net stem flux) across all ten species with at least five sampled trees and five flux measurements (left) and across the eight broadleaf species alone (right); n = 8–10 and 6–8 by trait coverage. *, nominal $p < 0.05$, not corrected for multiple tests (no association passed false-discovery-rate correction). (b) The three associations named in the Results, those meeting the reporting rule of Methods S5: bark density and the ratio, soil-moisture niche and *mcrA*, and longevity and methanotroph abundance. Filled, broadleaf species; open, conifers; dashed lines, linear fits to the broadleaf species, spanning their range in each panel. Realized soil moisture is each species' mean VWC across its measured stems; other traits are from GWDD and TRY (Table S8).

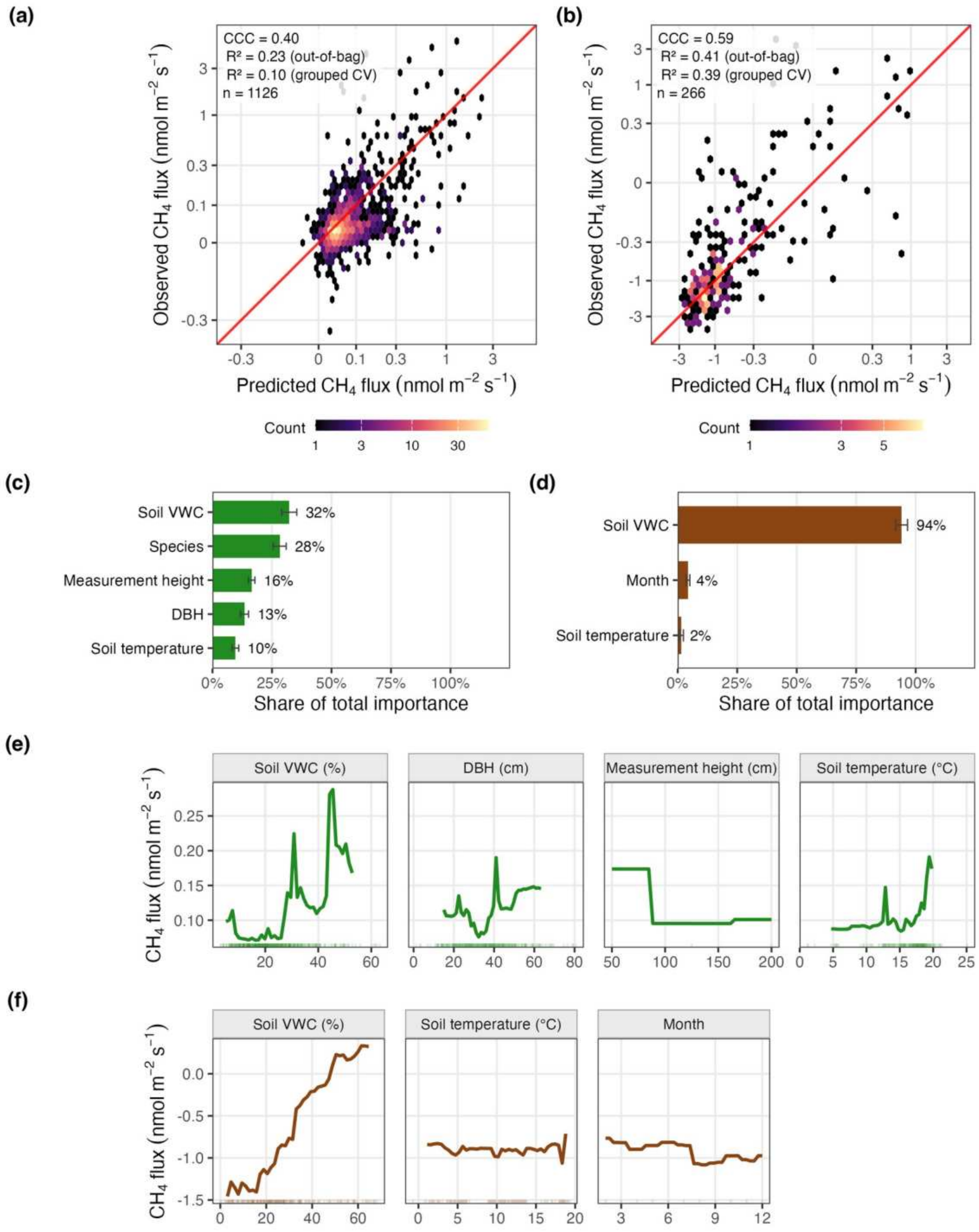


**Figure S25. Random-forest models of stem and soil flux.** (a, b) Observed against out-of-bag predicted flux for stems (a) and soils (b), on arcsinh axes, in hexagonal bins shaded by count (log scale); red line, 1:1. Each panel gives the concordance correlation coefficient and $R^2$ of the out-of-bag predictions, the grouped-cross-validation $R^2$ (whole trees or collars held out), and n. (c, d) Permutation importance on held-out data as a share of the total: in each of 50 random 70/30 splits of the measurements, a forest with the fitted model's mtry and node size, with 800 trees, was refitted on

70% and each predictor permuted 20 times in the held-out 30%; bars, mean over the splits; error bars, ±1 SE of that mean. (e, f) Partial dependence for each predictor other than species, over its 2nd–98th percentile; rugs, observed values.

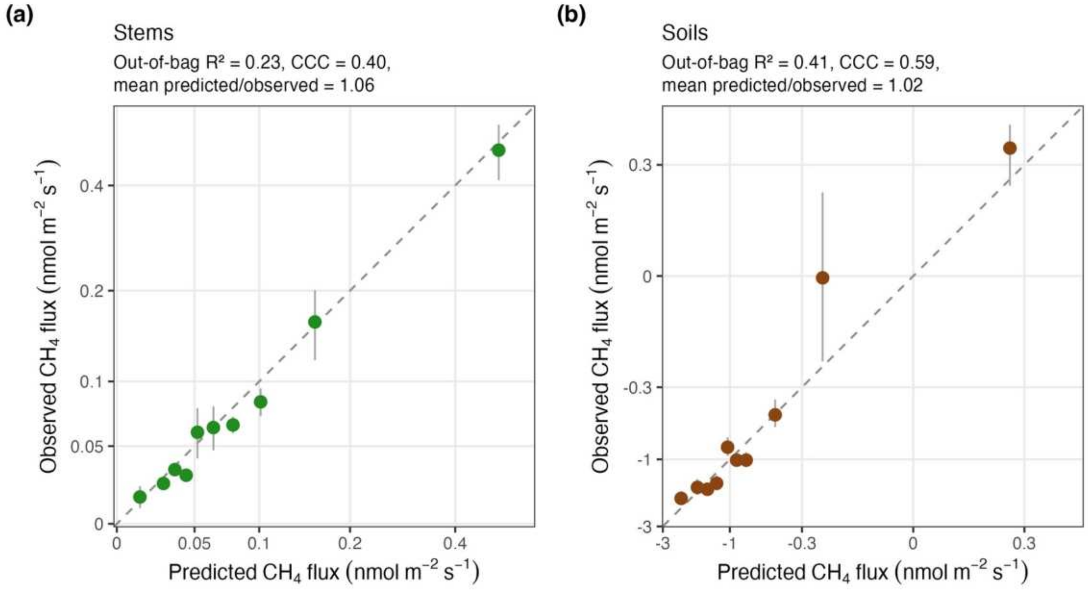


**Figure S26. Calibration of random-forest predictions.** Mean observed against mean out-of-bag predicted flux in bins of predicted flux, for stems (a) and soils (b); bars, ±1 SE; dashed line, 1:1. Out-of-bag $R^2$, concordance and mean predicted/observed annotated.

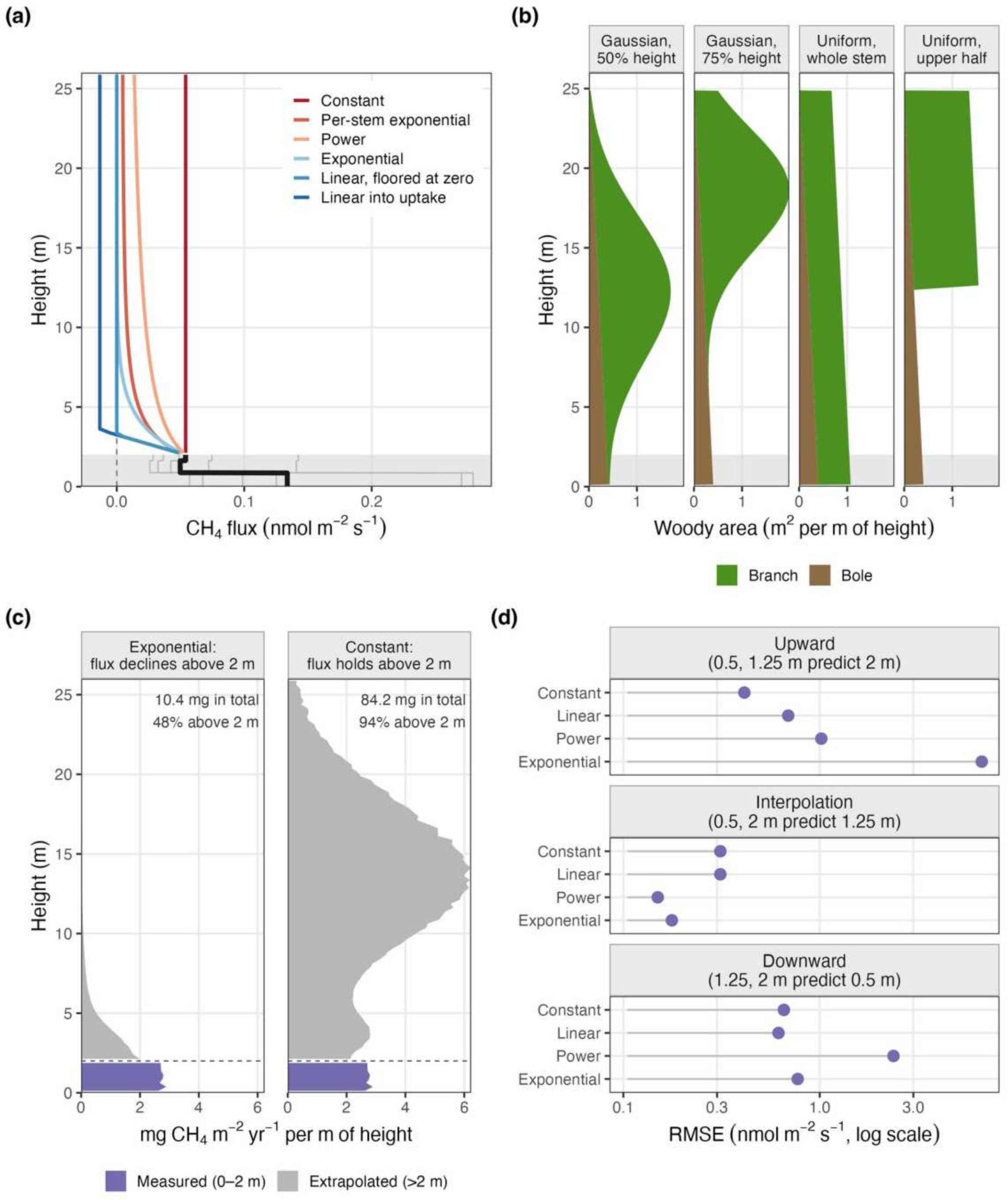


**Figure S27. The scaling assumptions.** (a) Stem $CH_4$ flux with height: the modeled 0–2 m profile (black, area-weighted over all stems; gray, the six commonest species; shaded, the measured band) and the six forms of flux above 2 m. (b) Bole (brown) and branch (green) woody area with height for a 25 m canopy tree under each branch placement, with a conic bole; a paraboloid bole gives nearly identical profiles. (c) Emission per m of height summed over the inventory for the two cases quoted in the text, exponential decay fitted to the measured heights and constant flux, at woody area index 2.93 with branch area centered at 75% of height; purple, measured band; gray, extrapolated. Totals are mg $CH_4$ $m^{-2}$ ground $yr^{-1}$. (d) Leave-one-height-out cross-validation: RMSE of each form when fitted to two of the three measured heights and used to predict the third (Methods S6).

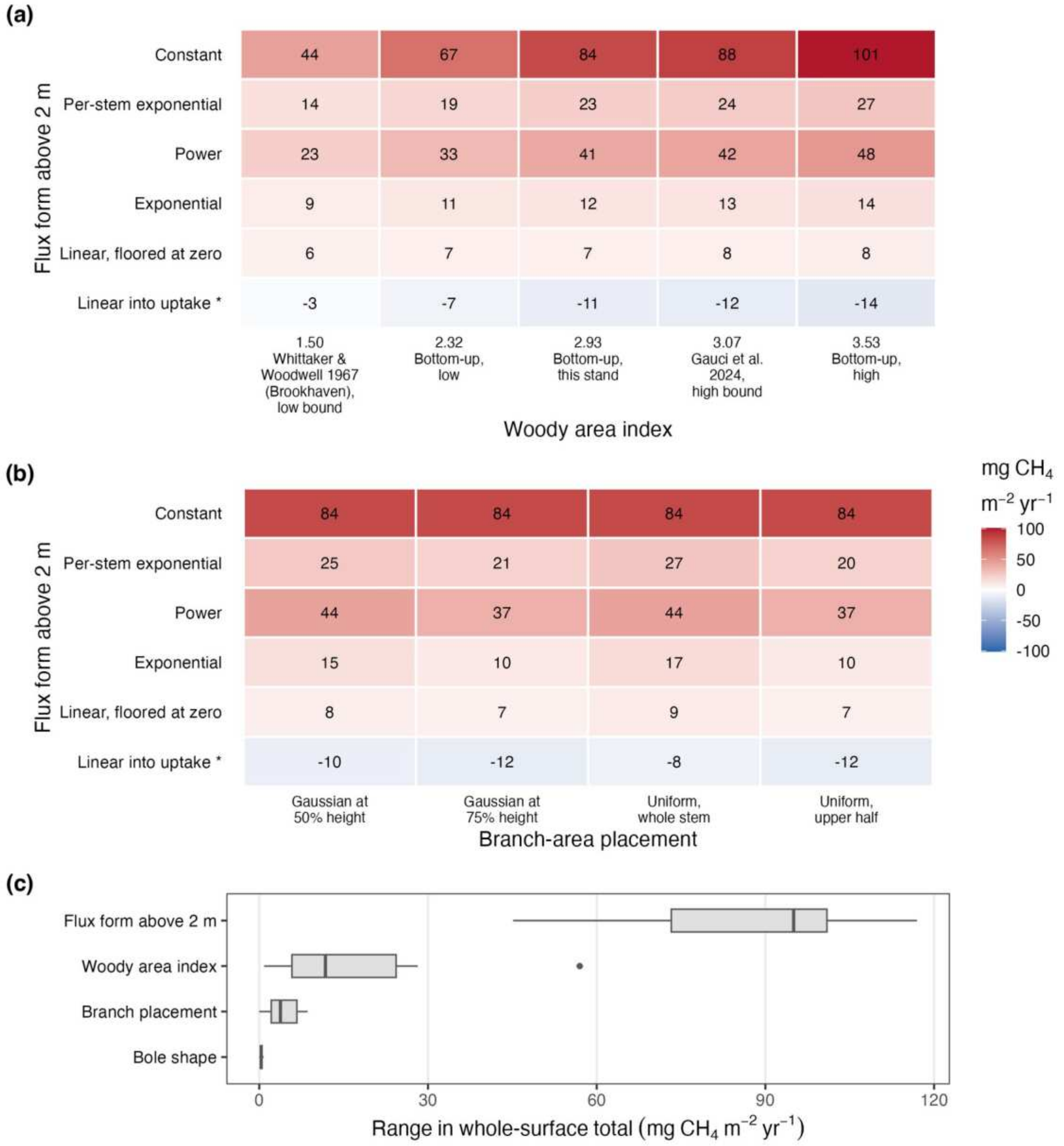


**Figure S28. Whole-woody-surface stem emission across the scaling grid.** (a) Median total for each flux form above 2 m and woody area index, taken over bole shapes and branch placements. (b) Median total for each flux form and branch placement, taken over woody area index and bole shape. Values are mg $CH_4$ $m^{-2}$ ground $yr^{-1}$ and include the measured 0–2 m band (5.47) in every cell; blue, net uptake; *, the linear-into-uptake form (Methods S2, S6). (c) Leverage of each assumption: for every combination of the other assumptions, the range in the total spanned by varying this one; boxes show the distribution of those ranges. The full grid of 240 combinations is archived (Data Availability).

### Supporting Information: Tables S1–S8

#### Methane emissions from upland tree stems reflect a balance of microbial production and consumption

Tables S1, S3–S5 and S7 are below. Tables S2 and S6 are long reference tables and are supplied as a separate Excel workbook (Tables S2 and S6), with each legend on its sheet; the legends are repeated here. The full datasets behind all tables are archived on Zenodo (Data Availability).

**Table S1.** ddPCR assays: primers, probe and thermocycling. Amplicon lengths for *pmoA* and *mmoX* are predicted *in silico* from the primer binding sites in *Methylococcus capsulatus* Bath (GenBank L40804 and M90050).

| Target | Primers and probe (5′→3′) | Chemistry | Thermocycling | Source |
|---|---|---|---|---|
| *mcrA* | F ACGACYTRCAGGAYCAGTGY; probe WGGWCCWAACTAYCCBAACTACG; R TGGTGWCCBACGTTCATYG (118 bp) | Probe (FAM) | 95 °C 10 min; 40 × [94 °C 30 s, 48 °C 80 s]; 98 °C 10 min; ramp 2 °C $s^{-1}$ | Arnold *et al.* (2024) |
| *pmoA* | 189f GGNGACTGGGACTTCTGG; mb661r CCGGMGCAACGTCYTTACC (508 bp) | EvaGreen | 95 °C 5 min; 40 × [95 °C 30 s, 52 °C 35 s, 72 °C 50 s]; 4 °C 5 min; 90 °C 5 min; ramp 1 °C $s^{-1}$ | Bourne *et al.* (2001) |
| *mmoX* | 536f CGCTGTGGAAGGGCATGAAGCG; 898r GCTCGACCTTGAACTTGGAGCC (363 bp) | EvaGreen | 95 °C 5 min; 40 × [95 °C 30 s, 52 °C 30 s, 72 °C 45 s]; 4 °C 5 min; 90 °C 5 min; ramp 1.5 °C $s^{-1}$ | Fuse *et al.* (1998) |

**Table S2. Methane-cycling taxa in the 16S data and their classification** (Excel workbook, sheet “Table S2”). Methanotrophs are classified by methane monooxygenase capacity. Known: genera that carry particulate or soluble methane monooxygenase, including the NC10 genus *Candidatus* Methylomirabilis. Putative: an upper bound: members of methanotroph-containing families unresolved at genus level, and genera in those families whose capacity is untested (no sequenced genome) or found in only some genomes (*Methylovirgula*), or represented by a single candidate homolog of unresolved source (*Rhodoblastus*; Table S3). Genera within those families with no annotated methane monooxygenase among their NCBI protein records are given in Table S3 and are not counted. For each taxon the sheet gives the number of amplicon sequence variants (ASVs) and its share of all non-organelle reads, pooled over the 575 study sample libraries. Classification follows Knief (2015), with genome or enrichment evidence for *Candidatus* Methylomirabilis (Ettwig *et al.*, 2010) and *Methylovirgula* (Gwak *et al.*, 2022). Methane monooxygenase in Methylacidiphilaceae is confirmed only in acidophilic genera (Op den Camp *et al.*, 2009; van Teeseling *et al.*, 2014); mesophilic relatives from peat (Dedysh *et al.*, 2021) and tree bark (Leung *et al.*, 2026) lack it, and members of the family are reported from tree bark (Jeffrey *et al.*, 2021). *Methylocella* is a facultative methanotroph that lacks pMMO (Dedysh *et al.*, 2005). The source for each taxon is given in the workbook. ASV counts are from the unrarefied 16S table of the 575 study sample libraries, and abundance is each taxon’s share of all non-organelle 16S reads pooled across those libraries; Methods S3 instead counts ASVs present in survey samples after rarefaction to 3,500 reads, so its totals are smaller.

**Table S3. Genome screen for methane monooxygenase in genera within mixed methanotroph families.** For each genus, NCBI genome assemblies and protein records named as methane monooxygenase subunits (queried 30 September 2026). Up to 60 named records per genus were aligned to PmoA of *Methylococcus capsulatus* and MmoX of *Methylosinus trichosporium* (best identity among alignments of at least 50 amino acids); a record counts as a true subunit at ≥40% identity over ≥150 amino acids to PmoA or ≥55% identity over ≥300 amino acids to MmoX, because protein names alone are unreliable. The capacity verdict follows from checking the records that pass: in one genome only, in some strains only (strain-specific), or untested where no genome is sequenced. The Table S2 tier is the genus’s classification there; a genus with a passing record is counted (Known or Putative), one whose records all fail is not, and one with no sequenced genome is Putative. *Methylocapsa* and *Methylocella* are positive controls. The one *Rhodoblastus* protein passing the PmoA threshold comes from a source genome that could not be confirmed, so it is treated as a candidate copper monooxygenase of uncertain function, not a verified MMO, and the genus is Putative. *Lichenibacterium* is the genus recorded as 1174-901-12 in SILVA. Beijerinckiaceae include genera that have lost methanotrophy

(Tamas *et al.*, 2014). *Methylobacterium* is screened because growth on methane was reported for *M. populi* (Van Aken *et al.*, 2004) but has not been confirmed (Dedysh *et al.*, 2004; Semrau *et al.*, 2011); no bacterial genome assembled from lichen metagenomes carried methane monooxygenase (Tagirdzhanova *et al.*, 2024).

| Genus | NCBI assemblies | Named MMO protein records | Best identity to PmoA (%) | Best identity to MmoX (%) | Record passing the subunit threshold | Capacity verdict | Table S2 tier |
|---|---|---|---|---|---|---|---|
| *Methylocapsa* | 15 | 56 | 69.9 | 97.3 | yes | positive control | Known |
| *Methylocella* | 88 | 228 | 28.4 | 96.9 | yes | positive control | Known |
| *Roseiarcus* | 85 | 0 | 0 | 0 | no | no | not counted |
| *Bosea* | 245 | 0 | 0 | 0 | no | no | not counted |
| *Microvirga* | 179 | 6 | 24.1 | 25.9 | no | no | not counted |
| *Methylovirgula* | 68 | 10 | 22.4 | 84.9 | yes | yes (strain-specific) | Putative |
| *Methylorosula* | 0 | 0 | 0 | 0 | untested (no genome) | untested (no genome) | Putative |
| *Rhodoblastus* | 72 | 3 | 63.2 | 20.8 | yes | yes (one genome) | Putative |
| *Psychroglaciecola* | 0 | 0 | 0 | 0 | untested (no genome) | untested (no genome) | Putative |
| *Lichenibacteri* | 4 | 0 | 0 | 0 | no | no | not counte |

| Genus | NCBI assemblies | Named MMO protein records | Best identity to PmoA (%) | Best identity to MmoX (%) | Record passing the subunit threshold | Capacity verdict | Table S2 tier |
|---|---|---|---|---|---|---|---|
| *um* | | | | | | | d |
| *Methylobacterium* | 583 | 36 | 32.7 | 32.7 | no | no | not counted |
| *Methylorubrum* | 123 | 2 | 25.7 | 28.7 | no | no | not counted |

**Table S4. Methanotrophs in the 16S data at several taxonomic resolutions, their agreement with ddPCR, and the felled oak by tissue.** (a) Survey: mean share of each sample's community (%; chloroplast and mitochondrial reads excluded as in the main analysis; libraries of at least 1,000 reads, not rarefied, so values differ from Fig. 5 and the text, which use the rarefied data), with Spearman ρ against ddPCR *pmoA* / *mmoX* copies per reaction in the same extracts. Known, genera carrying methane monooxygenase; Putative, genus-unresolved members of methanotroph-containing families and genera of untested or partial capacity (an upper bound); tree-placed Putative, the genus-unresolved Beijerinckiaceae that the ASV tree places within 0.03 substitutions per site of a methanotroph genus (Methods S3). (b) Known + tree-placed Putative by placement threshold. (c) Felled black oak: mean share of the community by tissue (%), with the number of libraries in which the group was detected; one tree, libraries of at least 1,000 reads after the same exclusion; the oak data are not on the ASV tree.

| Grouping | Heartwood | Sapwood | Mineral soil | Organic soil |
|---|---|---|---|---|
| Methylococcales (order) | 0.000 (– / –) | 0.000 (– / –) | 0.009 (0.10 / 0.16) | 0.008 (0.07 / 0.15) |
| Methylacidiphilales (order) | 0.094 (0.04 / −0.13) | 0.099 (0.26 / −0.11) | 0.258 (−0.21 / −0.18) | 1.133 (−0.39 / −0.51) |
| Methylomirabilales, NC10 (order) | 0.000 (– / –) | 0.000 (– / –) | 0.029 (−0.07 / 0.05) | 0.012 (0.02 / 0.18) |
| Beijerinckiaceae, all (family) | 1.586 (−0.01 / 0.04) | 4.338 (0.29 / 0.03) | 3.051 (−0.41 / −0.42) | 1.861 (−0.36 / −0.43) |
| Known | 0.016 (−0.17 | 0.031 (0.11 / | 0.975 (−0.35 | 0.061 (0.06 / |

| Grouping | Heartwood | Sapwood | Mineral soil | Organic soil |
|---|---|---|---|---|
| | / −0.02) | 0.10) | / −0.43) | −0.01) |
| Known + tree-placed Putative | 0.091 (−0.03 / −0.09) | 0.148 (0.06 / −0.02) | 1.124 (−0.31 / −0.40) | 0.079 (0.04 / −0.04) |
| Known + Putative | 0.556 (0.02 / −0.01) | 1.316 (0.22 / 0.01) | 1.406 (−0.34 / −0.42) | 1.260 (−0.36 / −0.49) |

| Placement threshold (substitutions per site) | Placed ASVs | Heartwood | Sapwood | Mineral soil | Organic soil |
|---|---|---|---|---|---|
| 0.02 | 59 | 0.087 | 0.114 | 1.119 | 0.076 |
| 0.03 | 74 | 0.091 | 0.148 | 1.124 | 0.079 |
| 0.04 | 91 | 0.109 | 0.205 | 1.126 | 0.079 |

| Grouping | Bark | Branch | Foliage | Sapwood | Heartwood | Fine root | Coarse root | Organic soil | Mineral soil |
|---|---|---|---|---|---|---|---|---|---|
| Known | 0.50 (2/4) | 0.60 (3/4) | 0.62 (2/4) | 0.00 (0/7) | 0.02 (1/14) | 0.00 (0/3) | 0.00 (0/5) | 0.31 (2/4) | 0.00 (0/4) |
| *Methylocella* | 0.45 (1/4) | 0.60 (3/4) | 0.62 (2/4) | 0.00 (0/7) | 0.02 (1/14) | 0.00 (0/3) | 0.00 (0/5) | 0.00 (0/4) | 0.00 (0/4) |
| *Methylocapsa* | 0.06 (1/4) | 0.00 (0/4) | 0.00 (0/4) | 0.00 (0/7) | 0.00 (0/14) | 0.00 (0/3) | 0.00 (0/5) | 0.31 (2/4) | 0.00 (0/4) |
| Known + Putative | 1.69 (4/4) | 2.22 (4/4) | 2.52 (3/4) | 1.72 (5/7) | 2.52 (14/14) | 0.75 (3/3) | 0.27 (2/5) | 0.47 (3/4) | 0.73 (3/4) |
| Beijerinckiaceae, all | 6.64 (4/4) | 21.97 (4/4) | 45.19 (4/4) | 3.35 (6/7) | 4.24 (14/14) | 1.14 (3/3) | 1.06 (3/5) | 1.44 (4/4) | 1.25 (3/4) |
| Methylococcales | 0.00 (0/4) | 0.00 (0/4) | 0.00 (0/4) | 0.00 (0/7) | 0.00 (0/14) | 0.00 (0/3) | 0.00 (0/5) | 0.00 (0/4) | 0.00 (0/4) |
| Methylacidiphil | 0.05 (1/4) | 0.00 (0/4) | 0.00 (0/4) | 0.00 (0/7) | 0.00 (1/14) | 0.75 (3/3) | 0.00 (0/5) | 0.16 (3/4) | 0.73 (3/4) |

| Grouping | Heartwood | Sapwood | Mineral soil | Organic soil |
|---|---|---|---|---|
| ales | | | | |

**Table S5. Agreement between ddPCR and 16S detection in the same samples, by compartment.** Each gene is compared with its 16S group: methanogens for *mcrA*, and Known plus Putative methanotrophs for *pmoA* and *mmoX*. Samples are counted by whether each method detected the group; detection rates are the percentage of samples in which each method detected it. r and p, Pearson correlation of $\log_{10}$ abundances in the samples where both methods detect (computed where there are at least four).

| Gene | Compartment | Samples | Both detect | ddPCR only | 16S only | Neither | Detected by ddPCR (%) | Detected by 16S (%) | r | p |
|---|---|---|---|---|---|---|---|---|---|---|
| *mcrA* | Soil | 254 | 15 | 126 | 1 | 112 | 56 | 6 | 0.86 | <0.001 |
| *mcrA* | Wood | 257 | 57 | 158 | 1 | 41 | 84 | 23 | 0.81 | <0.001 |
| *mmoX* | Soil | 255 | 219 | 36 | 0 | 0 | 100 | 86 | −0.59 | <0.001 |
| *mmoX* | Wood | 256 | 142 | 100 | 9 | 5 | 95 | 59 | 0.03 | 0.699 |
| *pmoA* | Soil | 256 | 220 | 36 | 0 | 0 | 100 | 86 | −0.26 | <0.001 |
| *pmoA* | Wood | 257 | 148 | 101 | 4 | 4 | 97 | 59 | 0.16 | 0.053 |

**Table S6. Diameter at breast height of measured trees by campaign, location, species and status** (Excel workbook, sheet “Table S6”). Number of trees and DBH (cm; mean ± SD, or a single value where one tree was measured).

**Table S7. Methanotroph identifications reported from trees and plant tissues, and the evidence behind them.** MMO, methane monooxygenase.

| Study | Plant and tissue | Method | Reported as methanotrophs | Evidence for MMO | Status |
|---|---|---|---|---|---|
| Van Aken *et al.* | *Populus*, endophyte | Isolate | *Methylobacterium* | Growth on methane | Not confirmed |

| Study | Plant and tissue | Method | Reported as methanotrophs | Evidence for MMO | Status |
|---|---|---|---|---|---|
| (2004) | | | *populi* | reported | (Dedysh *et al.*, 2004; Semrau *et al.*, 2011); no annotated MMO among NCBI protein records (Table S3) |
| Doronina *et al.* (2004) | Woody tissues, including *Picea pungens* needles | Isolates | *Methylocystis*-related strain | Methanotroph genus; isolate also grew on methanol | Facultative; oxidation in planta not tested (Putkinen *et al.*, 2021) |
| Iguchi *et al.* (2012) | Phyllosphere, including Norway spruce and *Pinus parviflora* needles | Isolation under 20% $CH_4$ | *Methylomonas*, *Methylocystis* | Methanotroph genera; isolates also grew on methanol | Oxidation capacity on leaves not tested (Putkinen *et al.*, 2021) |
| Putkinen *et al.* (2021) | Pine and spruce shoots | Metagenomes (database search and capture) | Few canonical methanotrophs | No canonical *pmoA*; two alphaproteobacterial *mmoX* fragments; monooxygenases related to propane and butane | Canonical methanotrophs scarce; $CH_4$ oxidation by the other monooxygenases unknown |

| Study | Plant and tissue | Method | Reported as methanotrophs | Evidence for MMO | Status |
|---|---|---|---|---|---|
| | | | | monooxygenases | |
| Jeffrey *et al.* (2021) | *Melaleuca quinquenervia* bark (wetland) | 16S and *pmoA* amplicons, qPCR, inhibition | *Methylomonas* (dominant); novel Methylacidiphilaceae | *pmoA* for *Methylomonas*; 16S family membership for Methylacidiphilaceae | *Methylomonas* secure; related Methylacidiphilales genomes from the same bark lack MMO (Leung *et al.*, 2026) |
| Leung *et al.* (2026) | Bark of eight Australian tree species, wetland to upland | Metagenomes, genomes, microcosms | *Methylomonas* (dominant in wetland bark); *Methylocystis*; *Methylocella*, including in upland *Eucalyptus* bark | Genomes and contigs with *pmoA* and *mmoX* | Secure; two Methylacidiphilales genomes lack MMO |
| Moisan *et al.* (2025); Moisan *et al.* (2026) | Floodplain trees: leaves, bark, wood | 16S, with facultative and obligate methanotrophs selected | *Methylobacterium*–*Methylorubrum* dominant in leaves and bark (2026) | None; the genus lacks MMO | Methane-oxidizer signal largely methylotrophs |
| This study | Upland trees: heartwood and sapwood; | 16S (genus rule; tree placement as a | *Methylocapsa* (mineral soil); *Methyloce* | Genus carries MMO; ddPCR genes | Known methanotrophs 0.02–0.03% of wood |

| Study | Plant and tissue | Method | Reported as methanotrophs | Evidence for MMO | Status |
| --- | --- | --- | --- | --- | --- |
| | felled oak tissues; soils | check); ddPCR *pmoA*, *mmoX* | *lla* (oak bark, branches, foliage); NC10 traces | | communities, about 0.1% with tree-placed Putative ASVs; concentrated in the oak's outer and aerial tissues |

**Table S8. Sources of the species trait values (Methods S5).** Values for the 13 traits analyzed; TRY dataset IDs in parentheses. All TRY data are cited as Kattge *et al.* (2020). The source of every species value, with the TRY record IDs, is archived with the data (Data Availability). GWDD, Global Wood Density Database.

| Trait | Source | Dataset (TRY ID) | Reference |
| --- | --- | --- | --- |
| Wood density; bark density | GWDD v2.2 | — | Fischer *et al.* (2026) |
| Stem carbon; bark carbon | TRY | Wood carbon database (433) | Martin *et al.* (2018) |
| Wood C:N; bark volume relative to wood | TRY | Thom 2019 (662) | D. Thom, unpublished |
| Coarse-wood decomposition rate | TRY | Functional Ecology of Trees database (68) | Wirth & Lichstein (2009) |
| Waterlogging tolerance | TRY | Tree Tolerance Database (49) | Niinemets & Valladares (2006) |
| Fine-root diameter; fine-root tissue density | TRY | Fine-Root Ecology Database (339); fine-root traits across seed plants (759) | Iversen *et al.* (2017); Valverde-Barrantes *et al.* (2020) |

| Trait | Source | Dataset (TRY ID) | Reference |
|---|---|---|---|
| Mycorrhizal type | TRY | Fine-Root Ecology Database (339); Global $^{15}N$ Database (130); Mycorrhiza Database (189); Mycorrhizal Intensity Database (162); fine-root traits across seed plants (759) | Iversen *et al.* (2017); Craine *et al.* (2009); Koele *et al.* (2012); Akhmetzhanova *et al.* (2012); Valverde-Barrantes *et al.* (2020) |
| Maximum longevity | TRY | Global Seed Mass, Plant Height Database (28); Functional Ecology of Trees database (68) | Moles *et al.* (2004); Wirth & Lichstein (2009) |
| Soil-moisture niche | This study | — | Fig. S6 |


**Akhmetzhanova AA, Soudzilovskaia NA, Onipchenko VG, Cornwell WK, Agafonov VA, Selivanov IA, Cornelissen JHC**. **2012**. A rediscovered treasure: Mycorrhizal intensity database for 3000 vascular plant species across the former soviet union: Ecological archives E093-059. *Ecology* **93**: 689–690.

**Arnold W, Gewirtzman J, Raymond PA, Bradford MA, Butler C, Peccia J**. **2024**. A method for sampling the living wood microbiome. *Methods in Ecology and Evolution* **15**: 1084–1096.

**Bourne DG, McDonald IR, Murrell JC**. **2001**. Comparison of *pmoA* PCR primer sets as tools for investigating methanotroph diversity in three Danish soils. *Applied and Environmental Microbiology* **67**: 3802–3809.

**Craine JM, Elmore AJ, Aidar MPM, Bustamante M, Dawson TE, Hobbie EA, Kahmen A, Mack MC, McLauchlan KK, Michelsen A, *et al.*** **2009**. Global patterns of foliar nitrogen isotopes and their relationships with climate, mycorrhizal fungi, foliar nutrient concentrations, and nitrogen availability. *New Phytologist* **183**: 980–992.

**Dedysh SN, Beletsky AV, Ivanova AA, Danilova OV, Begmatov S, Kulichevskaya IS, Mardanov AV, Ravin NV**. **2021**. Peat-inhabiting Verrucomicrobia of the order Methylacidiphilales do not possess methanotrophic capabilities. *Microorganisms* **9**: 2566.

**Dedysh SN, Dunfield PF, Trotsenko YA**. **2004**. Methane utilization by *Methylobacterium* species: New evidence but still no proof for an old controversy. *International Journal of Systematic and Evolutionary Microbiology* **54**: 1919–1920.

**Dedysh SN, Knief C, Dunfield PF**. **2005**. *Methylocella* species are facultatively methanotrophic. *Journal of Bacteriology* **187**: 4665–4670.

**Doronina NV, Ivanova EG, Suzina NE, Trotsenko YuA**. **2004**. Methanotrophs and methylobacteria are found in woody plant tissues within the winter period. *Microbiology* **73**: 702–709.

**Ettwig KF, Butler MK, Le Paslier D, Pelletier E, Mangenot S, Kuypers MMM, Schreiber F, Dutilh BE, Zedelius J, de Beer D, *et al.* 2010**. Nitrite-driven anaerobic methane oxidation by oxygenic bacteria. *Nature* **464**: 543–548.

**Fischer FJ, Chave J, Zanne A, Jucker T, Fajardo A, Fayolle A, Lima RAF de, Vieilledent G, Beeckman H, Hubau W, *et al.* 2026**. Beyond species means – the intraspecific contribution to global wood density variation. *New Phytologist* **249**: 2630–2651.

**Fuse H, Ohta M, Takimura O, Murakami K, Inoue H, Yamaoka Y, Oclarit JM, Omori T**. **1998**. Oxidation of trichloroethylene and dimethyl sulfide by a marine *Methylomicrobium* strain containing soluble methane monooxygenase. *Bioscience, Biotechnology, and Biochemistry* **62**: 1925–1931.

**Gwak J-H, Awala SI, Nguyen N-L, Yu W-J, Yang H-Y, von Bergen M, Jehmlich N, Kits KD, Loy A, Dunfield PF, *et al.* 2022**. Sulfur and methane oxidation by a single microorganism. *Proceedings of the National Academy of Sciences, USA* **119**: e2114799119.

**Iguchi H, Sato I, Sakakibara M, Yurimoto H, Sakai Y**. **2012**. Distribution of methanotrophs in the phyllosphere. *Bioscience, Biotechnology, and Biochemistry* **76**: 1580–1583.

**Iversen CM, McCormack ML, Powell AS, Blackwood CB, Freschet GT, Kattge J, Roumet C, Stover DB, Soudzilovskaia NA, Valverde-Barrantes OJ, *et al.* 2017**. A global fine-root ecology database to address below-ground challenges in plant ecology. *New Phytologist* **215**: 15–26.

**Jeffrey LC, Maher DT, Chiri E, Leung PM, Nauer PA, Arndt SK, Tait DR, Greening C, Johnston SG**. **2021**. Bark-dwelling methanotrophic bacteria decrease methane emissions from trees. *Nature Communications* **12**: 2127.

**Kattge J, Bönisch G, Díaz S, Lavorel S, Prentice IC, Leadley P, Tautenhahn S, Werner GDA, Aakala T, Abedi M, *et al.* 2020**. TRY plant trait database – enhanced coverage and open access. *Global Change Biology* **26**: 119–188.

**Knief C. 2015**. Diversity and habitat preferences of cultivated and uncultivated aerobic methanotrophic bacteria evaluated based on *pmoA* as molecular marker. *Frontiers in Microbiology* **6**: 1346.

**Koele N, Dickie IA, Oleksyn J, Richardson SJ, Reich PB. 2012**. No globally consistent effect of ectomycorrhizal status on foliar traits. *New Phytologist* **196**: 845–852.

**Leung PM, Jeffrey LC, Bay SK, Gomez-Alvarez P, Hall M, Johnston SG, Dittmann J, Deschaseaux E, Hopkins B, Haskell J, *et al.* 2026**. Bark microbiota modulate climate-active gas fluxes in Australian forests. *Science* **391**: eadu2182.

**Martin AR, Doraisami M, Thomas SC. 2018**. Global patterns in wood carbon concentration across the world's trees and forests. *Nature Geoscience* **11**: 915–920.

**Moisan M-A, Maire V, Isabelle J, Philippo D, Martineau C. 2025**. Tissue humidity and pH as important species traits regulating tree methane emissions in floodplain wetland forests. *New Phytologist* **248**: 1713–1727.

**Moisan M-A, Maire V, Morency M-J, Martineau C. 2026**. Tree tissues and species traits modulate the microbial methane-cycling communities of the tree phyllosphere. *Scientific Reports* **16**: 19771.

**Moles AT, Falster DS, Leishman MR, Westoby M. 2004**. Small-seeded species produce more seeds per square metre of canopy per year, but not per individual per lifetime. *Journal of Ecology* **92**: 384–396.

**Niinemets Ü, Valladares F. 2006**. Tolerance to shade, drought, and waterlogging of temperate northern hemisphere trees and shrubs. *Ecological Monographs* **76**: 521–547.

**Op den Camp HJM, Islam T, Stott MB, Harhangi HR, Hynes A, Schouten S, Jetten MSM, Birkeland N, Pol A, Dunfield PF. 2009**. Environmental, genomic and taxonomic perspectives on methanotrophic Verrucomicrobia. *Environmental Microbiology Reports* **1**: 293–306.

**Putkinen A, Siljanen HMP, Laihonen A, Paasisalo I, Porkka K, Tiirola M, Haikarainen I, Tenhovirta S, Pihlatie M. 2021**. New insight to the role of microbes in the methane exchange in trees: Evidence from metagenomic sequencing. *New Phytologist* **231**: 524–536.

**Semrau JD, DiSpirito AA, Vuilleumier S. 2011**. Facultative methanotrophy: False leads, true results, and suggestions for future research. *FEMS Microbiology Letters* **323**: 1–12.

**Tagirdzhanova G, Saary P, Cameron ES, Allen CCG, Garber AI, Escandón DD, Cook AT, Goyette S, Nogerius VT, Passo A, *et al.* 2024**. Microbial occurrence and symbiont detection in a global sample of lichen metagenomes. *PLoS Biology* **22**: e3002862.

**Tamas I, Smirnova AV, He Z, Dunfield PF. 2014**. The (d)evolution of methanotrophy in the Beijerinckiaceae—a comparative genomics analysis. *The ISME Journal* **8**: 369–382.

**Valverde-Barrantes OJ, Maherali H, Baraloto C, Blackwood CB. 2020**. Independent evolutionary changes in fine-root traits among main clades during the diversification of seed plants. *New Phytologist* **228**: 541–553.

**Van Aken B, Peres CM, Doty SL, Yoon JM, Schnoor JL. 2004**. *Methylobacterium populi* sp. nov., a novel aerobic, pink-pigmented, facultatively methylotrophic, methane-utilizing bacterium isolated from poplar trees (*Populus deltoides×nigra* DN34). *International Journal of Systematic and Evolutionary Microbiology* **54**: 1191–1196.

**van Teeseling MCF, Pol A, Harhangi HR, van der Zwart S, Jetten MSM, Op den Camp HJM, van Niftrik L. 2014**. Expanding the verrucomicrobial methanotrophic world: description of three novel species of *Methylacidimicrobium* gen. nov. *Applied and Environmental Microbiology* **80**: 6782–6791.

**Wirth C, Lichstein JW. 2009**. The imprint of species turnover on old-growth forest carbon balances - insights from a trait-based model of forest dynamics. In: Old-growth forests. Springer Berlin Heidelberg, 81–113.
