## Supplementary material for "Methane emissions from upland tree stems reflect a balance of microbial production and consumption": Notes S1: ddPCR protocol for methane-cycling genes

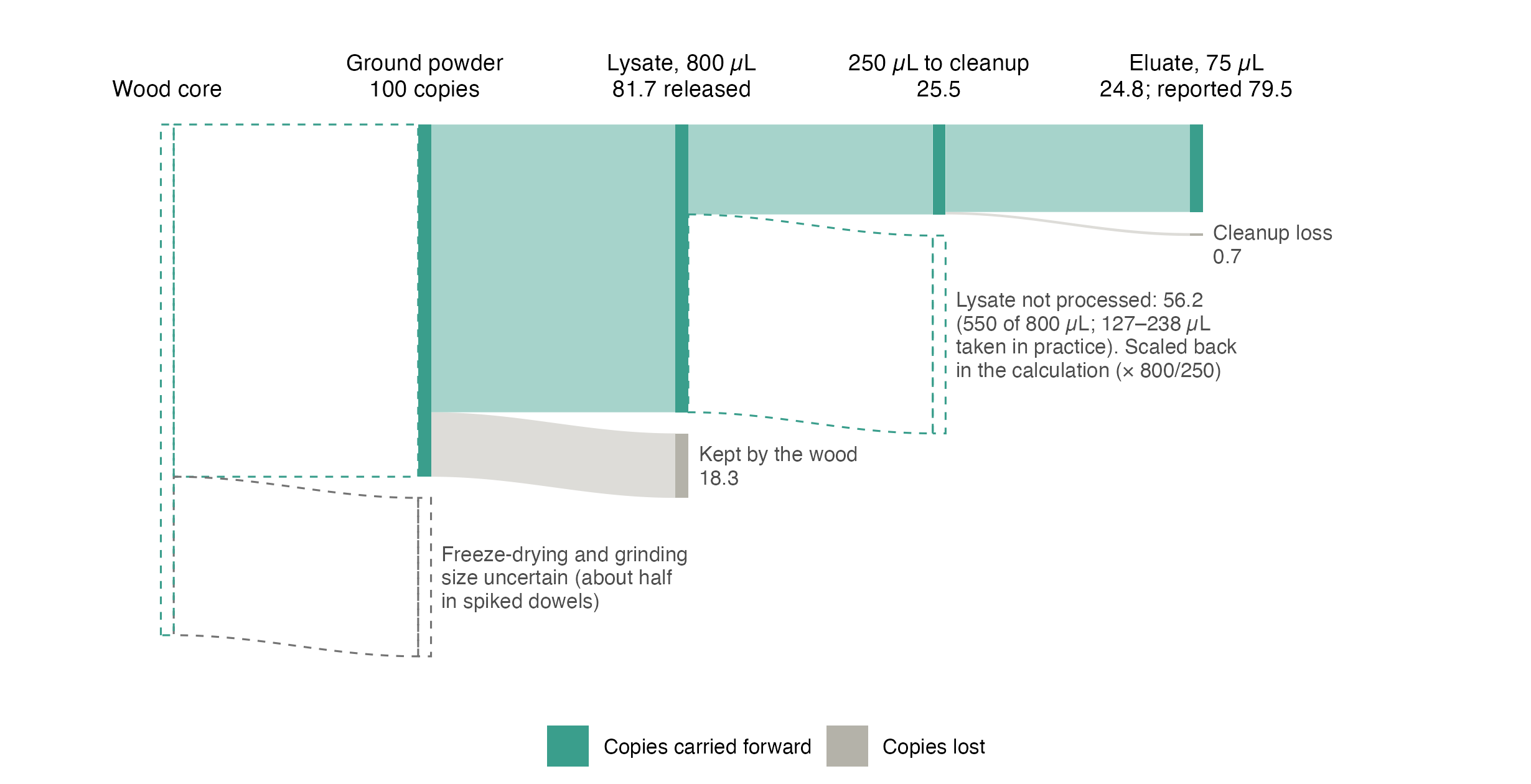

*Where target copies are lost between the core and the reaction.* Freeze-drying and cryo-grinding lose copies before extraction (dashed: about half in spiked sterile dowels, a different medium from field cores, so the size of this loss in cores is uncertain). Of 100 copies in the ground powder, the wood releases 81.7 (the mean field-wood recovery relative to the lysis-buffer control reported by Arnold *et al.* (2024)), 250 of the 800 µL of lysate is processed, and the cleanup keeps 97% (the lysis-buffer control’s 30.4% divided by the 31% processed), so about 25 reach the eluate. The calculation scales the eluate back to the whole lysate (× 800/250), so about 79 are reported; the lysate not processed (dashed) is not a loss in reported values. The volume actually drawn off above the pellet varies (127–238 µL in the method study’s records), so reported values can differ about two-fold between otherwise identical samples. Transferring 2.5 µL of eluate into each reaction is a subsample that the calculation scales back, not a loss. Values from Arnold *et al.* (2024).

**Record for every plate**

- sample masses;
- elution, template and reaction volumes;
- reagent lots;
- the thresholds, with an image of the amplitude plot;
- accepted droplets per well;
- standards, no-template controls and blanks, labeled as such in the export.

**Arnold W, Gewirtzman J, Raymond PA, Bradford MA, Butler C, Peccia J**. **2024**. [A method for sampling the living wood microbiome](https://doi.org/10.1111/2041-210X.14311). *Methods in Ecology and Evolution* **15**: 1084–1096.

**Bourne DG, McDonald IR, Murrell JC**. **2001**. [Comparison of *pmoA* PCR primer sets as tools for investigating methanotroph diversity in three Danish soils](https://doi.org/10.1128/aem.67.9.3802-3809.2001). *Applied and Environmental Microbiology* **67**: 3802–3809.

**Forootan A, Sjöback R, Björkman J, Sjögreen B, Linz L, Kubista M**. **2017**. [Methods to determine limit of detection and limit of quantification in quantitative real-time PCR (qPCR)](https://doi.org/10.1016/j.bdq.2017.04.001). *Biomolecular Detection and Quantification* **12**: 1–6.

**Fuse H, Ohta M, Takimura O, Murakami K, Inoue H, Yamaoka Y, Oclarit JM, Omori T**. **1998**. [Oxidation of trichloroethylene and dimethyl sulfide by a marine *Methylomicrobium* strain containing soluble methane monooxygenase](https://doi.org/10.1271/bbb.62.1925). *Bioscience, Biotechnology, and Biochemistry* **62**: 1925–1931.

**Knief C**. **2015**. [Diversity and habitat preferences of cultivated and uncultivated aerobic methanotrophic bacteria evaluated based on *pmoA* as molecular marker](https://doi.org/10.3389/fmicb.2015.01346). *Frontiers in Microbiology* **6**: 1346.
